# A Mathematical Model of the Within-Host Kinetics of SARS-CoV-2 Neutralizing Antibodies Following COVID-19 Vaccination

**DOI:** 10.1101/2022.05.11.491557

**Authors:** Lisette de Pillis, Rebecca Caffrey, Ge Chen, Mark D. Dela, Leif Eldevik, Joseph McConnell, Shahrokh Shabahang, Stephen A. Varvel

## Abstract

Compelling evidence continues to build to support the idea that SARS-CoV-2 Neutralizing Antibody (NAb) levels in an individual can serve as an important indicator of the strength of protective immunity against infection. It is not well understood why NAb levels in some individuals remain high over time, while in others levels decline rapidly. In this work, we present a two-population mathematical model of within-host NAb dynamics in response to vaccination. By fitting only four host-specific parameters, the model is able to capture individual-specific NAb levels over time as measured by the AditxtScore^™^ for NAbs. The model can serve as a foundation for predicting NAb levels in the long-term, understanding connections between NAb levels, protective immunity, and break-through infections, and potentially guiding decisions about whether and when a booster vaccination may be warranted.

## 1. Introduction

SARS-CoV-2 Neutralizing Antibody (NAb) levels in an individual have been shown to be correlated to the strength of protective immunity against infection [1, 2, 3, 4]. Antibodies develop both in response to infection and to vaccination. Both non-neutralizing and neutralizing antibodies are involved in the immune response to viral infection, serving to alert effector cells to the presence of pathogen in infected cells as well as to disrupt the ability of a virus to enter a host cell. As pointed out in [5], neutralizing antibodies that develop under viral load pressures serve as sentinels that provide insight into the associated humoral response.

The focus of the model in our study is on understanding changes in individual-specific NAb concentrations in fully vaccinated individuals using a novel flow-cytometry-based NAb assay. NAb activity levels can be measured using two general approaches: Bioassays, which determine the ability of NAb to prevent viral infection of cells in culture media, and binding assays, which evaluate the ability of NAb to prevent binding of SARS-CoV-2 spike protein to the ACE-2 receptor on the surface of human cells. While the former is the gold standard for determining the effectiveness of antibodies to neutralize viral entry into cells, bioassays require more time and higher safety level laboratories and are, therefore, more expensive. Binding assays can serve as a more practical and scalable methodology to assess NAb levels. The novel flow-cytometry test that was used in this study was compared to a bioassay [6] to determine its performance using samples from 44 positive and negative samples in a blinded study. Results showed 100% concordance between the two methodologies in quantifying NAb levels in the samples (unpublished data).

Over time, NAb levels in some individuals remain high, while in others levels decline rapidly. Since NAb levels have the potential to be used as an indicator of the protective potential of the immune response to a viral challenge, our goal is to develop a mathematical model that can serve as a starting point for improving our ability to predict an individual’s NAb response to mRNA vaccine dosing, persistence of immune strength, and how quickly immunity levels may wane. In this work, we present a two-population mathematical model of within-host NAb dynamics in response to vaccination. By fitting only four subject-specific parameters, model simulations can capture NAb level changes within an individual measured by the AditxtScore^™^ for NAbs. The model can serve a foundation for predicting NAb levels over time and guiding decisions about whether and when a booster vaccination may be warranted.

### mRNA Vaccines

Both of the newly developed mRNA vaccines widely available in the U.S. function differently from traditional live-attenuated or disabled virus vaccines. The SARS-CoV-2 mRNA vaccine encodes for the spike protein, which harbors the receptor binding domain (RBD), to elicit the immune response. One of the effects of the immune response is production of neutralizing antibodies (NAb) that bind the RBD thereby preventing viral entry into the host cells. The correlation between how mRNA vaccines work and elicit production of NAb, which then interfere with binding of the spike protein to the cell receptor required for viral entry into the cell, indicates that evaluating NAb levels and NAb trajectories over time is important in understanding the likelihood of an individual having protective immunity against infection.

### Mathematical Models of Within-Host Responses

Since the outbreak of SARS-CoV-2 near the end of 2019, a number of mathematical models have been created to help guide medical care providers and health policy makers in establishing approaches to stemming the spread of disease and determining best practices for treatment and prevention. While many useful models focus on modeling epidemiological dynamics and answering questions about population-level effects of interventions such as vaccination and treatment (*c.f.* [7], [8]), our interest is in addressing questions about protective immunity within a vaccinated individual. Some excellent within-host mathematical models have been developed to focus on a variety of questions about response to virus, treatment, and vaccination. Most of these models use patient data to determine population-level parameterization, that is, parameter values that are meant to reflect the within-host responses of an “average” individual. New models are continually being created, so our sample of within-host models below is by no means a comprehensive list, but is meant to provide the context that motivated us to build our model.

Li et al. [9] have a within-host viral dynamic model of live infection with SARS-CoV-2 using chest radiograph score data to determine model parameters. The focus of the model is on simulating viral growth within the lung. With a system of ordinary differential equations, it tracks three populations: uninfected and infected pulmonary epithelial cells and viral load. The model is used to explore the effect of treatment timing and patient immune strength, and was later analyzed mathematically in [10] with the aim of laying a foundation for exploring treatment interventions *in silico*. There are nine model parameters, seven of which are fit at a population level using data from two published studies [11] and [12] via Markov-Chain Monte Carlo optimizations.

Farhang-Sardroodi et al. [13] created a within-host model of the immune response to an adenovirus-based vaccine. Using ordinary differential equations, they investigated the impact of various dosing strategies, and captured dose-dependent responses. Model parameters were fit to clinical trial data for the AstraZeneca/Oxford vaccine [14]. Data from a binding and neutralization study collected from COVID-19 recovered patients [15] were used to compare antibody-level predictions. The overarching aim of the Farhang-Sarhoodi investigation was to determine how best to conserve vaccine doses while providing necessary levels of protection. The model includes seven population state variables: non-replicating vaccine cell particles, helper T cells, cytotoxic T cells, IFN-*γ*, IL-6, plasma B-cells, and an antibody population.The antibodies in this model are stimulated by plasma B-cells which are indirectly stimulated by the presence of vaccine cell particles. The model antibodies in turn clear or neutralize vaccine cell particles. There are twenty-one model parameters, twelve of which are found by fitting the model to data to determine population-level ranges.

The goal of the model by Kim et al. [16], which does not account for vaccination, is to use viral-load data to compare within-host dynamics of SARS-CoV2, MERS-CoV, and SARS-CoV with the aim of gaining more insight into SARS-CoV2 behaviors and improved treatment strategies. Using simplifying assumptions the model is reduced to a set of two bilinear ordinary differential equations that track levels over time of the number of coronavirus RNA copies and the fraction of infected target cells. With this simple form, the authors are able to insert treatment terms to explore hypothetical combination therapies. Patient data were fit simultaneously using a nonlinear mixed effects approach to determine model parameter values. With this model the authors determine that therapies that block virus production are likely to be effective only if initiated before the viral load peak. There are eight population-level parameter values for each of the three virus types.

The model by Sadria and Layton [17] is built to track the interactions between SARS-CoV-2 and the immune response, and is meant to provide a testbed for simulating the effects of drug treatments against SARS-CoV-2 infection. The strength of this model is in how comprehensive it is. The authors track the control of SARS-CoV-2 infection by both the innate and adaptive immune responses. Data from viral load studies are used to determine model parameters, and three treatment options are simulated: Remdesivir, convalescent plasma, and a hypothetical therapy that inhibits virus entry into host cells. The populations tracked by the model include viral load, healthy cells, latent cells (which serve as hosts for replicating virus), infected cells, antigen presenting cells, interferon, effector cells, plasma cells, virus-specific antibodies that serve to neutralize and eliminate virus, the fraction of damaged cells, and a measure of specificity (a metric that increases as plasma cells produce antibodies that are more compatible with viral antigen). The model provides a foundation for the development of a platform for *in silico* testing of potential therapies and vaccines for COVID-19.There are twenty-nine population-level model parameters.

Other mathematical models of SARS-CoV-2 within-host dynamics include [18], [19], [20], [21], and [22] with each model focusing on somewhat different questions. Model parameters are fit to a variety of data sets measuring viral load, antibody levels, and certain immune response rates, and tend to be determined at a population level.

We have learned a great deal since the start of the world-wide outbreak about how SARS-CoV-2 affects individuals, but it is still not well-understood why some infected individuals experience only mild symptoms, with some even remaining asymptomatic, while for others, infection with SARS-CoV-2 can result in severe respiratory symptoms and even death. As Chatterjee et al. [22] point out, the reasons for the extreme heterogeneity in outcomes of SARS-CoV-2 infection across individuals is still unclear, but they hypothesize that this heterogeneity arises from variations in the strength and timing of an individual’s immune response. Most within-host models provide model parameterization for immune responses to infection at a population level, but we want to move toward being able to help an individual understand how robust their own immune response is likely to be. Our aim is to provide a mathematical model that is as simple as possible (our model has only two state variables), with as few parameters as possible (only four subject-specific parameters must be fit for an individual), so that determining subject-specific parameters is a tractable task. With the model in this paper we can provide subject-specific insight regarding individual levels of likely immune protection against infection in a fully vaccinated individual using neutralizing antibody (NAb) levels as our indicator.

## 2. Materials and Methods

### Assay Data

The AditxtScore^™^ test for neutralizing antibodies to SARS-CoV-2 is a novel flow cytometry based competitive inhibition assay for the measurement of total neutralizing antibodies to SARS-CoV-2 in human plasma samples. Microparticles coated with the recombinant SARS-CoV-2 RBD antigen are incubated with biotinylated angiotensin converting enzyme-2 (ACE-2), human subject plasma or phosphate buffered saline (PBS), and fluorescent labeled streptavidin. Neutralizing antibodies in the subject plasma sample bind to the RBD antigen and inhibit binding of ACE-2 to the RBD antigen. Following incubation, the beads are washed and then measured by flow cytometry to determine the degree of inhibition of ACE-2 binding. The degree of inhibition of the ACE-2 binding is proportional to the amount of neutralizing antibodies present in the human subject sample. Zero or near zero percent ACE-2 binding inhibition is observed when phosphate buffered saline is used as sample or when no neutralizing antibodies are present in the subject sample. Human subject plasma samples with higher concentrations of neutralizing antibodies will produce % binding inhibition values up to 100%. One hundred percent binding inhibition is achieved when ACE-2 is completely inhibited from binding to the RBD coated microparticles by the neutralizing antibodies in the sample. Sample % ACE-2 binding inhibition values are compared to a standard curve with known neutralizing antibody values in International Units per milliliter (IU/mL) to convert percent inhibition values into units of IU/mL for each subject sample tested. A standard curve was generated using dilutions of the human NIH SARS-CoV-2 serology standard, Lot # COVID-NS01097, characterized and made available by Frederick National Laboratory for Cancer Research (FNLCR), Frederick, Maryland, USA. The FNLCR standard has been assigned Potency for Functional Activity (Neutralizing Unitage) of 813 IU/ml as calibrated to the primary standard WHO SARS-CoV-2 Serology Standard. A dilution series of standard was prepared with standard concentrations accounting for values of 2032.5, 1626, 1016, 813, 406.5, 203.2, 101.6, 50.8, and 25.4 IU/mL. IU/mL values were plotted against neutralizing antibody % inhibition values measured by flow cytometry for each standard dilution. The resulting standard curve was fit using a polynomial curve fit function and the curve equation was used to generate IU/mL values from % inhibition values for all samples measured. Figure 1 shows the calibration curve derived over four days of runs and the resulting polynomial equation.

**Figure 1:**
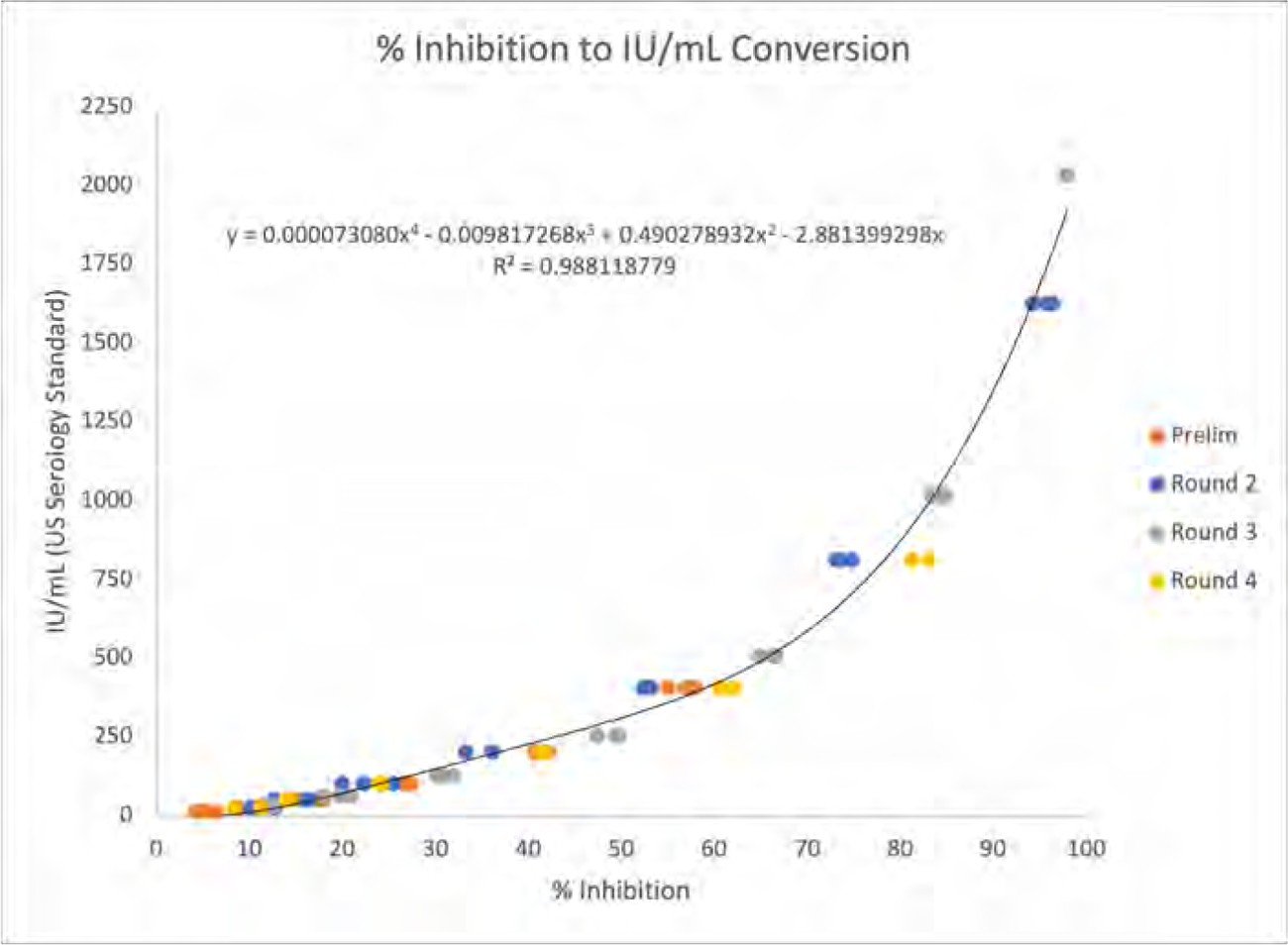
Calibration curve for converting % inhibition to IU/mL. Resulting polynomial fit: *y = (7.3 × 10^−5^)x^4^ − (9.8 × 10^−3^)x^3^ + (4.9 × 10^−1^)x^2^ − 2.9x*.

Precision of the method was determined using 4 subjects with 4 different IU/mL values, 6 assays per day for 3 days. Precision varied somewhat among subjects (details are included in Appendix B). The coefficient of variation (%CV) was inversely proportional to IU/mL, thus precision is higher for higher NAb levels in IU/mL. For our visualizations of the data, we used a coarse-grained assignment of precision results for four concentration ranges. Values are shown in Table 1:

**Table 1:** Coarse-grained precision of method to determine AditxtScore^™^ for neutralizing antibodies (NAb) to SARS-CoV-2, given in four IU/mL interval ranges. Precision as measured by coefficient of variation (% CV) increases with IU/mL.

| IU/mL | Avg. %CV |
| --- | --- |
| 0 – 89 | 20 |
| 90 – 149 | 15 |
| 150 – 699 | 10 |
| 700 – 2200 | 5 |

### NAb cut-point values

Cut-points for NAb concentrations were based in part on an analysis by Khoury et al. [1], where NAb activity elicited by seven different SARS-CoV-2 vaccines and a reference group of convalesced non-vaccinated subjects was associated with the observed reduction in subsequent infections over the next several months compared to placebo. By determining NAb concentrations in a comparable cohort of convalesced subjects and referring to their analysis, we estimated that for the wild type SARS-CoV-2, NAb values 90 IU/mL - 150 IU/mL were associated with 60 *−* 75% protection are were considered to be in the “weak response (WR)” range, while values above 150 IU/mL were associated with *>* 75% protection and were considered a “positive response (R),” and values above 300 IU/mL were associated with *>* 90% protection and were considered to be a “strong response (SR).” The 90 IU/mL cutpoint was validated against a set of known positive/negative samples made available by the Frederick National Laboratory for Cancer Research (FNLCR) with a sensitivity and specificity of 100%.

### Mathematical Model

The mathematical model we introduce is a two-state system describing NAb response to vaccine. The system of ordinary differential equations captures the high-level mechanistic dynamics of a vaccine-triggered immune response within an individual. Model states are:

- A: Neutralizing antibody (NAb). Units: IU/mL
- V: Proxy for transfected cells in response to mRNA vaccine. Units: mL

The dynamics of NAb over time are described by two nonlinear ordinary differential equations. The time scale is a 24-hour day:

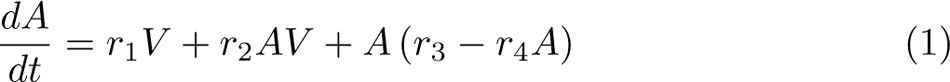

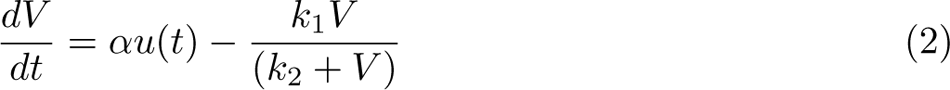

Here *αu*(*t*) represents a normalized vaccine dose in mL/day. The role of the vaccine dose in this model is simply as a trigger to engage an immune response; we do not explore modeling the effect of varying dosage levels. State variable *V* (*t*) in mL is a proxy for the activity of cells transfected as a result of vaccination, and therefore *V* decays with a Michaelis-Menten dynamic. In a traditional dose-response model, a term of this form would capture the pharmakokinetics of medication in the system, but mRNA vaccines behave differently, so *V* is not the “amount of vaccine in the body.” Nonetheless, for the purposes of predicting antibody dynamics over time, the system as modeled reflects the initiation of the immune response with the Michaelis-Menten clearance term conceptualized as “vaccine clearance,” and has the same form as equation (1) in [23] that captures drug concentration pharmacokinetics. The presence of the transfected cells *V* initiates an antibody response, represented by state variable *A*(*t*). The dynamics of the antibody population within the individual, once triggered, are represented with a logistic term.

In particular, in equation (1):

- *r*_1_*V* represents initiation of antibody activity in response to vaccine.
- *r*_2_*AV* represents an antibody boost in response to vaccine.
- *A* (*r*_3_ *− r*_4_*A*) models intrinsic antibody dynamics as logistic. In equation (2):
- *αu*(*t*) represents a vaccine dose that drives the presence of transfected cells. This is modeled by a discrete pulse on the days vaccine is administered, and has value 0 otherwise.
- 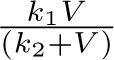 is a Michaelis-Menten type decrease in transfected cells over time.

The model has a total of four parameters that we consider subject-specific, and to which we fit individual data:

| Parameter | Units | Description |
| --- | --- | --- |
| $r_1$ | IU/(day mL<sup>2</sup>) | Antibody initialization in response to vaccine dose |
| $r_2$ | 1/(day mL) | Increase in existing antibody in response to vaccine dose |
| $r_3$ | 1/day | Proportional to intrinsic antibody growth rate |
| $r_4$ | mL/(day IU) | Proportional to intrinsic antibody decay rate and intra-species competition |

Three parameters are fixed at the same values for all subjects. Parameter *α* scales the vaccine dose *u*. Parameters *k*_1_ and *k*_2_ scale Michaelis-Menten decay dynamics for state variable *V*. Function *u* is a proxy for a vaccine dose, state variable *V* is a proxy for the subsequent transfection of cells. Our interest is in tracking the subject-specific neutralizing antibody response to vaccine doses in equation (1), so for equation (2) we simply select parameter values that produce simulation outcomes that are biologically reasonable. We therefore choose *α, k*_1_, and *k*_2_ to be fixed at values that are consistent with those of other pharmakokinetic drug models, and are within the ranges used in [23]. Keeping the parameter values in equation (2) the same for all subjects allows us to achieve good model fits to data by varying only parameters *r_i_, i* = 1 *· · ·* 4 in equation (1). (Parameter values used in the simulations are included in Appendix A.)

| Parameter | Units | Description |
| --- | --- | --- |
| $k_1$ | 1/day | Michaelis-Menten scaling for clearance of $V$. |
| $k_2$ | mL | Michaelis-Menten half-saturation. |

### 2.1. Parameter Fitting and Numerical Solution

Subject-specific parameters were fit with a variety of built-in MATLAB methods that are part of the Global Optimization toolbox [24]. These included Multistart, Particle Swarm Optimization, and Global Optimization, in addition to MATLAB’s `fmincon` and `lsqnonlin` for refining initial parameter guesses. Different cost functions were tested. The ODE solutions we present used a weighted sum of squares cost function for the global optimization that includes a penalty for solutions that are larger at some time points than we consider physically feasible. Error reported is the average unweighted square root of the sum of squares. Of each of the parameter fitting approaches, the Global Optimization approach tended to yield the best fit solutions, which we will present. Markov-Chain Monte Carlo (MCMC) methods can be used to enhance the process of fitting models to data. MCMC methods are sampling methods, and are not primarily used to find the best fit parameters to a particular data set. Optimizers such as those cited above are better choices for finding optimal parameter values. Starting with optimum parameter choices, MCMC produces a chain (a large set) of likely parameter combinations and generates a distribution of model outcomes by sampling parameter combinations from the chain. Using the package `mcmcstat` [25] for MATLAB, we ran MCMC on each subject’s data set to produce predictive envelopes for model outcomes. The `mcmcstat` package provides tools to generate and analyze Metropolis-Hastings MCMC chains using multivariate Gaussian proposal distribution [26] [27]. The parameter values found by MATLAB’s Global Optimization routine were used as initial parameter guesses. For our data sets, we set a burn-in time of 500 iterations, and generated an MCMC chain of length 5000. Predictive envelopes were determined by sampling the chain 1000 times, then determining predictive quartiles of 50%, 90%, 95%, and 99%.

Model simulations were run using a stiff ODE solver `ode15s` in MATLAB 2021a [28]. The assay data provide results in percent neutralization, which ranges from 0 to 100. To increase numerical stability, the data were divided by 10 which scaled them linearly to range between 0 and 10. After solutions were computed and parameter values were found, results were then scaled up to units of IU/mL via the percent neutralization to IU/mL conversion function described in section 2. This function can take on values between 0 and 2200.

## 3. Results

### Global Optimization Model Fits to Data

The four subject-specific parameters representing the strength of response to vaccine and the intrinsic dynamics of antibody levels were fit to time-series NAb data collected through venous blood draws from 27 subjects over a period of several months. Each subject was, to the best of our knowledge, not previously infected with SARS-CoV-2, and each subject received two mRNA vaccine doses: some received two Pfizer doses, and some received two Moderna doses. From a sample of 27 subjects, 9 received Pfizer and the rest received Moderna. Using this sample size, we were not able to detect any significant differences in NAb trajectories correlated to whether the vaccination was Pfizer or Moderna. The number of samples, the number of days between samples, and the timing of the two vaccine doses varied from subject to subject. Once model development was completed and run on the 27 subject data sets, we further tested the model with NAb values from 6 additional subjects (1 received Pfizer, 5 received Moderna) and were able to achieve good fits without modification to the model.

In Figures 2 to 5 we explored the immune protection categories to which a subject belonged at the time a subject’s last sample was taken, and the immune protection category predicted by the model simulation projected to day 400. The first day of the model simulation is day 0. Since the sample dates of the subjects are heterogeneous, there is also variation in the time gap between the day the final sample was collected and day 400 of the simulation. In the examples we show, we see varying persistence of immune response between the final measurement of NAb levels and the projected NAb levels.

**Figure 2:**
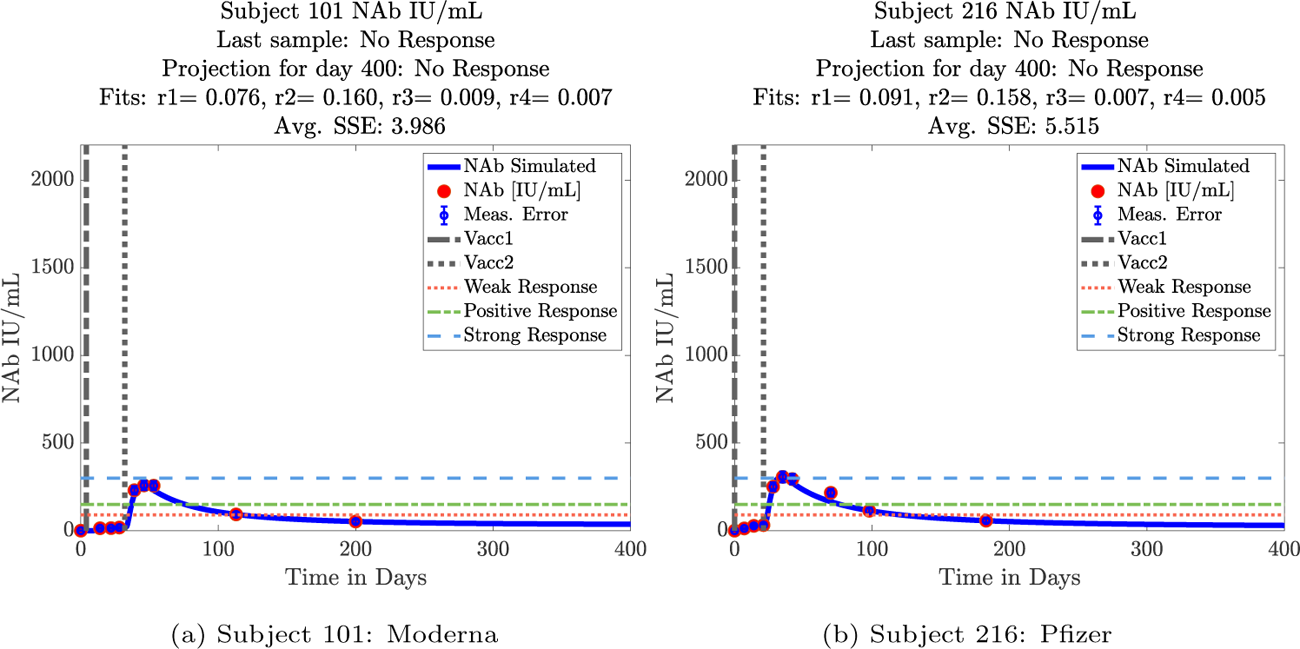
Subject category: NR-NR. For these subjects, both the last sample taken, and the simulated prediction of NAb levels on day 400 indicate NAb levels below the 90 IU/mL threshold.

Figure 2 shows two example subjects in the “NR-NR” grouping. These are individuals whose last collected sample around day 200 put them into the noresponse (NR) category, and the projections, as we might expect, also put these subjects into the NR category by day 400.

In Figure 3 the last sample collected for subjects 146 and 200 indicated a weak response. The projection for subject 146 (3a) drops to no response (NR) by day 400, whereas subject 200 (3b) is projected to persist in the weak response (WR) range.

**Figure 3:**
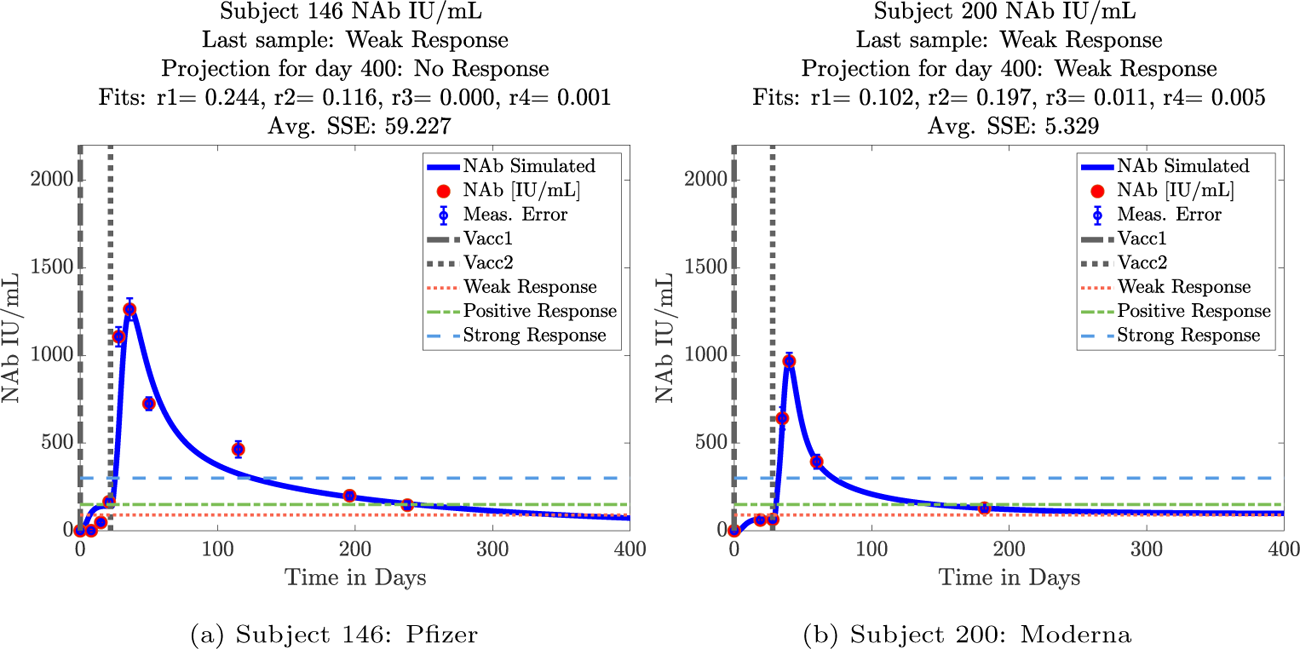
Subject category: WR-NR and WR-WR. For these subjects, the last sample collected indicated a weak response. The projection for subject 146 (3a) predicts dropping to no response (NR) by day 400, whereas subject 200 (3b) is projected to persist in the weak response (WR) range.

In Figure 4 we see two subjects whose final collected sample puts them into the positive response (R) category, but the response of subject 19 is projected to decline to a weak response by day 400, whereas subject 76 is projected to persist in the positive response category through day 400.

**Figure 4:**
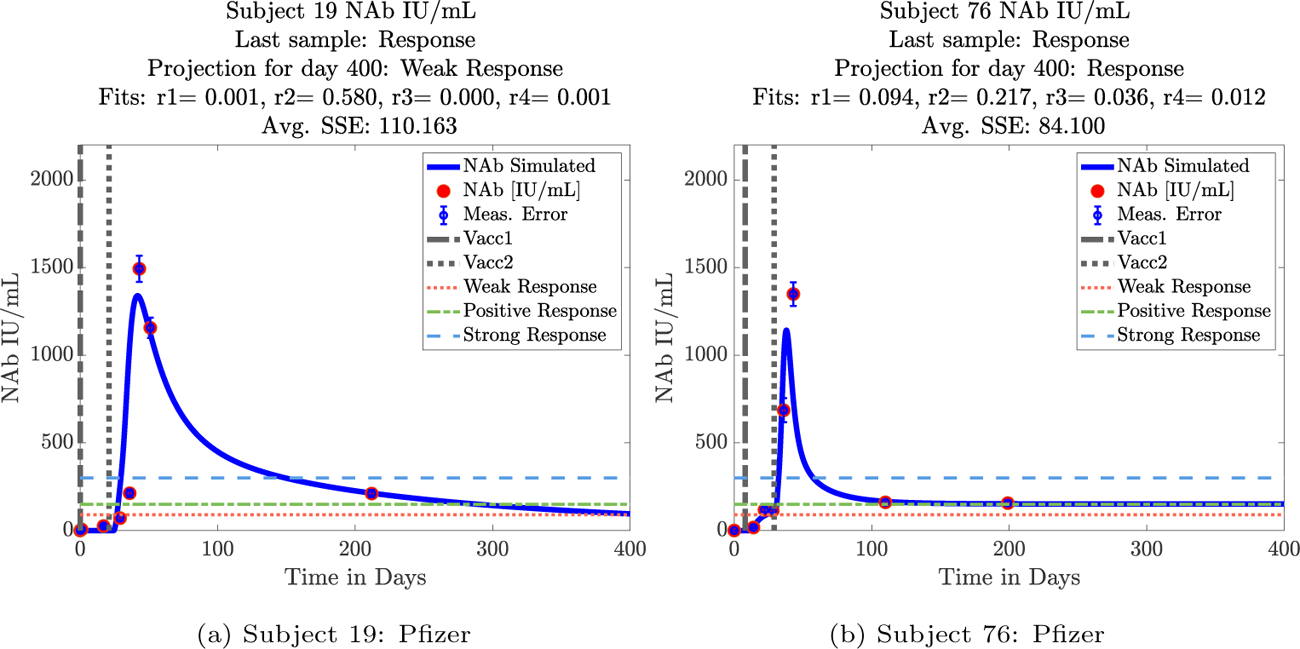
Subject category: R-WR and R-R. For these subjects, the last sample taken around day 200 showed a positive response (NAb between 150 IU/mL and 300 IU/mL). For subject 19 in panel (4a) projected NAb levels drop to between 90 IU/mL and 150 IU/mL, the weak response range. Subject 76 in panel (4b) is projected to remain in the positive response range.

In Figure 5 we have three subjects, all of whom were in the strong response (SR) category when their final samples were collected (between days 100 and 200). We see, however, different rates of immune strength decline. Subject 226 still shows a strong immune response when the final sample is collected around day 125, but is predicted to drop into the weak response range by day 400. Subject 100 also shows a strong response in the final collected sample around day 200, yet their response decays to the positive response range by day 400. Subject 23 shows a strong response in the final collected sample, just after day 200, and simulations predict they should remain in the strong response range by day 400.

**Figure 5:**
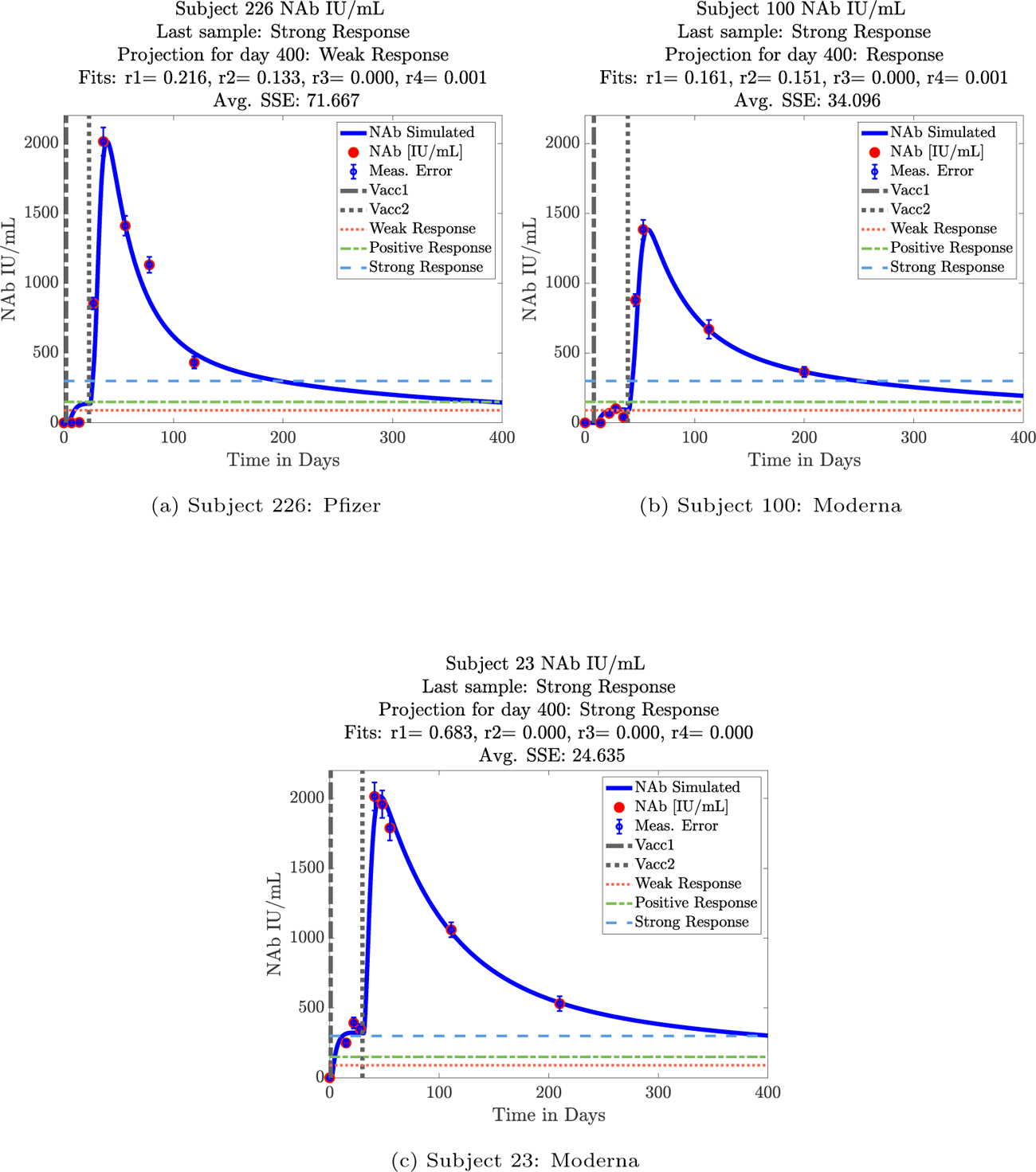
Subject category: SR-WR, SR-R, SR-SR. For these subjects, the last sample taken indicated a strong response with NAb levels above 300 IU/mL. For subject 226 (5a), the simulation predicts a drop to the weak response range (WR) by day 400. Subject 100 (5b) drops from strong response to positive response (R) by day 400, while for subject 23 (5c), the simulated prediction of NAb levels on day 400 remained above 300 IU/mL, the strong response range (SR).

### MCMC Model Fits to Data

Next we observed the results of using MCMC on the same subject data set with the same underlying mathematical model. Initial guesses for parameter values used were the values generated by the MATLAB Global Optimization fits. Although Global Optimization computations yielded better overall fits to data, the advantage of the MCMC approach is that we can use the chain of parameter combinations (5000 possible combinations of the four fit parameters in our case) to generate predictive quartiles of 50%, 90%, 95%, and 99%. The quartiles are shown in gray bands of varying darkness, with the lighter band encompassing 99% of the chain samples, and the darkest band holding 50% of the chain samples. The black line inside the bands connects the median of the chain samples at each data point. Using the predictive quartile envelopes generated by the MCMC approach, we can provide subject-specific likely protective immunity intervals, or “envelopes.” We carried out the MCMC algorithm on all 33 subjects, and present a representative subset of subjects from categories of the global optimization fittings already shown. From the no response (NR) category, we show subjects 65 and 179 in Figure 6. The predictive envelope places these subjects between having no response to having a weak response by day 400 with 99% likelihood. From the positive response category, we looked at subjects 100 and 139 in Figure 7. The predictive envelope places these subjects within a range from a positive response to a strong response with 99% likelihood. From the strong response category, we see in Figure 8 that subjects 21 and 23 are both predicted to remain solidly within the strong response band with 99% likelihood.

**Figure 6:**
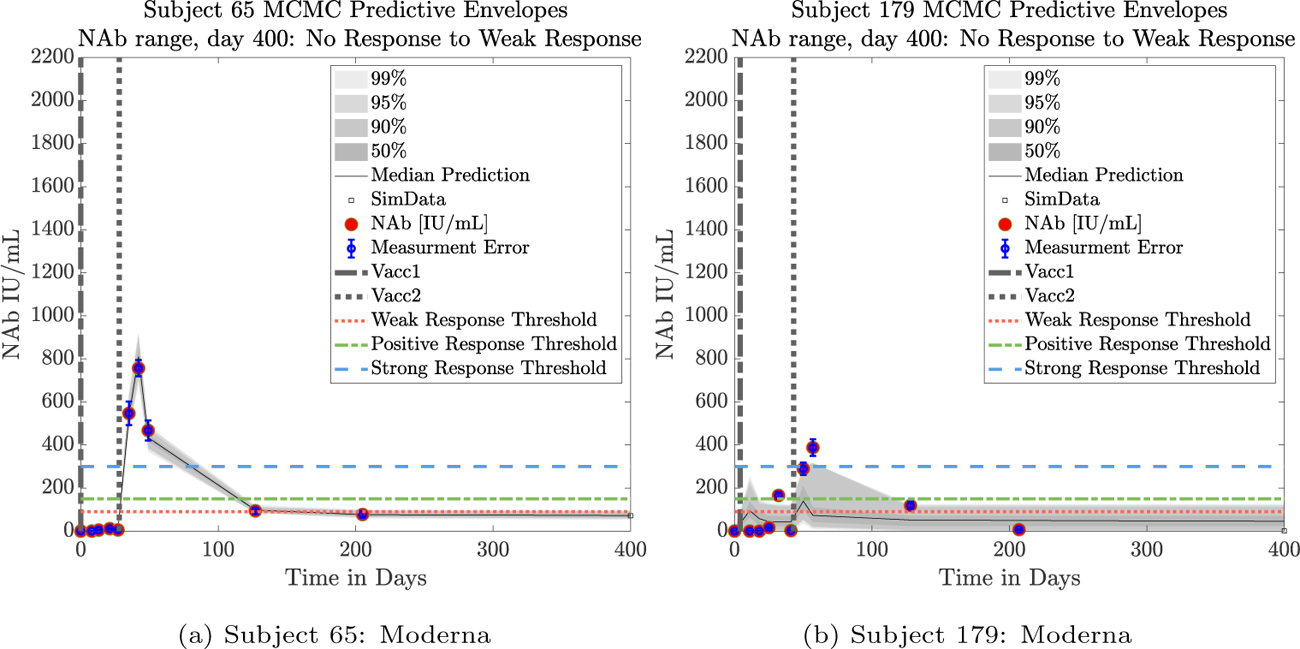
Predictive Envelope Range: No response to weak response predicted at day 400.

**Figure 7:**
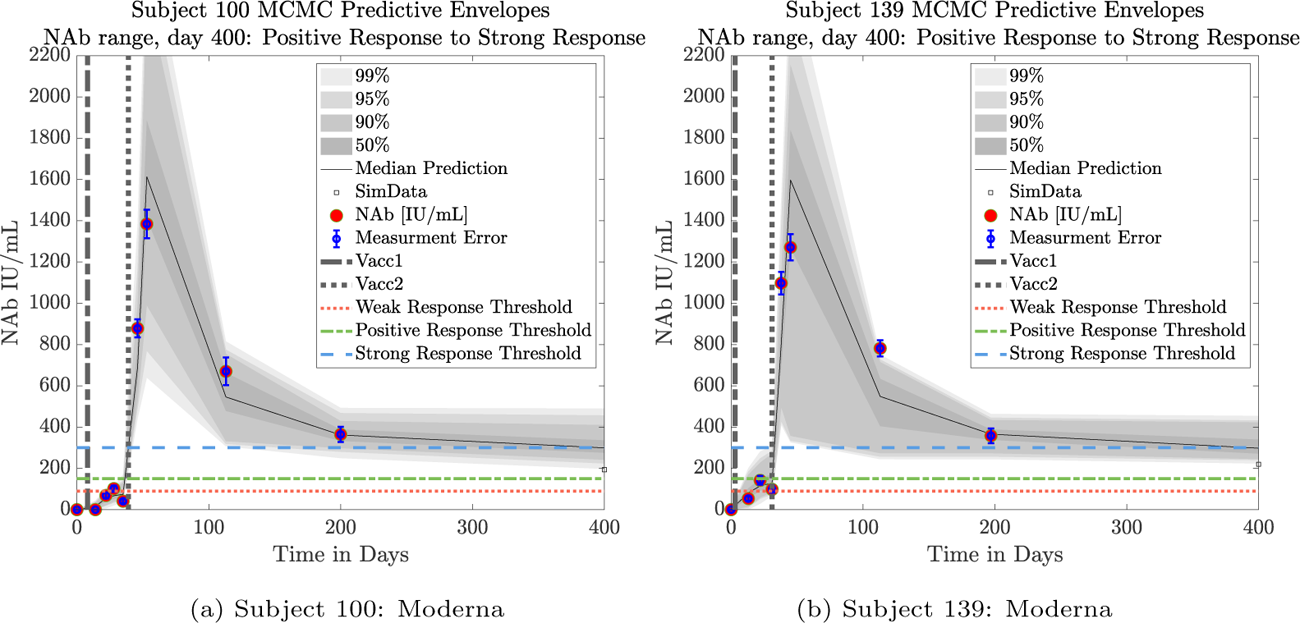
Predictive Envelope Range: Positive response to strong response predicted at day 400.

**Figure 8:**
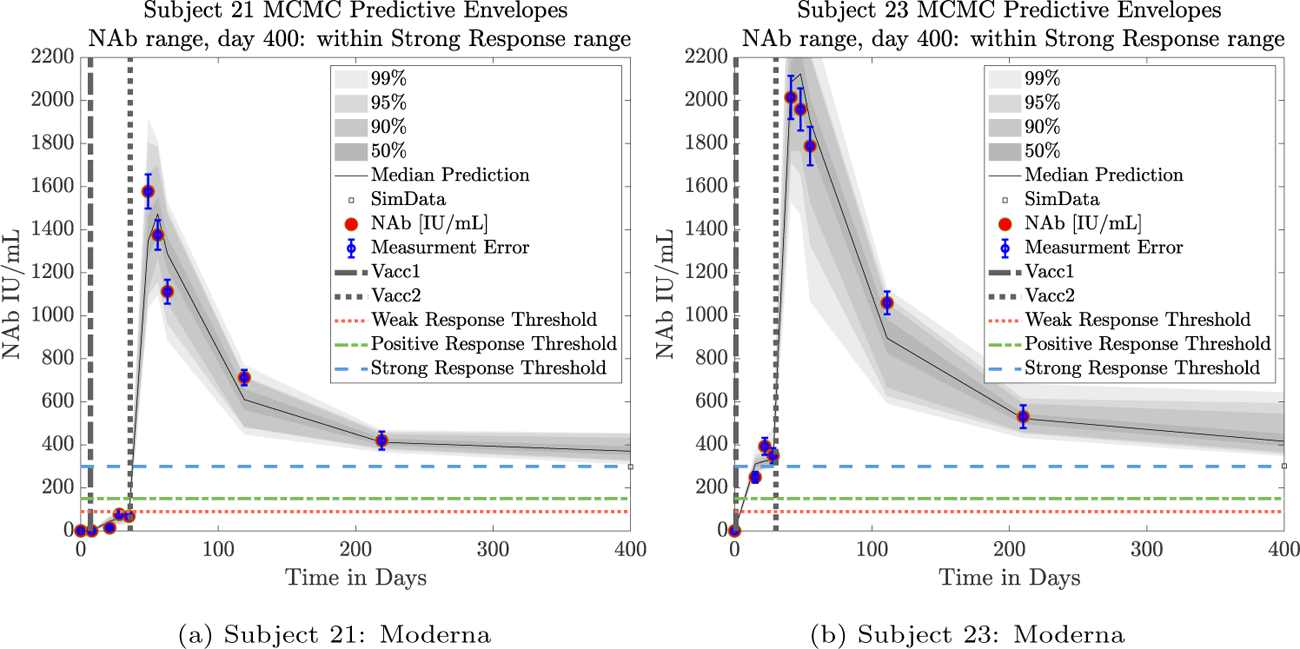
Predictive Envelope Range: Strong response predicted at day 400.

## 4. Discussion

Our new mathematical model tracks levels of subject-specific neutralizing antibodies within an individual over time. Starting with 27 sets of longitudinal NAb level data from twice-vaccinated individuals, we developed our model to simulate NAb level changes in IU/mL to help predict the subject-specific within-host dynamics of antibody decay. Once the model was developed and tested on the 27 data sets, we tested the model on 6 additional longitudinal NAb level data sets, giving us a total of 33 data sets on which we ran the model. The model is sufficiently simple to be tractable, and can provide a high-level view of changing NAb levels within a specific host by fitting only four parameters to individual subject data. One of the advantages of the novel Aditxt^™^ flow cytometry assay is that one can run more assays for less cost, thereby making the collection of longitudinal subject data easier. Since our model does well with more longitudinal data points, the availability of subject-specific data from this type of assay makes the implementation of our model practical. The model can be considered mechanistic in that the parameters can be connected to biological meaning. The model state representing the action of an mRNA vaccine serves as a proxy for transfected cells. The model state representing the levels of NAbs in a system in IU/mL is a proxy for the neutralizing strength of the NAbs present in the system. The timing of the delivery of first and second vaccine doses varies from individual to individual. Even the with non-uniform collection of the number of samples and timing between samples, and the varying time gaps between the first and second vaccine administration for each subject, the model has sufficient flexibility to be able to achieve good fits to subject data.

We applied two model fitting schemes (global optimization and MCMC) to the 33 data sets, each subject data set having between four and eleven time points at which NAb levels were measured. The global optimization approach yielded very good fits to subject data, and the MCMC approach produced predictive envelopes for likely subject-specific model outcomes.

The model is simple yet sufficiently complex to be able to capture a range of dynamics evidenced in the cohort of 33 subjects. Our analysis brought to light some NAb dynamic patterns worth noting. These patterns could be explored more deeply once more subject data sets are collected. The first of these is a connection between married couples in the same household. The second is an observation that although a relatively weak initial response to a second vaccine dose tends to predict weak persistence of NAb levels over time, strong responses do not necessarily guarantee strong NAb persistence.

### Shared Household Married Couples: Sex Differences

The cohort data set includes 5 male-female married couples. Understanding sex difference in response to vaccine can be complicated because of so many confounding factors that vary from individual to individual, including risks encountered via household practices, profession, and quality of adherence to safety protocols. Since these married couples live in the same household, one factor is normalized. The comparison within male-female married couples revealed a consistent pattern with projections that were simulated over a 400-day time span. From this small sample set, it appears that the response in females tends to be more robust than that in the male partner. This observation is consistent with prior studies demonstrating stronger immune responses, and specifically stronger antibody responses to vaccines, in women, *c.f.*[29]. This distinction shows up more clearly when comparing individuals within married-couple pairs, but the sex difference is not as clear when these subjects are mixed back into the general population. In Figure 9 comparing subjects 4 (male partner) and 5 (female partner), it is clear that both the initial response to vaccine and persistence over time is stronger in subject 5. In Figure 10, the difference in response between married subjects 20 and 21 is not as strong, and it may appear that the male partner has a more robust response, but the predictive envelopes indicate a similarly strong response in the female partner. In Figure 11 it is apparent that subject 30 (female partner) has a stronger overall NAb response than does subject 25 (male partner). The difference in response is also distinct in Figure 12 between subject 64 (female partner) whose initial response was much stronger than that of subject (65) male partner, and persistence is somewhat stronger as well. In Figure 13, subject 173 (female partner) has a strong persistent response to vaccine whereas subject 175 (male partner) has both an initially relatively low response and by day 400 no response is predicted. The pattern of the stronger female response in the majority of cases indicates that further exploration of within-household married couple responses may be warranted.

**Figure 9:**
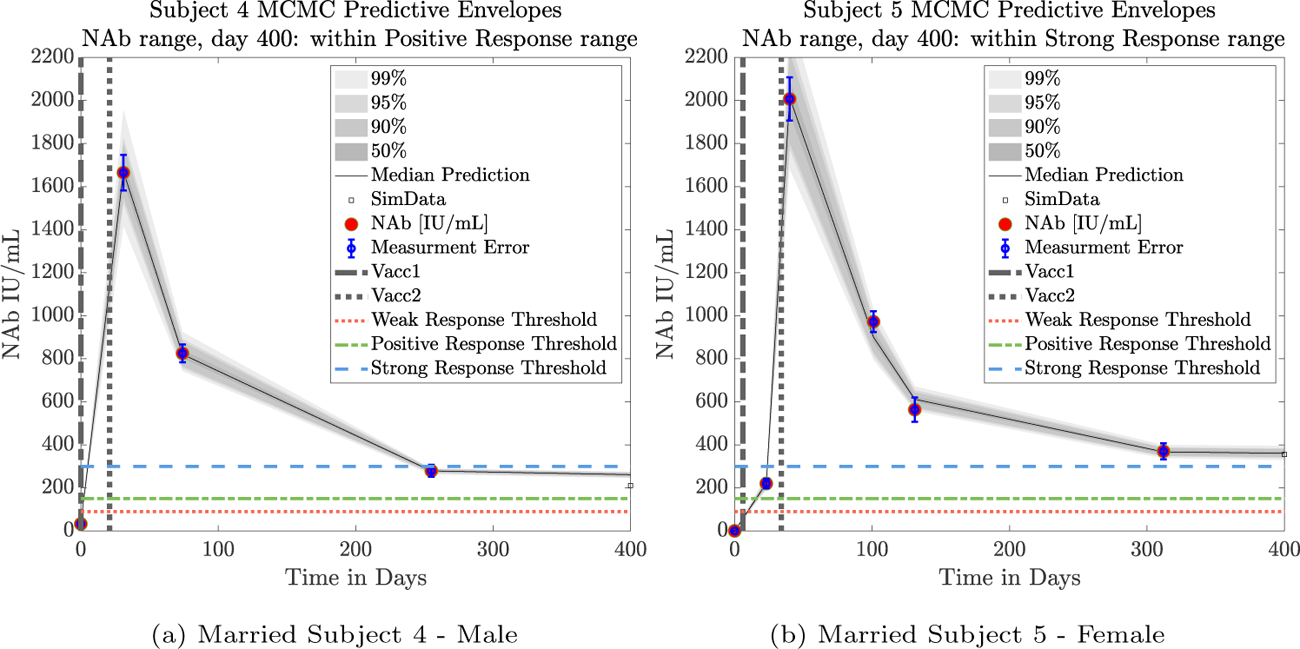
Married Subjects 4 (M), 5 (F): Male-Female comparison of response predicted through day 400.

**Figure 10:**
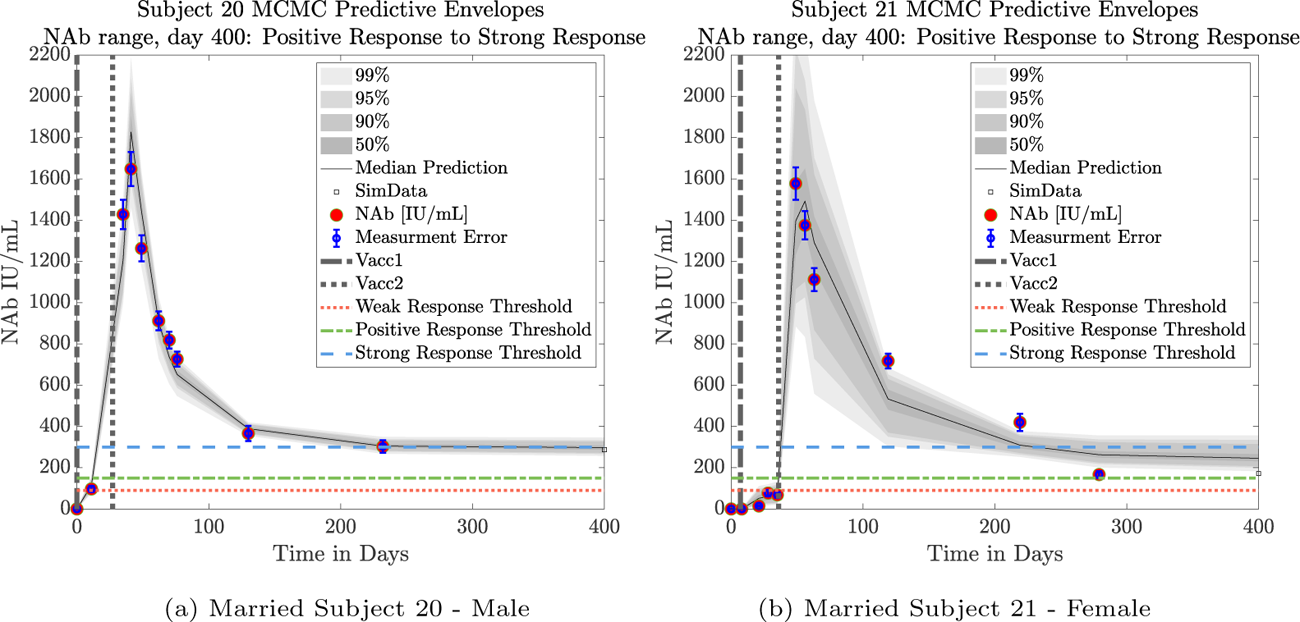
Married Subjects 20 (M), 21 (F): Male-Female comparison of response predicted through day 400.

**Figure 11:**
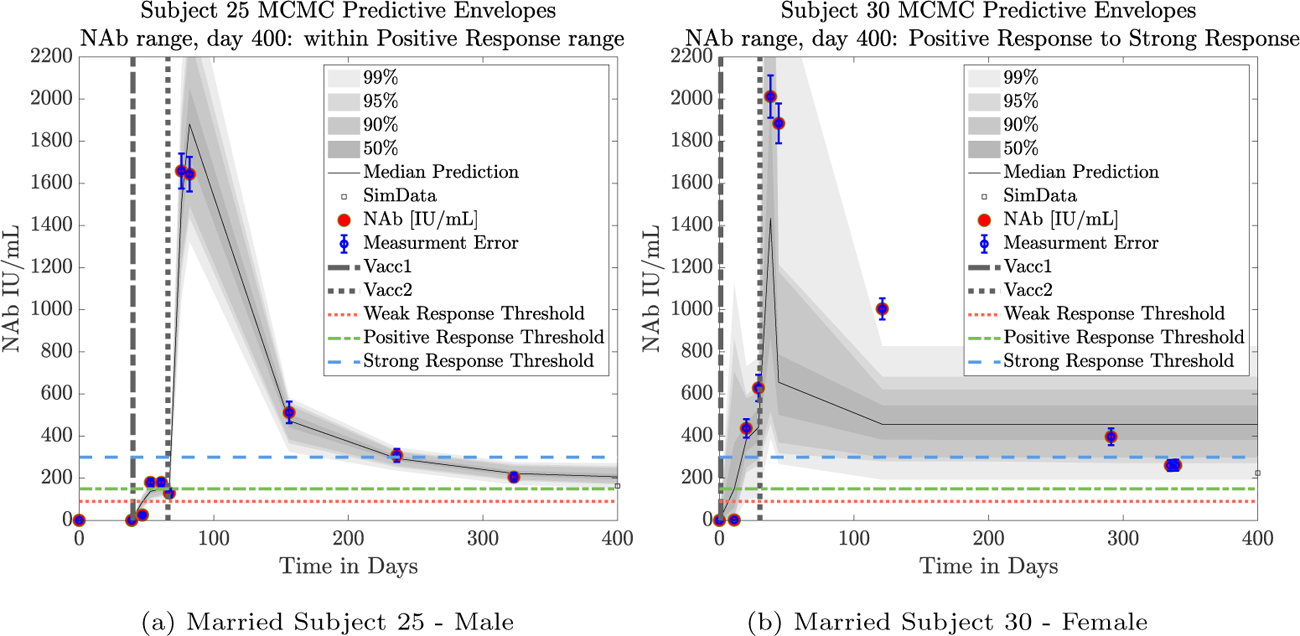
Married Subjects 25 (M), 30 (F): Male-Female comparison of response predicted through day 400.

**Figure 12:**
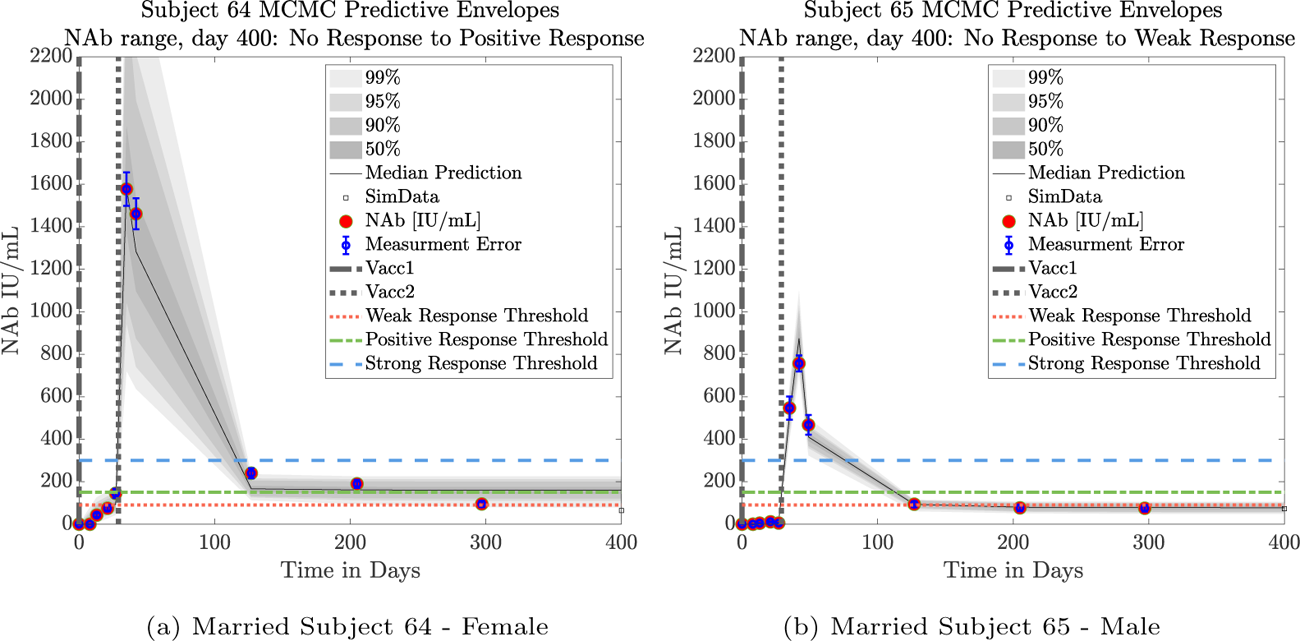
Married Subjects 64 (F), 65 (M): Female-Male comparison of response predicted through day 400.

**Figure 13:**
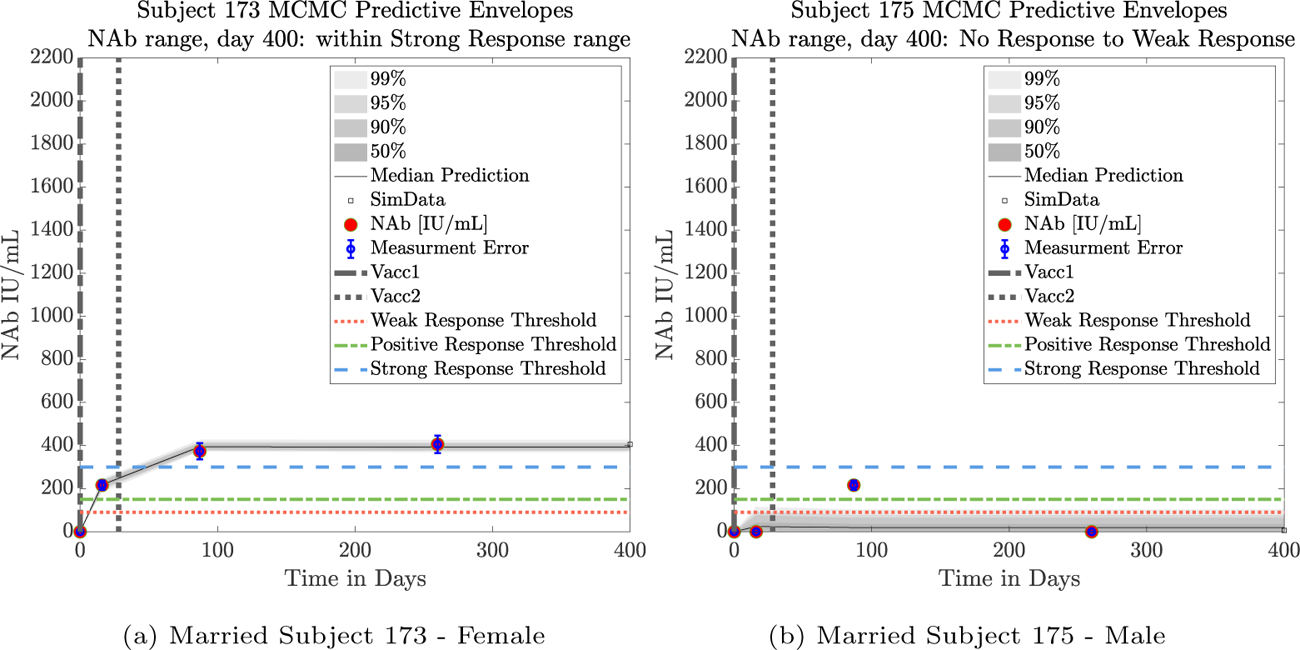
Married Subjects 173 (F), 175 (M): Female-Male comparison of response predicted through day 400.

### Strong Initial Response versus Persistence

Model simulations show a fairly weak maximum NAb response to the first vaccine dose among nearly all individuals in the cohort, indicating at best low levels of protection after only one administration of the vaccine. The second vaccine dose shows a much stronger response, both in maximum NAb level and in persistence of higher levels. This behavior is captured by the model as a response to a vaccine challenge when there is preliminary immune system priming in place. In the data sets we analyzed, it was clear that an initially strong response to the second vaccine dose did not always predict high NAb levels over time. In certain cases, a strong response persisted, but in others, an initially strong response can be seen to decline to nearly no response several months later, whereas lesser initial responses can persist. When the initial response is relatively low, NAb levels remain low. Subjects 23, 44, 79 and 226 highlight the fact that a weak initial response is a good indicator that the response will remain weak, but a strong initial response does not guarantee persistence of high levels of NAbs in an individual over time. Subject 23 in Figure 14a shows a very robust response to the second vaccine dose, and by day 400, NAb levels remain above the strong response threshold. Subject 44 in Figure 14b shows a weak response to the second vaccine dose, and as expected, by day 400, the response had dropped below the weak response threshold. We also note that for these two subjects the MCMC-determined median outcomes of the simulations coincides with the best-fit simulations from Global Optimization. When we compare subjects 79 in Figure 15a and 226 in Figure 15b, however, we see that a stronger initial response does not guarantee stronger persistence. In addition, the MCMC-determined median outcomes for these two subjects do not coincide as closely with the best-fit simulation determined by Global Optimization. We also note that the predictive envelopes for these two subjects cover much broader ranges than do the envelopes for subjects 23 and 44. Subject 226, in particular, has a 99% predictive envelope that at day 400 covers all response thresholds from weak to strong. The best fit simulation, however, projects that NAb levels in subject 226 will have declined to the weak response category by day 400.

**Figure 14:**
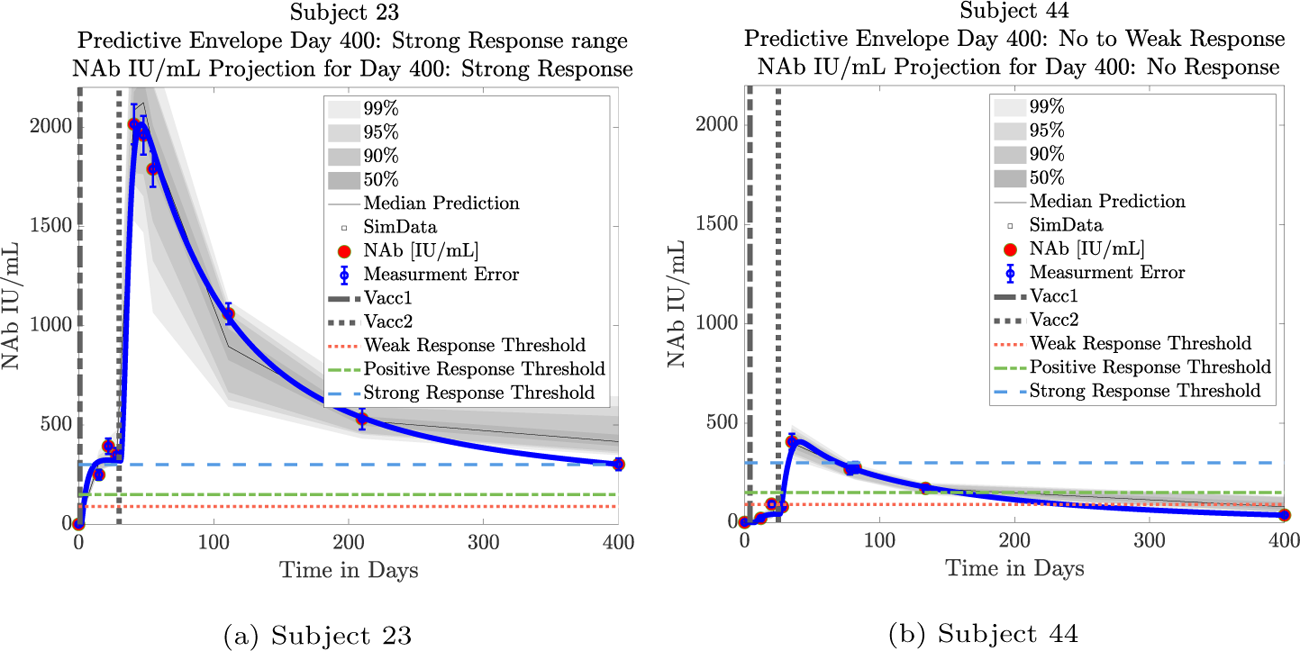
Predictive Envelopes with Best Fit Curve. Subjects 23 and 44: Strong and weak initial responses correspond to long-term high and low NAb levels, respectively. A very strong initial response in subject 23 to the second vaccine dose persists with strong protective immunity over time. A relatively weak initial response in subject 44 to the second vaccine dose continues to show weak protective immunity over time.

**Figure 15:**
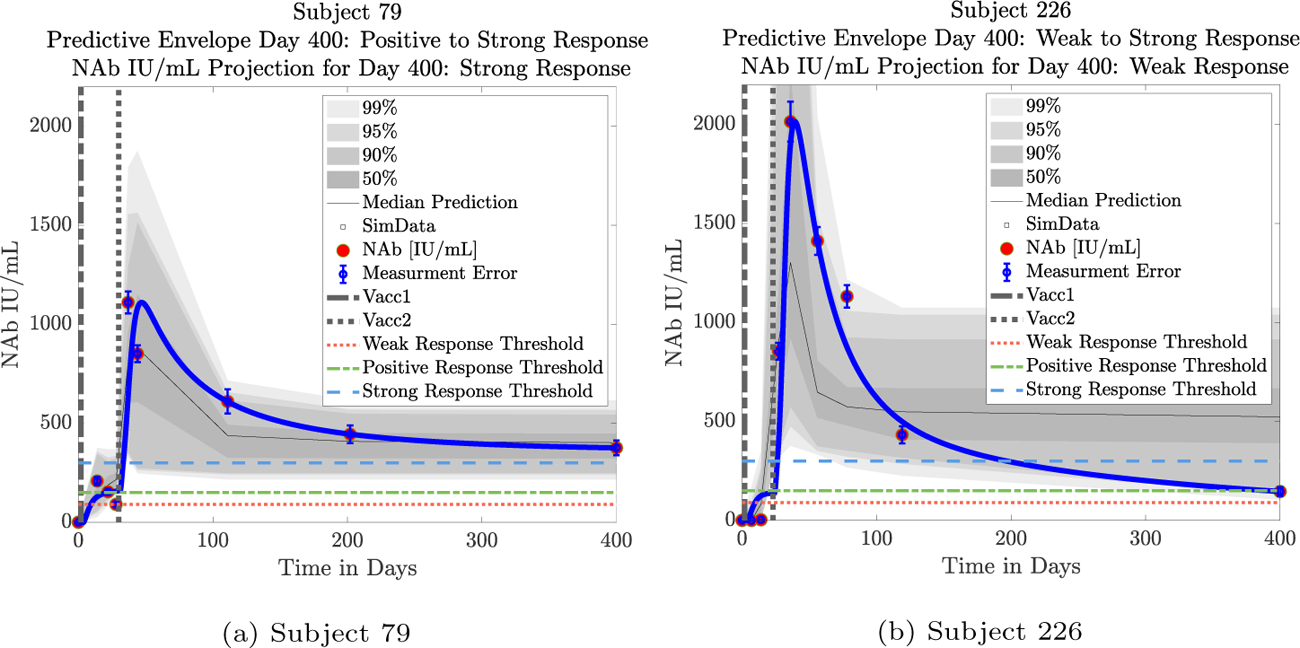
Predictive Envelopes with Best Fit Curve. Subjects 79 and 226: Stronger initial responses do not predict stronger long-term protective immunity over time. A moderate initial response in subject 79 to the second vaccine dose persists with strong protective immunity over time. A very strong initial response in subject 226 to the second vaccine dose declines to weak protective immunity over time.

### Future Directions

Vaccine efficacy for immunity against infectious agents is primarily assessed in clinical studies designed to evaluate reduction in disease incidence compared to placebo. Determination of differences between subjects that receive the active product versus placebo requires a large sample size, which extends the time and cost required for vaccine development. The ability to enroll a high number of subjects in these clinical trials is easier when infection rates are prevalent in pandemic conditions; however, subject recruitment becomes more challenging in the absence of a pandemic. Furthermore, clinical studies are not ethically feasible if the pathogen is extremely virulent and/or deadly. Finally, observation of changes in the incidence of infection may not provide information on the level of protective immunity for each individual subject. Immunity status is not all or nothing and can be impacted by a variety of factors including viral load and replication rate, level of initial immune response elicited by the vaccine, which may vary from one individual to another, time from immunization, mode of immunization, and other factors. Identification of a correlate of protection can reduce the number of subjects and evaluation time needed for clinical trials. A correlate of protection can also provide information about individual immune responses.

The purpose of this study was to select a correlate, neutralizing antibody (NAb), that has been reported in the literature as a measure of protection, and to create a mathematical model to determine its trajectory over time. This mathematical model can potentially help determine the rate of decay of NAb levels in individuals as well as responses in a population. The model was able to achieve good fits to individual subject data sets, even though there was significant individual variability in NAb dynamics, including in the strength of response to vaccine and persistence of NAb levels. We saw that long-term NAb persistence could not necessarily be predicted by the strength of the initial NAb level increase immediately following a second vaccine dose. Khoury and coworkers [1] used NAb levels in individuals who had convalesced from SARS-CoV-2 infection and individuals who received one of 7 commonly used COVID-19 vaccines to estimate the likelihood of protection. Using their findings as a starting point, we established multiple cut-points in NAb levels using a novel flow-cytometry-based test. Multiple cut points were identified to categorize NAb levels as none to minimal, weak protective response and strong protective response against the wild type virus. We analyzed the decay rate of NAb in 33 subjects who had no known history of SARS-CoV-2 infection and who had been fully vaccinated with two doses of the Moderna or Pfizer mRNA vaccines, and hypothesized protective immunity status using these cut points. While these cut points are meant to serve as guidelines to categorize the levels of NAb at a given time with respect to likelihood of protection, the levels required for protection may be impacted by the SARS-CoV-2 variant. Future studies are planned to test the model in different cohorts and with different SARS-CoV-2 variants, as well as to correlate the protective levels with incidence of breakthrough infections. Once the model has been tested in the context of these additional scenarios, we see several potential model applications, including more rapid evaluation of immune responses to vaccination to reduce time and expense of vaccine development; determination of immune responses on a case by case basis to identify vulnerable populations; and, more accurate assessment of the timing of boosters, again on a personalized level.

## Appendices

### A. Parameter Values and Vaccine Type

Fixed parameter values for equation (2):

- *α* = 2; *k*_1_ = 193; *k*_2_ = 747.

Subject-specific parameter value fits for equation (1):

**Table 2:**
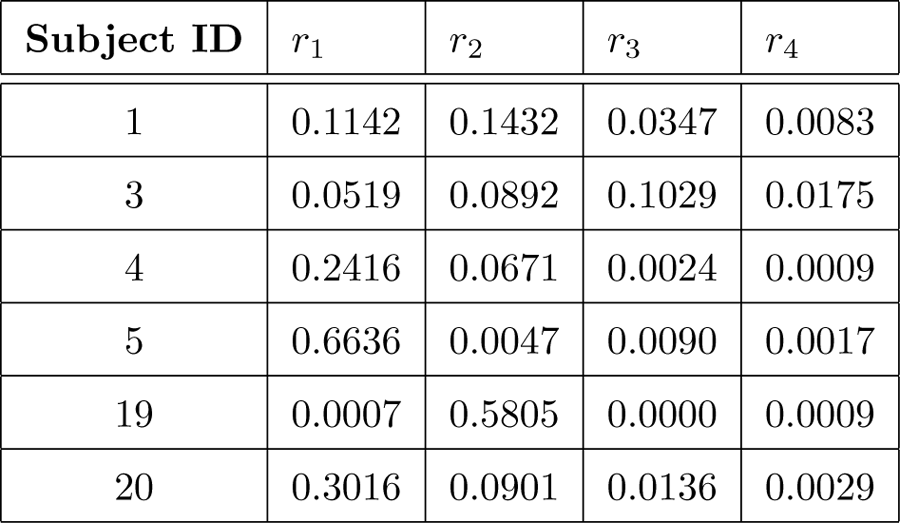
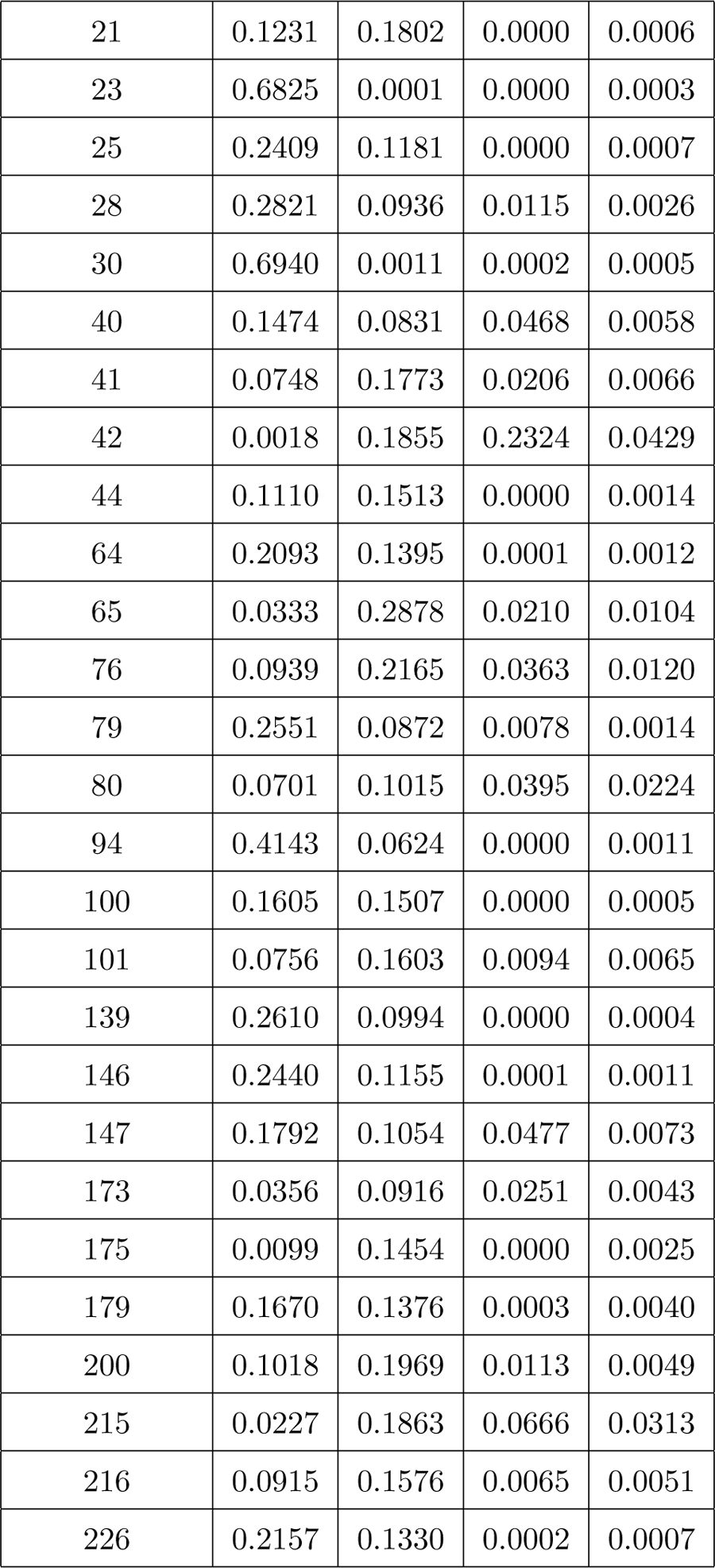
Best fit parameters *r*1*, r*2*, r*3*, r*4 per individual to 4 significant digits. Found via MATLAB’s Global Optimization.

Vaccine type for each subject:

**Table 3:** mRNA Vaccine Type per Subject.

| Subject | mRNA Vaccine |
| --- | --- |
| 1 | Moderna |
| 3 | Pfizer |
| 4 | Pfizer |
| 5 | Moderna |
| 19 | Pfizer |
| 20 | Moderna |
| 21 | Moderna |
| 23 | Moderna |
| 25 | Moderna |
| 28 | Moderna |
| 30 | Moderna |
| 40 | Moderna |
| 41 | Pfizer |
| 42 | Moderna |
| 44 | Pfizer |
| 64 | Moderna |
| 65 | Moderna |
| 76 | Pfizer |
| 79 | Moderna |
| 80 | Moderna |
| 94 | Moderna |
| 100 | Moderna |
| 101 | Moderna |
| 139 | Moderna |
| 146 | Pfizer |
| 147 | Moderna |
| 173 | Moderna |
| 175 | Moderna |
| 179 | Moderna |
| 200 | Moderna |
| 215 | Pfizer |
| 216 | Pfizer |
| 226 | Pfizer |

### B. Precision of Method for Determining Subject NAb Levels in IU/mL

To quantify the precision of the method for determining NAb IU/mL based on the percent inhibition AditxtScore^™^ assay for neutralizing antibodies to SARS-CoV-2, we ran the precision on 4 subjects with 4 different IU/mL values, 6 assays per day, for 3 days. Figure 16 shows a log-fit to the data. Interpolated percent variation values, based on the log fit, are in Table 4, given at 50 IU/mL intervals.

**Figure 16:**
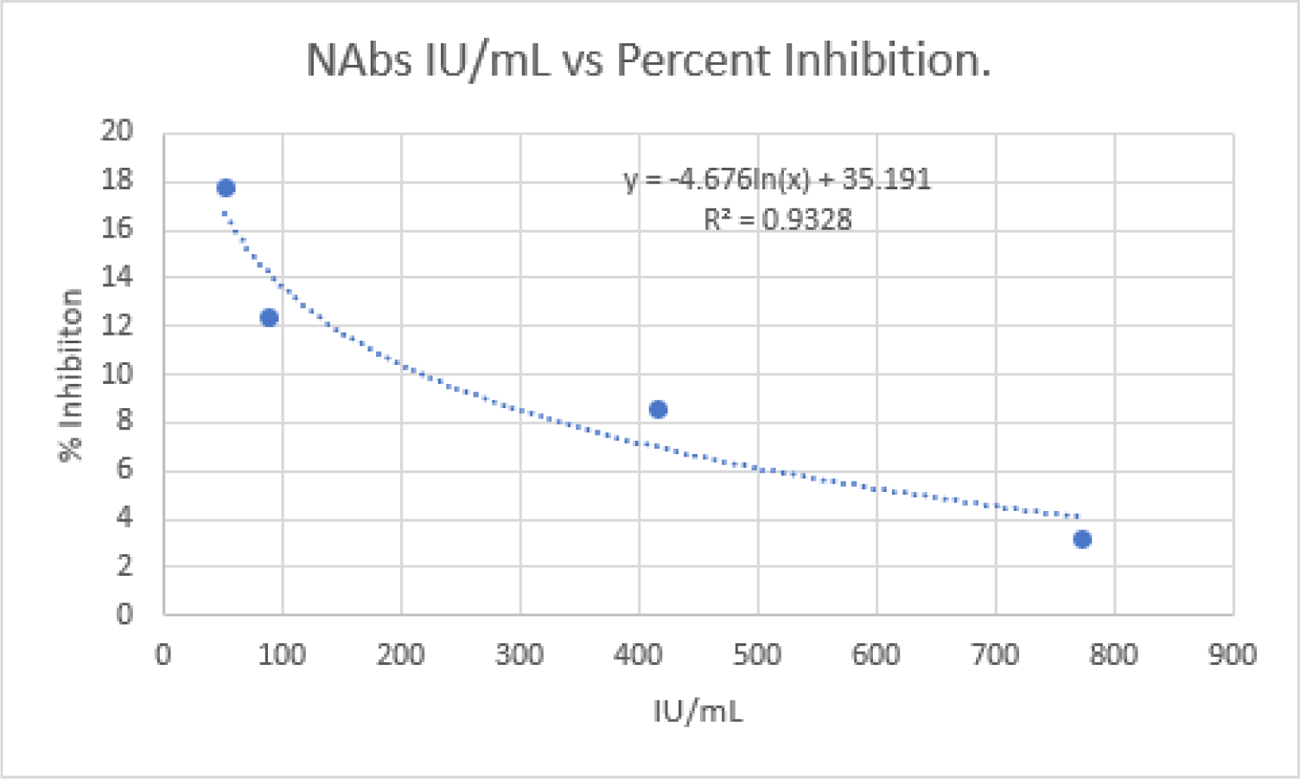
Approximate relationship between % inhibition as measured by the AditxtScore^™^ test for neutralizing antibodies to SARS-CoV-2 and NAbs in IU/mL.

**Table 4:** Precision of method to determine AditxtScore^™^ for neutralizing antibodies (NAb) to SARS-CoV-2, given in 50 IU/mL intervals. Precision is inversely proportional to IU/mL.

| Avg. IU/mL | Avg. %CV |
| --- | --- |
| 50 | 16.9 |
| 100 | 13.7 |
| 150 | 11.8 |
| 200 | 10.4 |
| 250 | 9.4 |
| 300 | 8.5 |
| 350 | 7.8 |
| 400 | 7.2 |
| 450 | 6.6 |
| 500 | 6.1 |
| 550 | 5.7 |
| 600 | 5.3 |
| 650 | 4.9 |
| 700 | 4.6 |
| 750 | 4.2 |
| 800 | 3.9 |

### C. Simulation Plots

Shown are simulations using best-fit global optimization parameters as well as plots of the MCMC predictive envelopes for each subject. Also shown for each subject are the MCMC analytics: the MCMC parameter chains, corner plots, and histograms generated by the mcmcplot command from the mcmcstat package [25].

**Figure 17:**
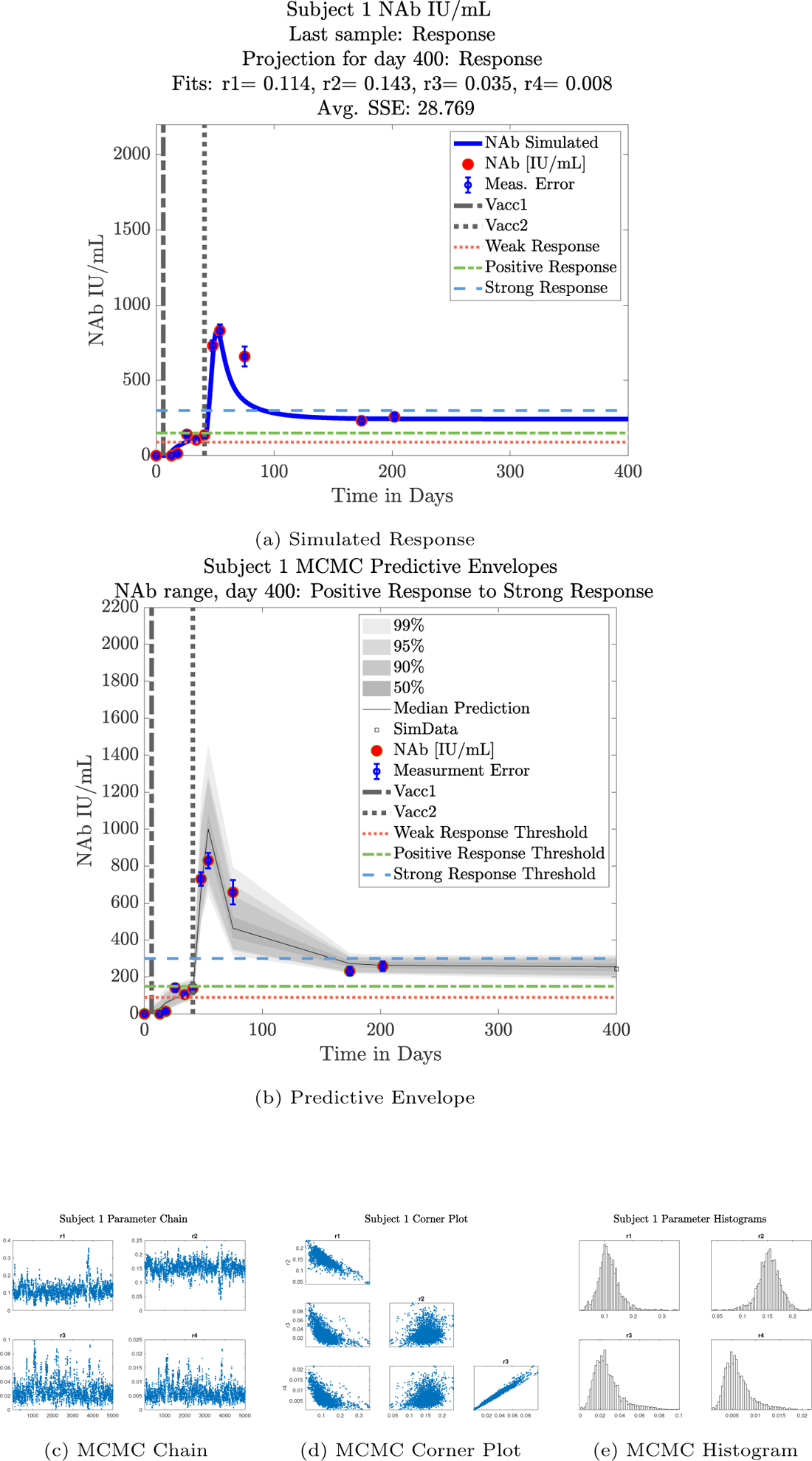
Subject 1

**Figure 18:**
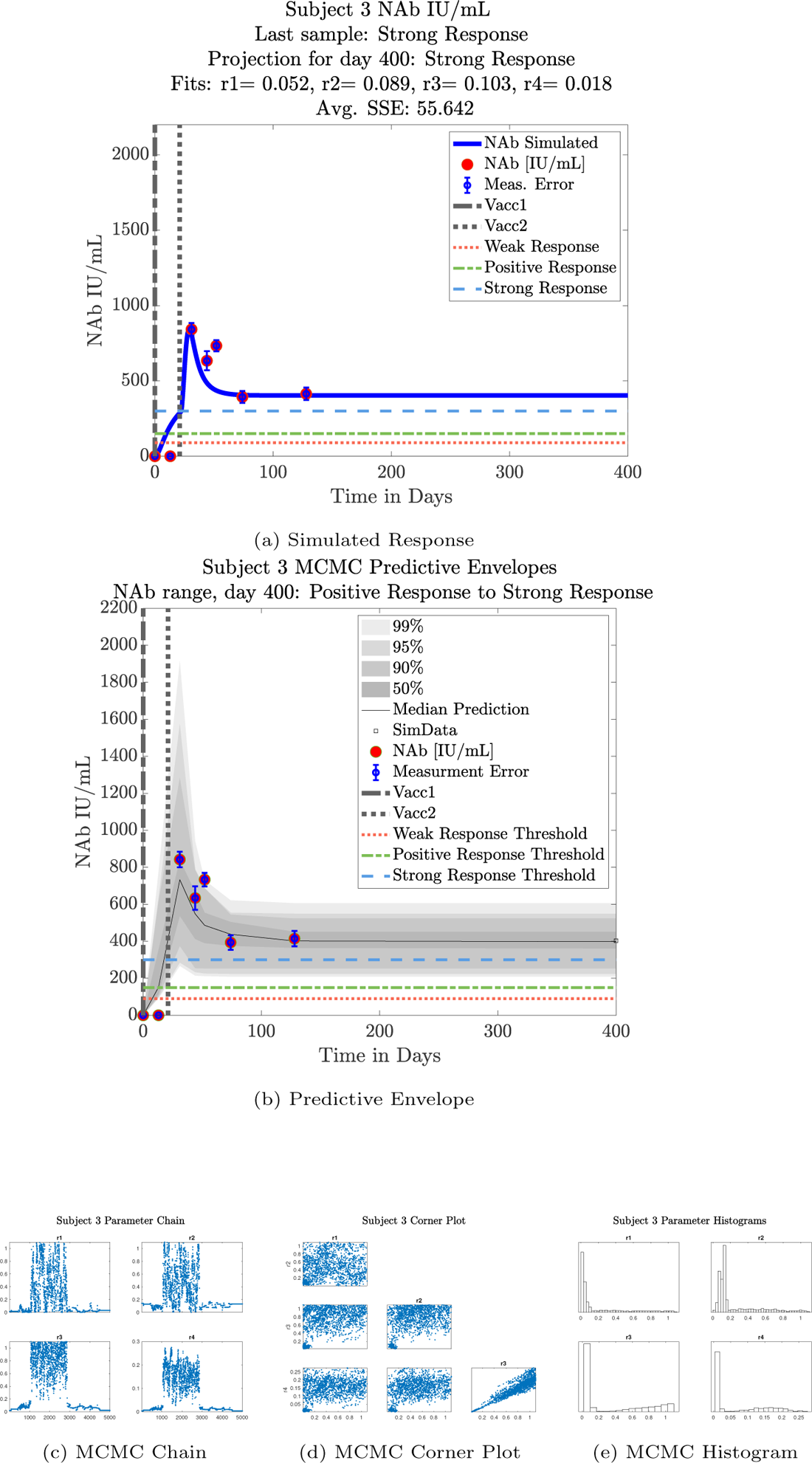
Subject 3

**Figure 19:**
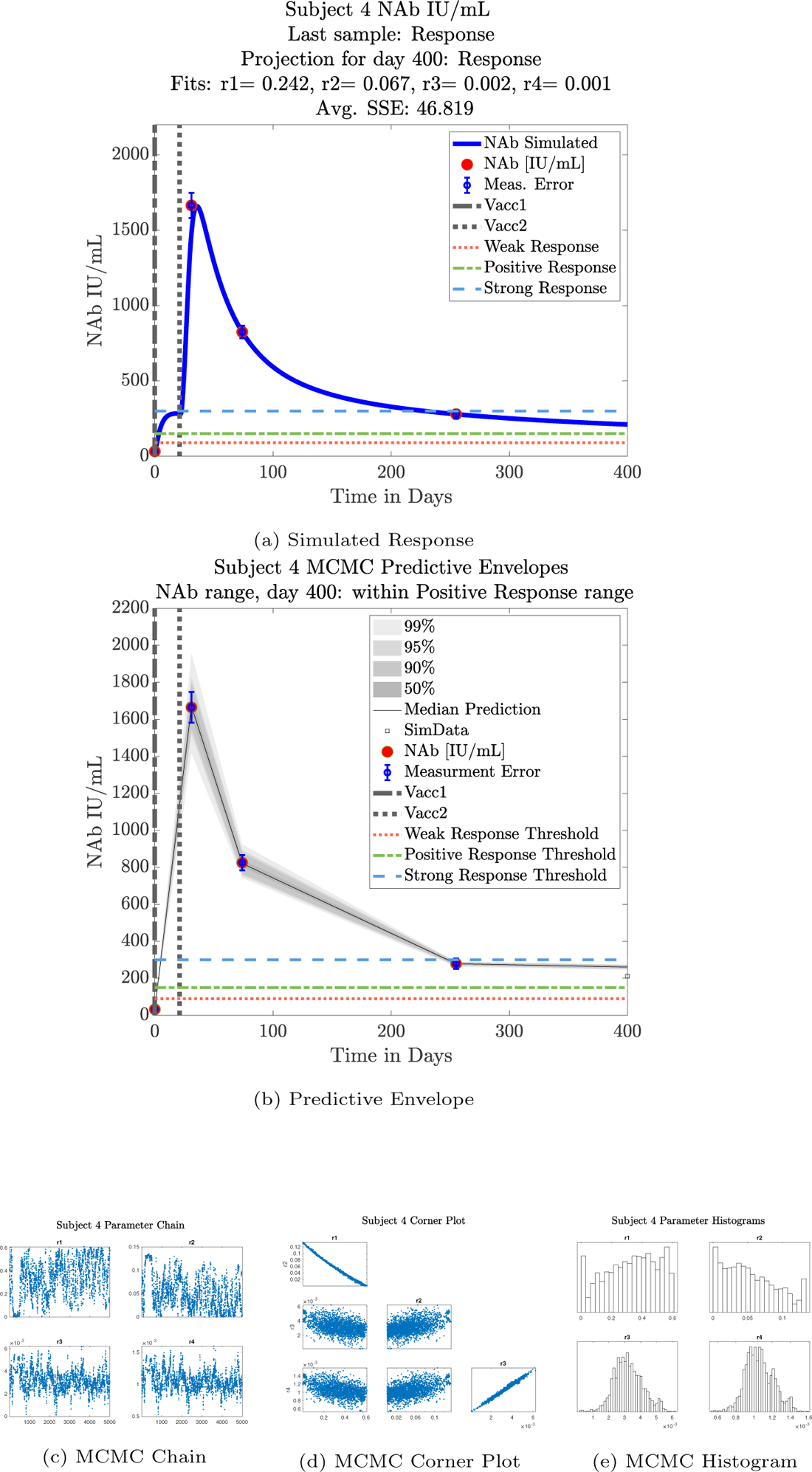
Subject 4

**Figure 20:**
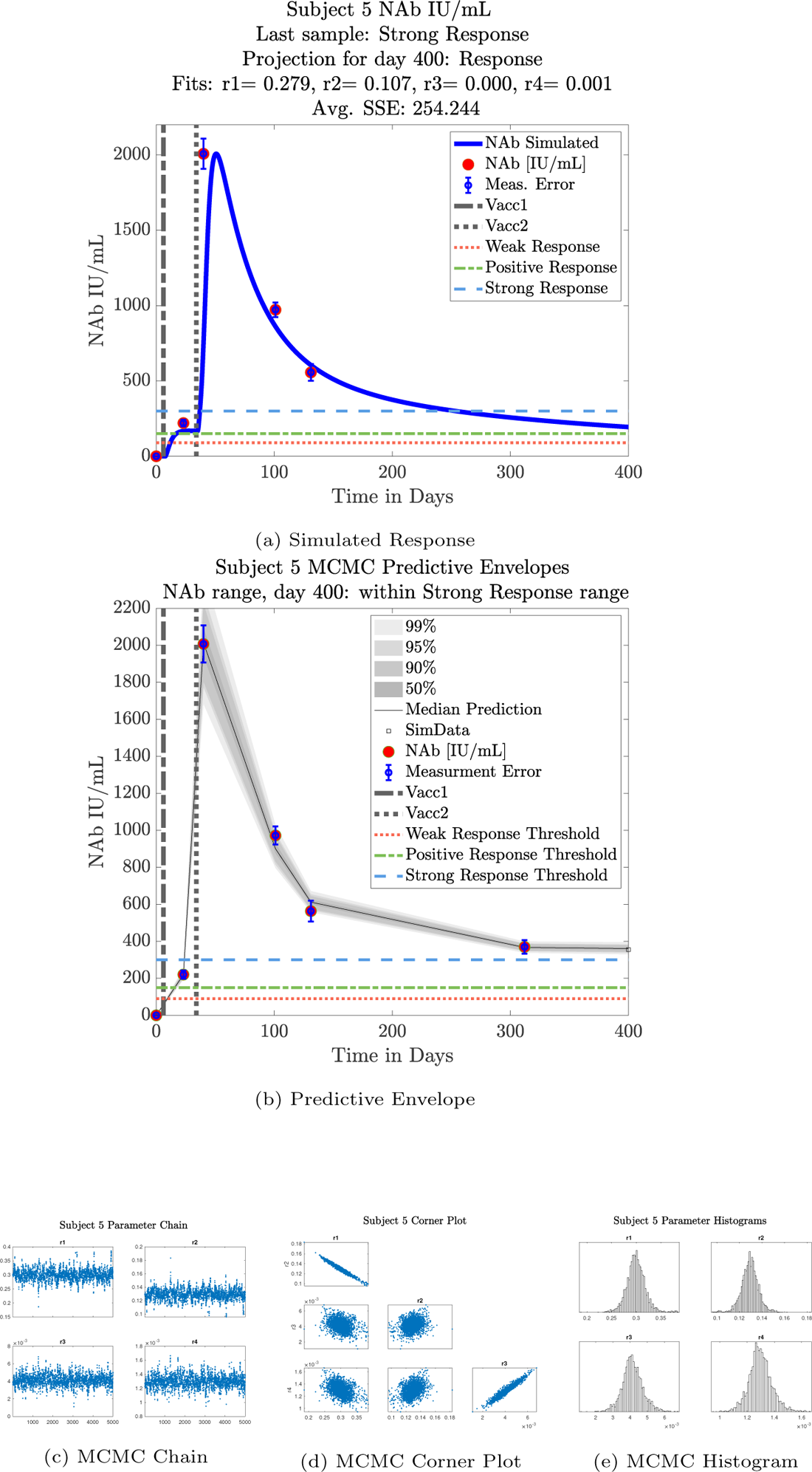
Subject 5

**Figure 21:**
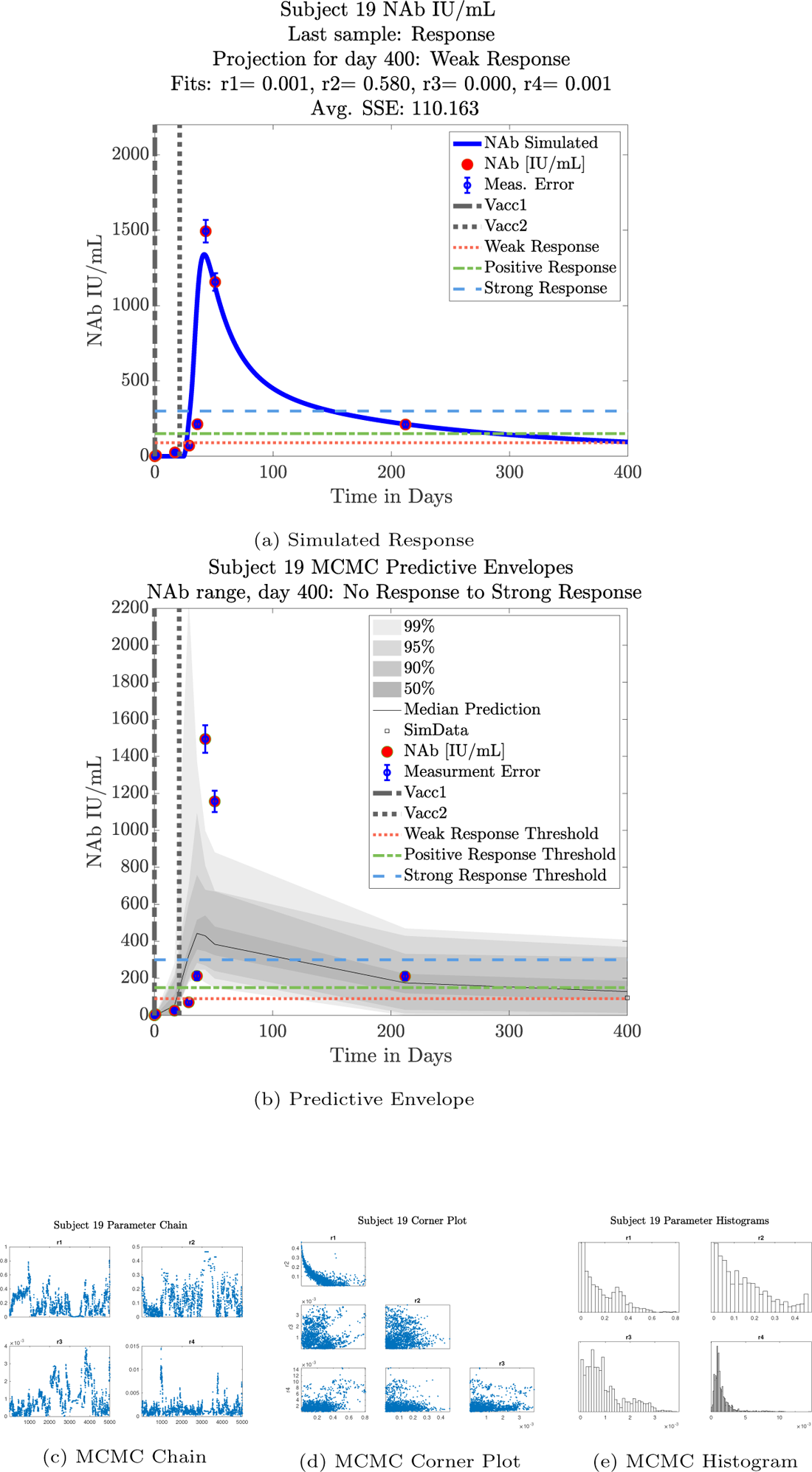
Subject 19

**Figure 22:**
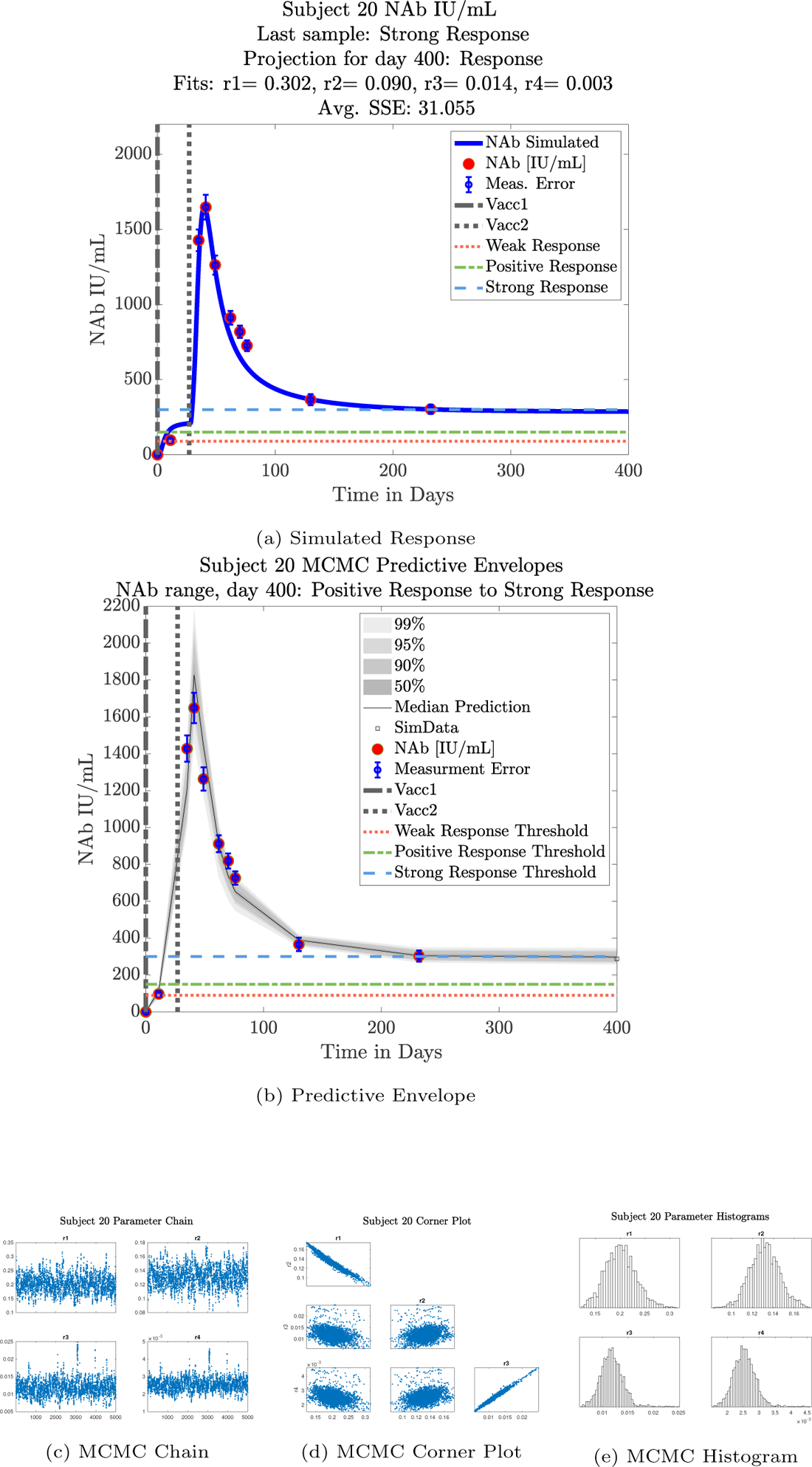
Subject 20

**Figure 23:**
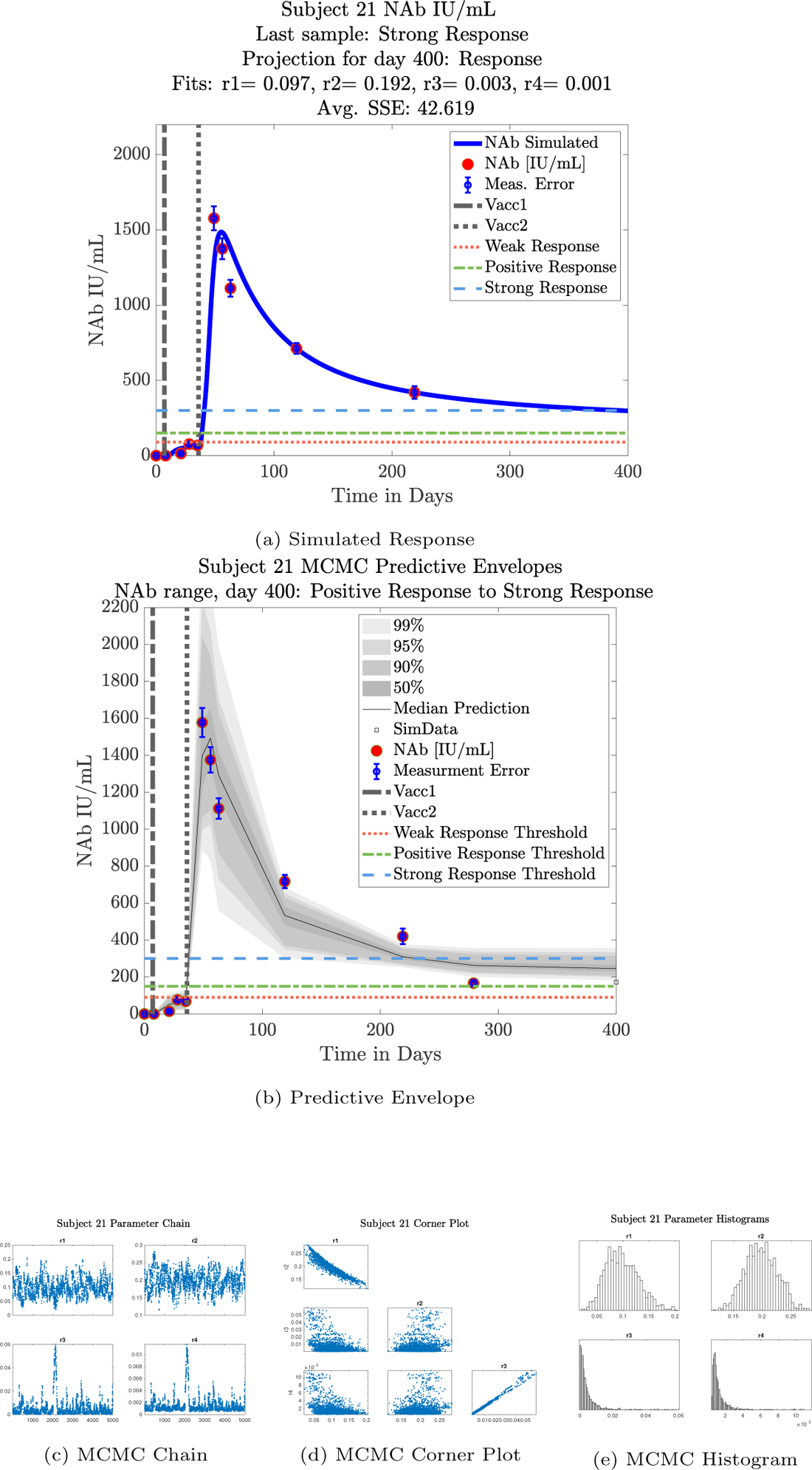
Subject 21

**Figure 24:**
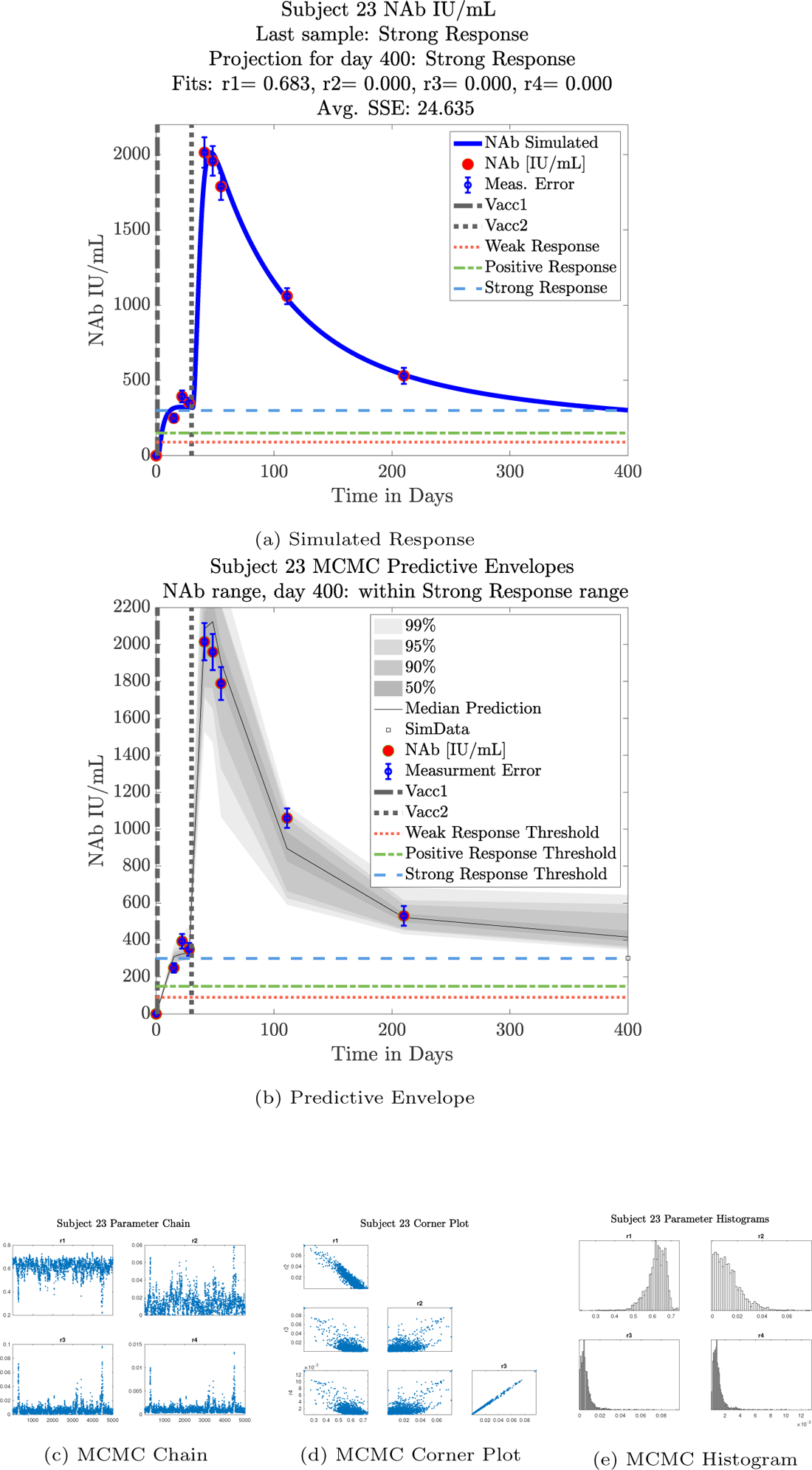
Subject 23

**Figure 25:**
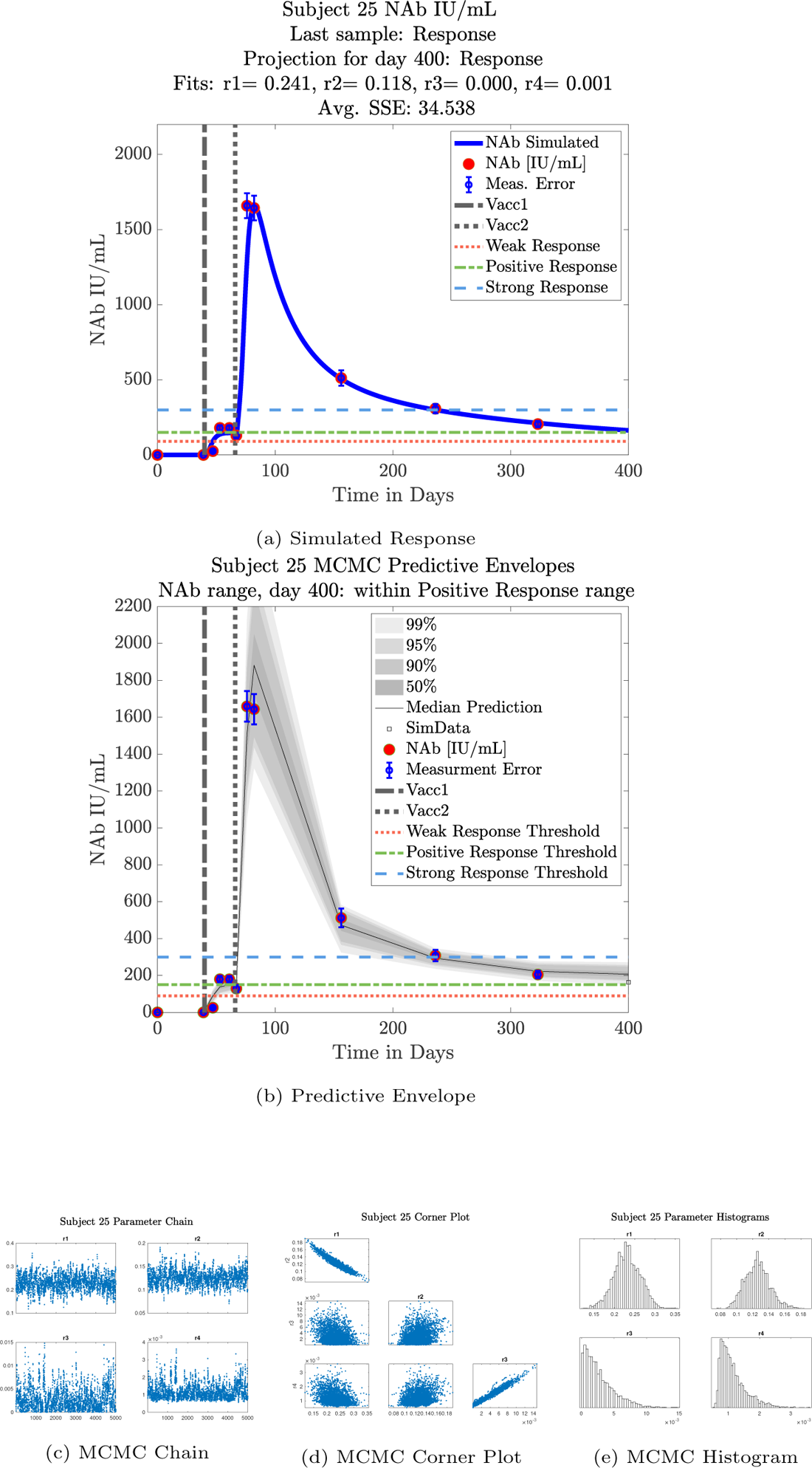
Subject 25

**Figure 26:**
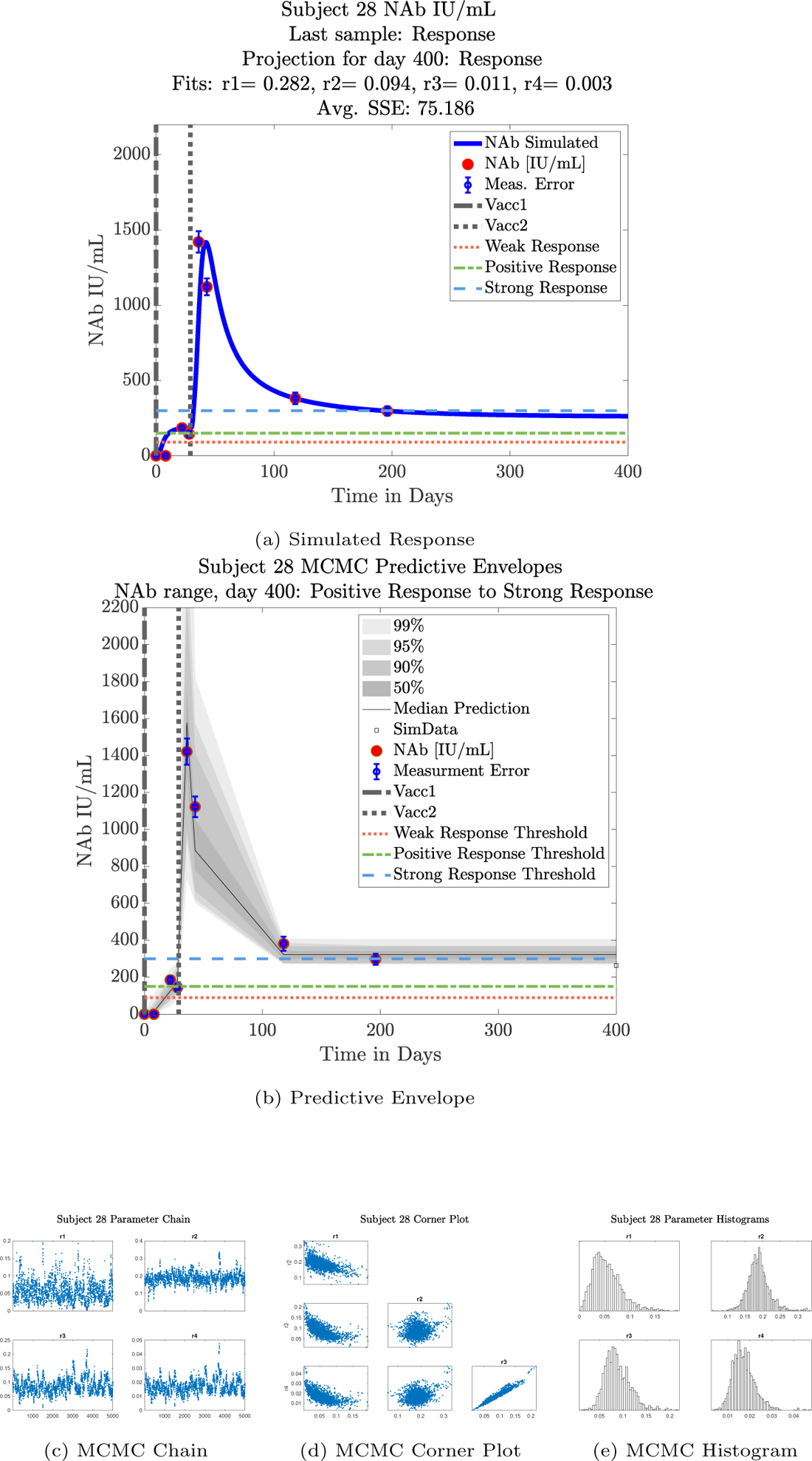
Subject 28

**Figure 27:**
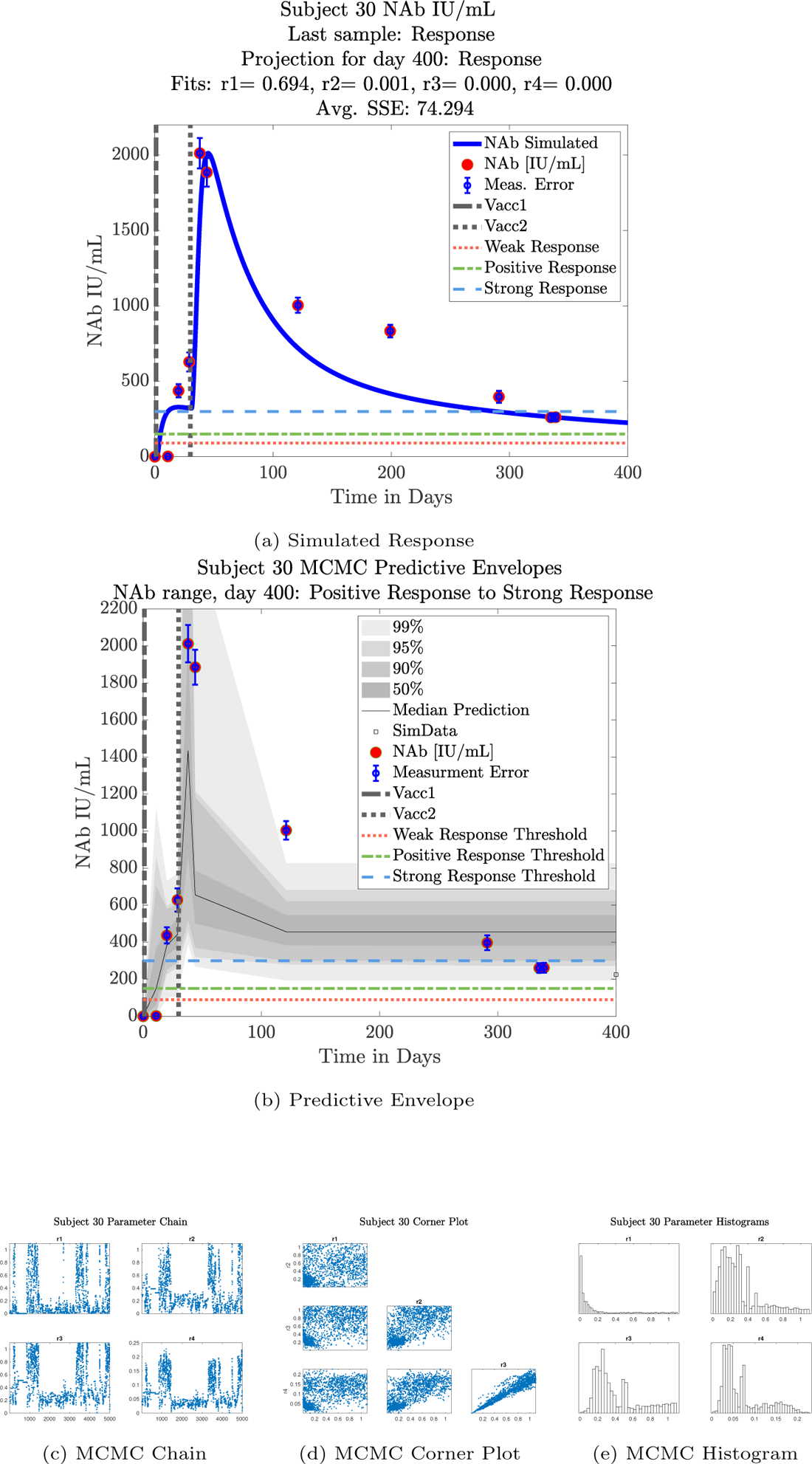
Subject 30

**Figure 28:**
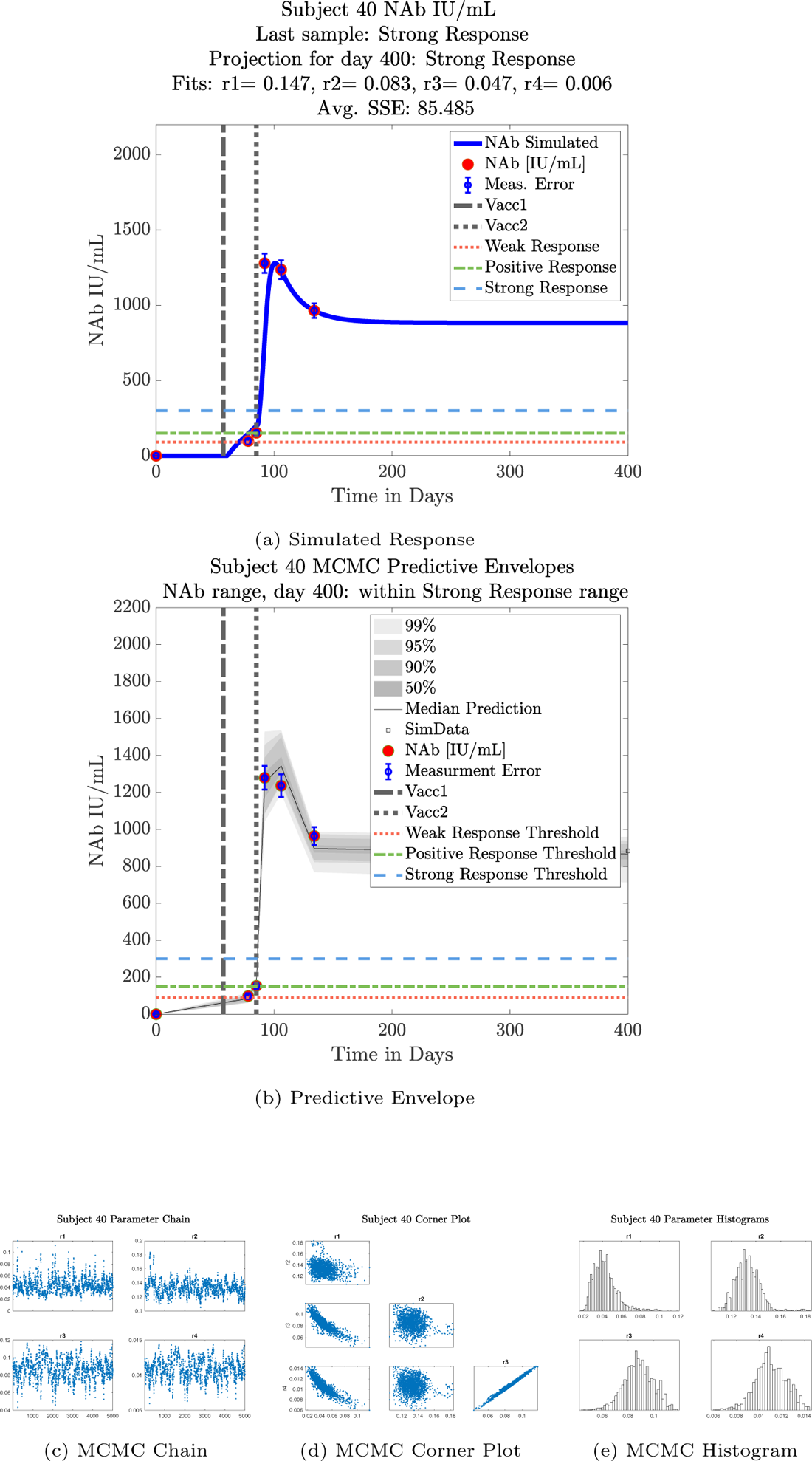
Subject 40

**Figure 29:**
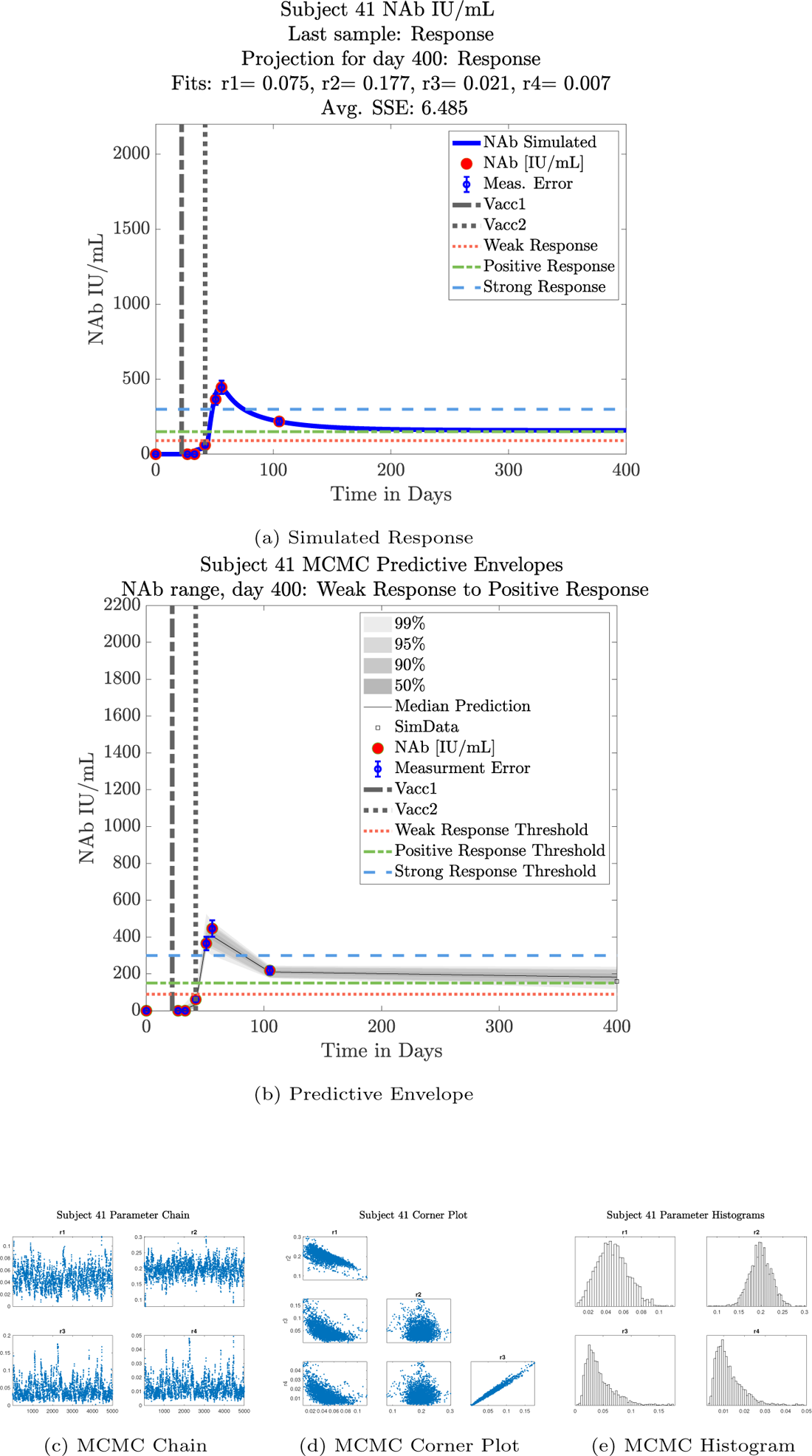
Subject 41

**Figure 30:**
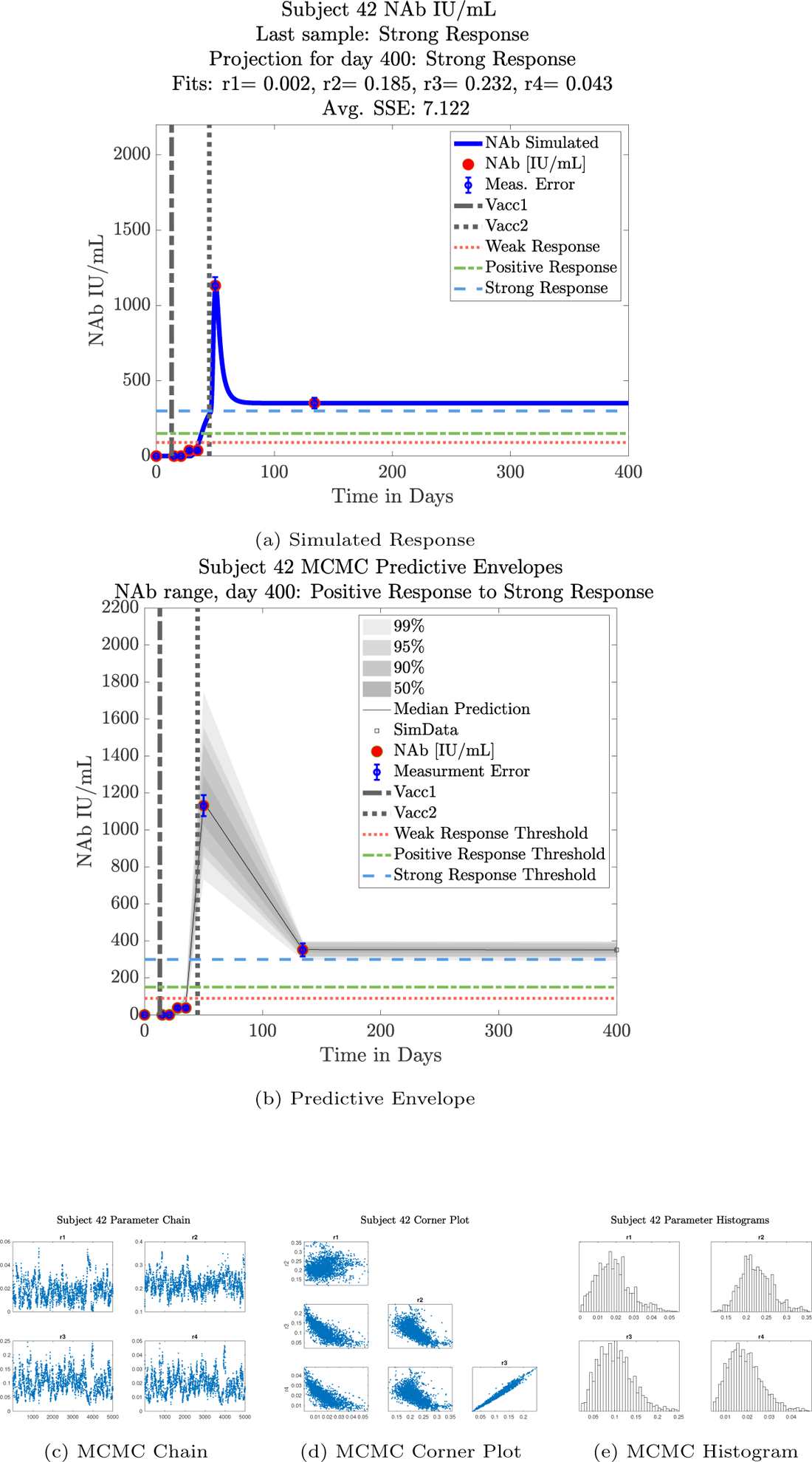
Subject 42

**Figure 31:**
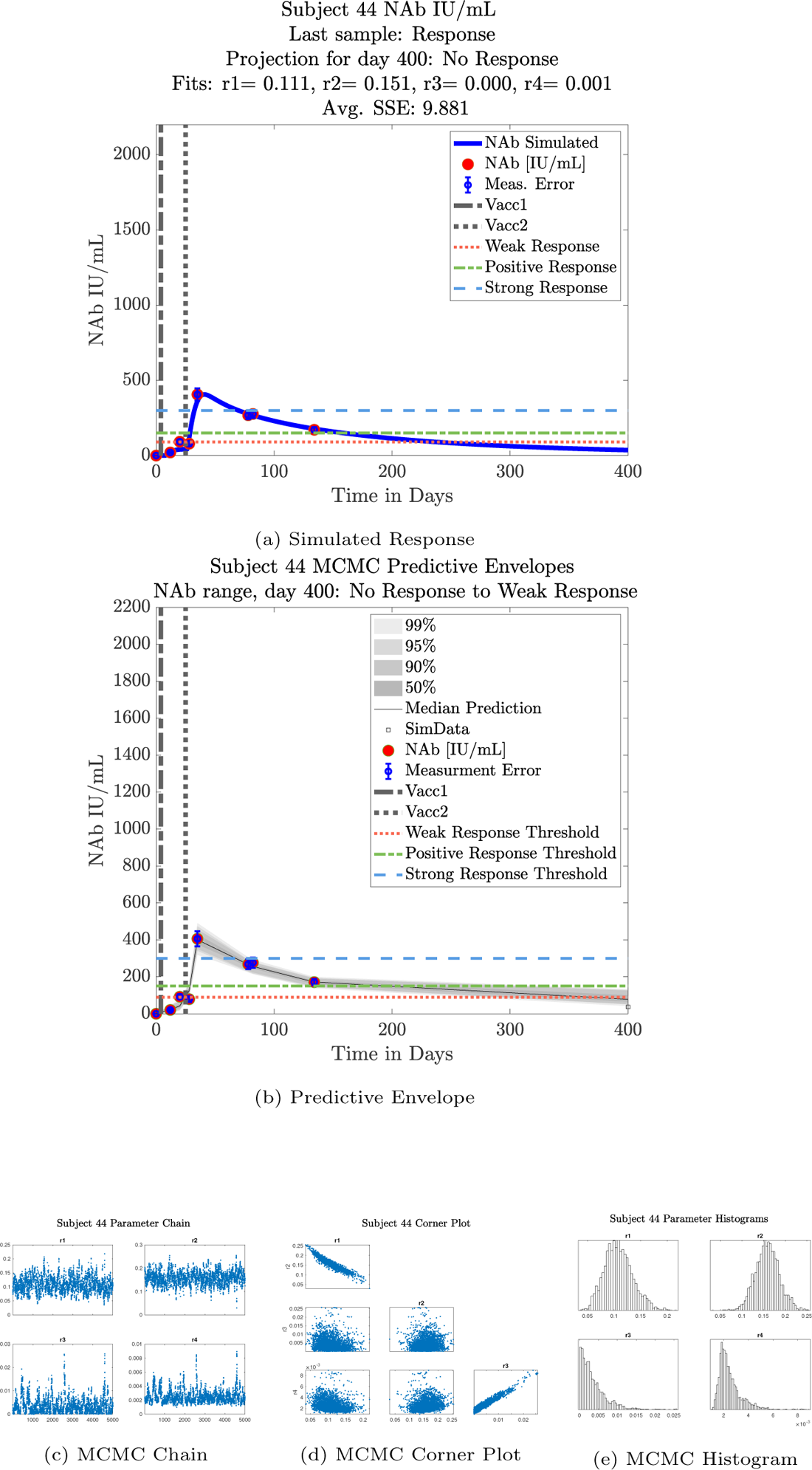
Subject 44

**Figure 32:**
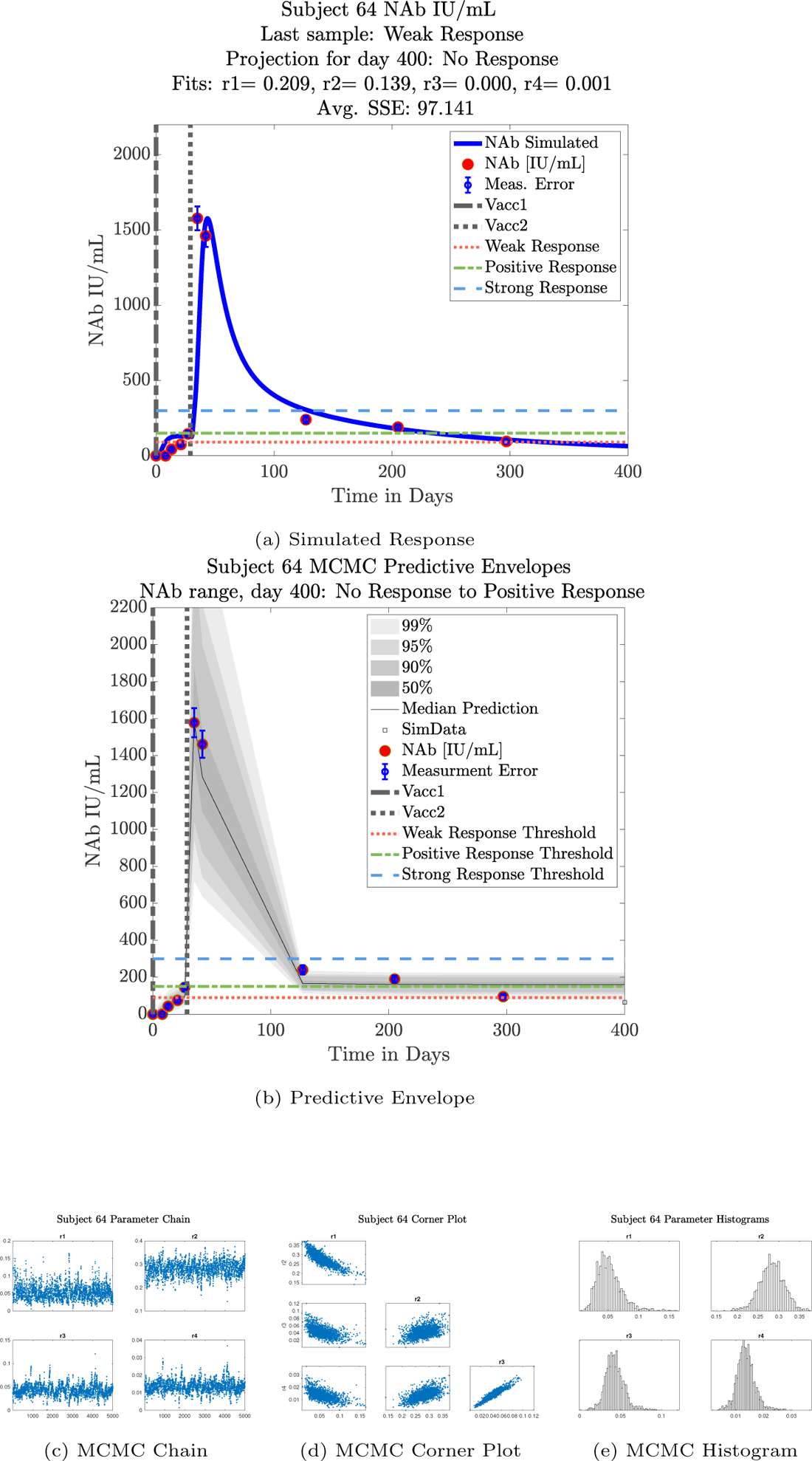
Subject 64

**Figure 33:**
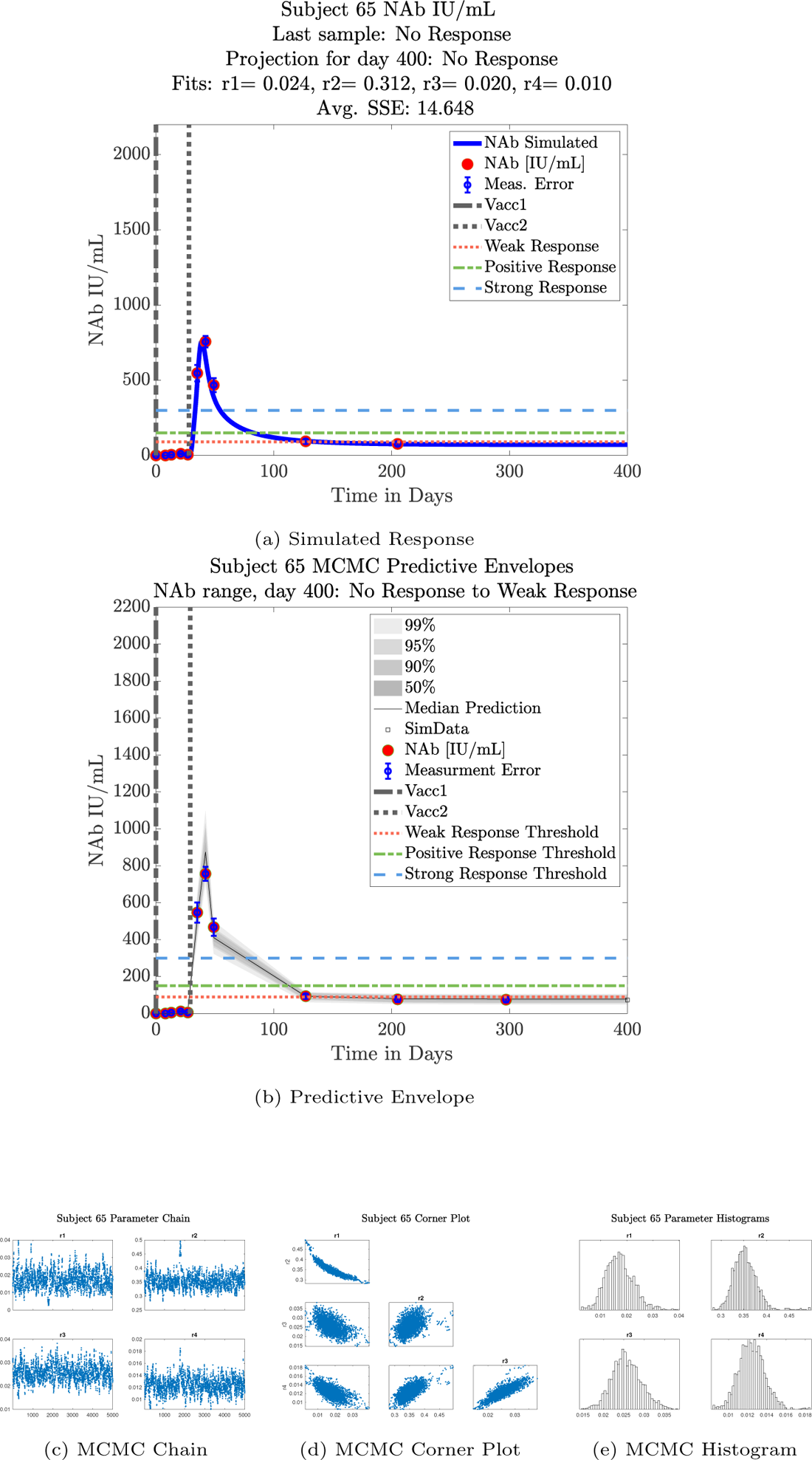
Subject 65

**Figure 34:**
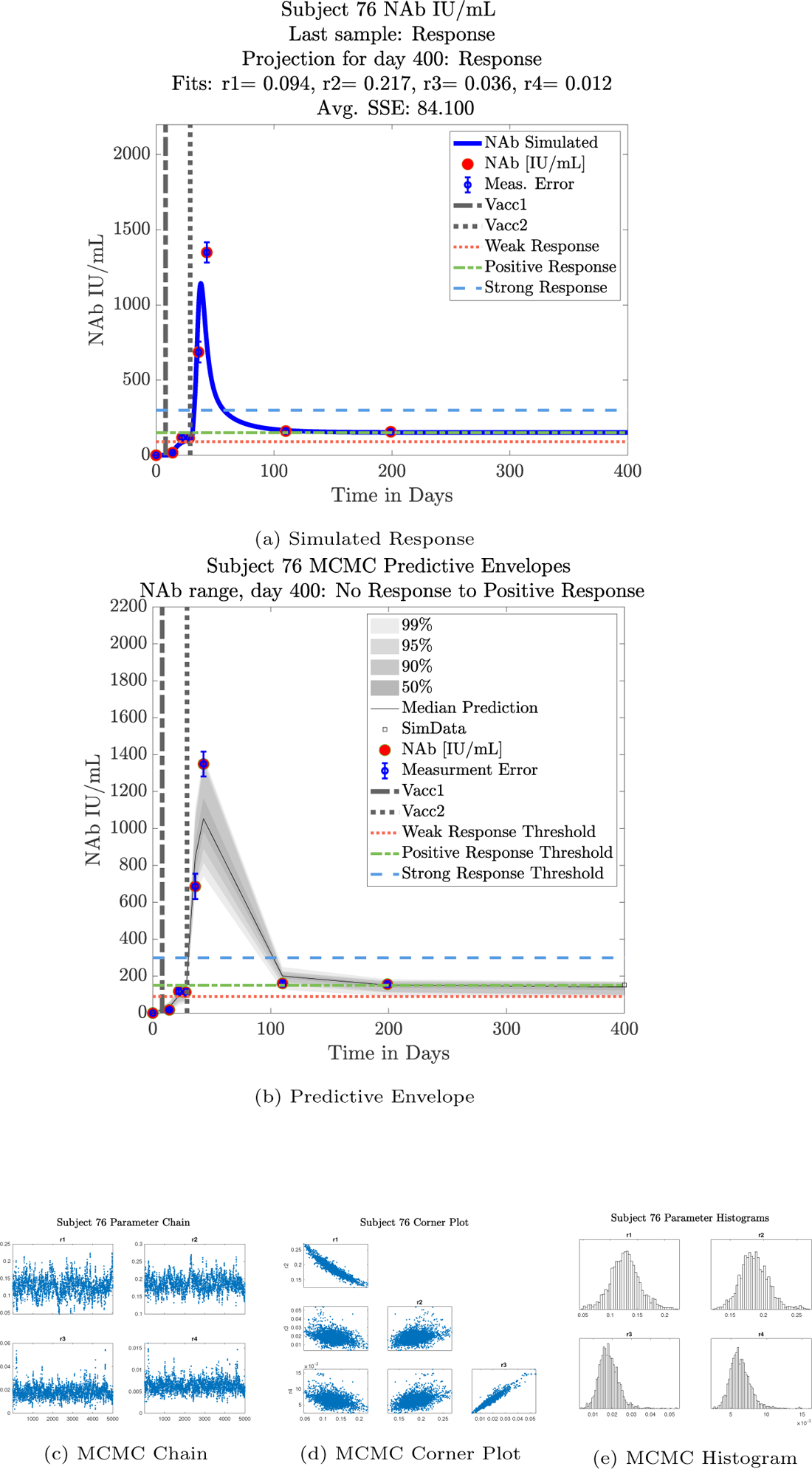
Subject 76

**Figure 35:**
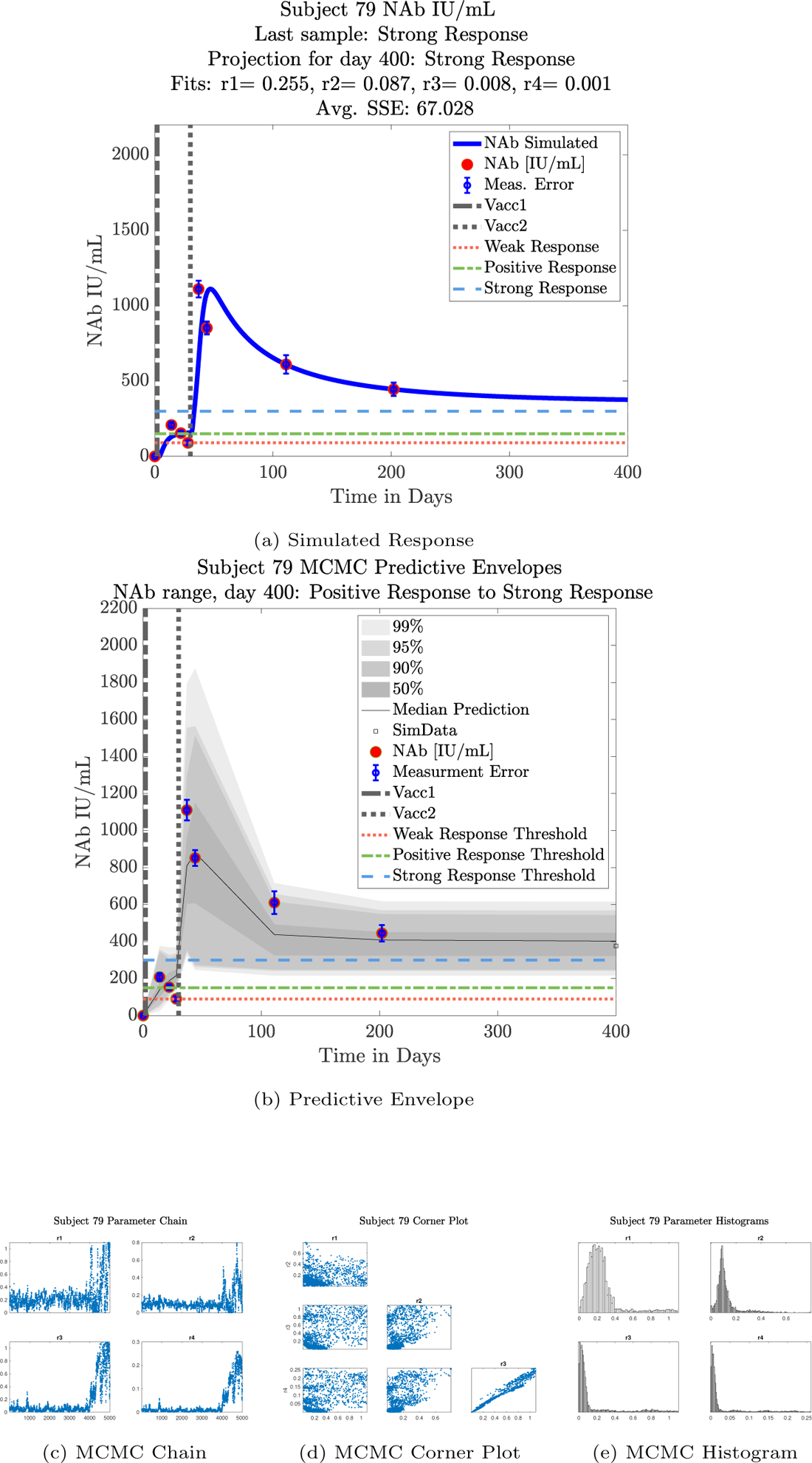
Subject 79

**Figure 36:**
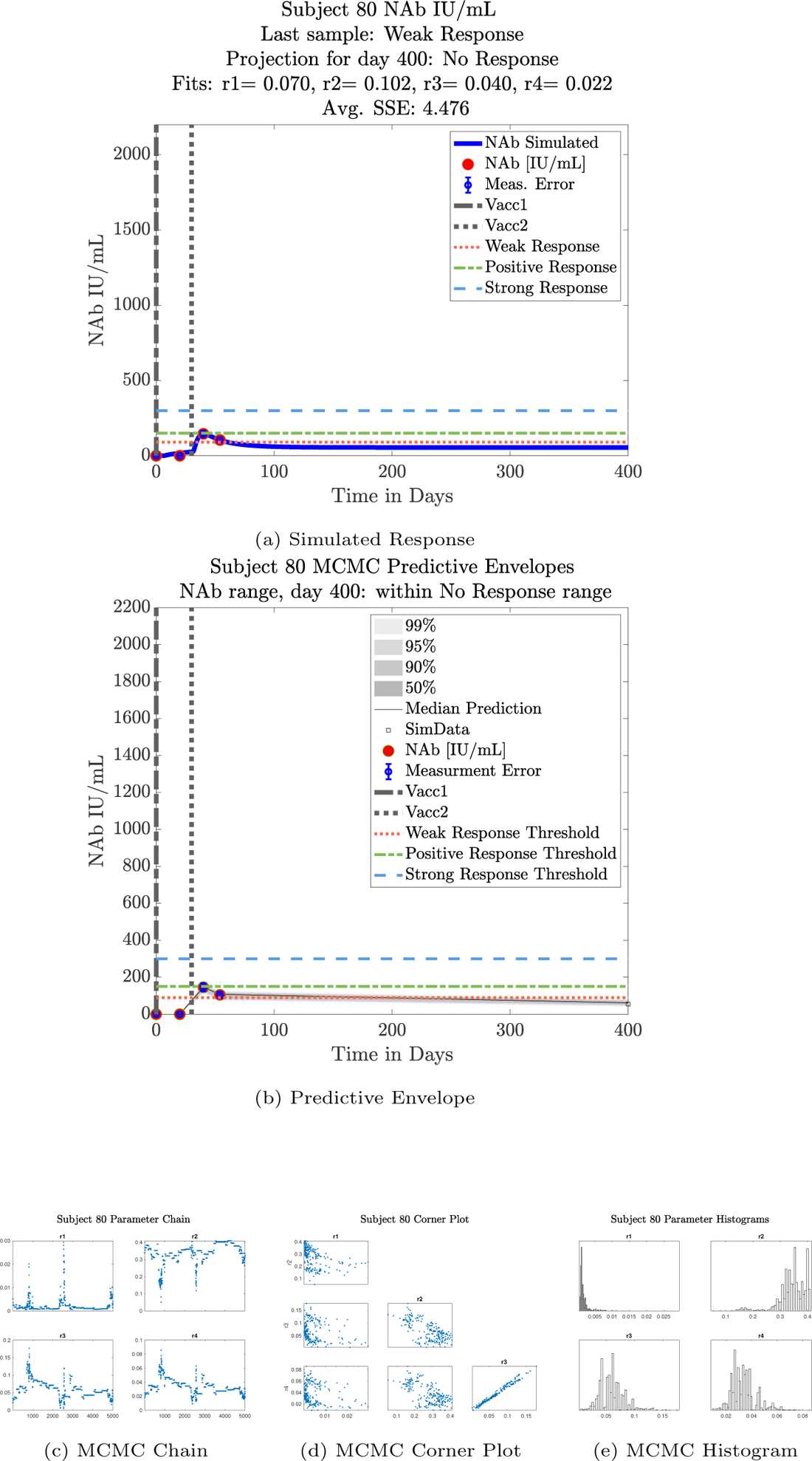
Subject 80

**Figure 37:**
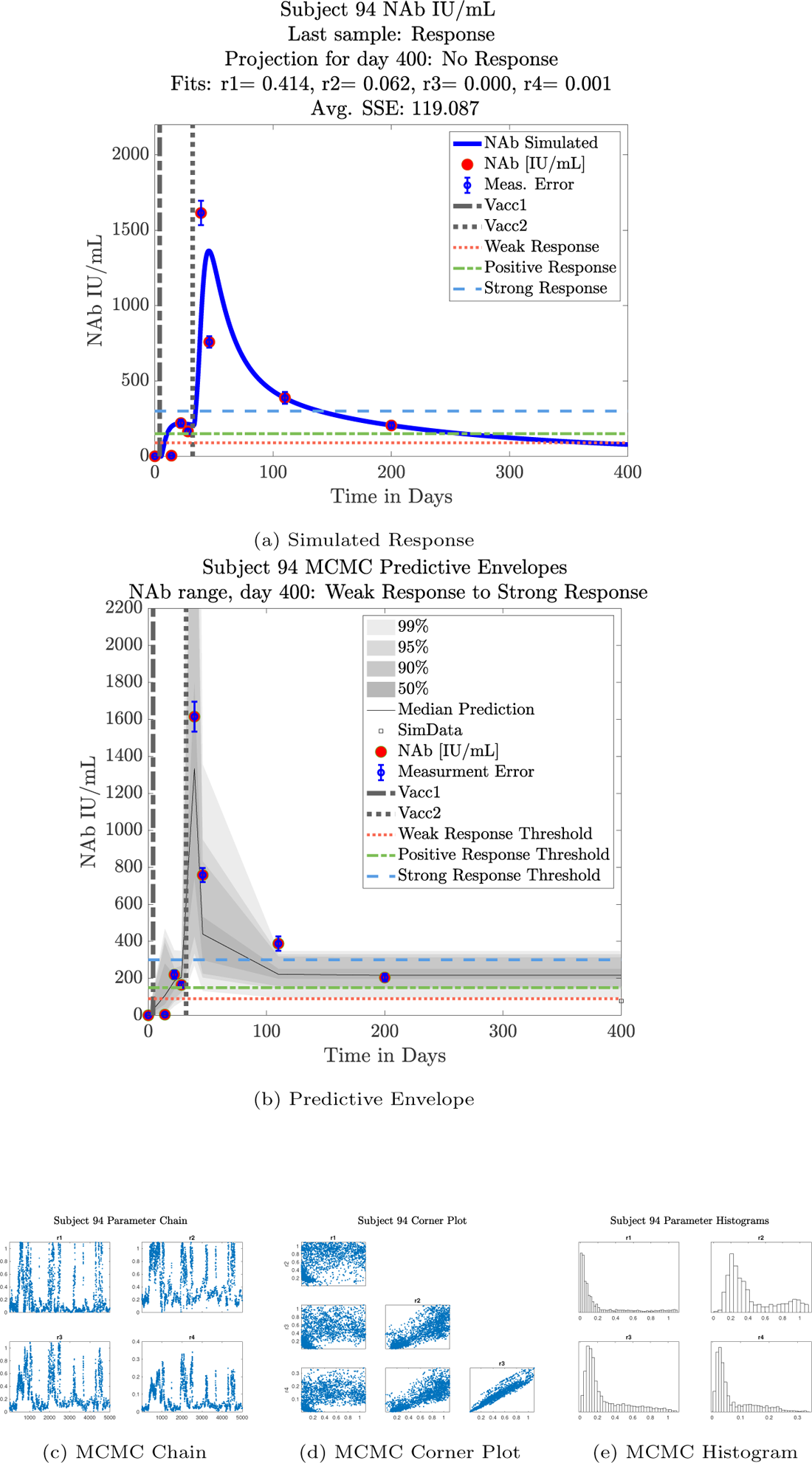
Subject 94

**Figure 38:**
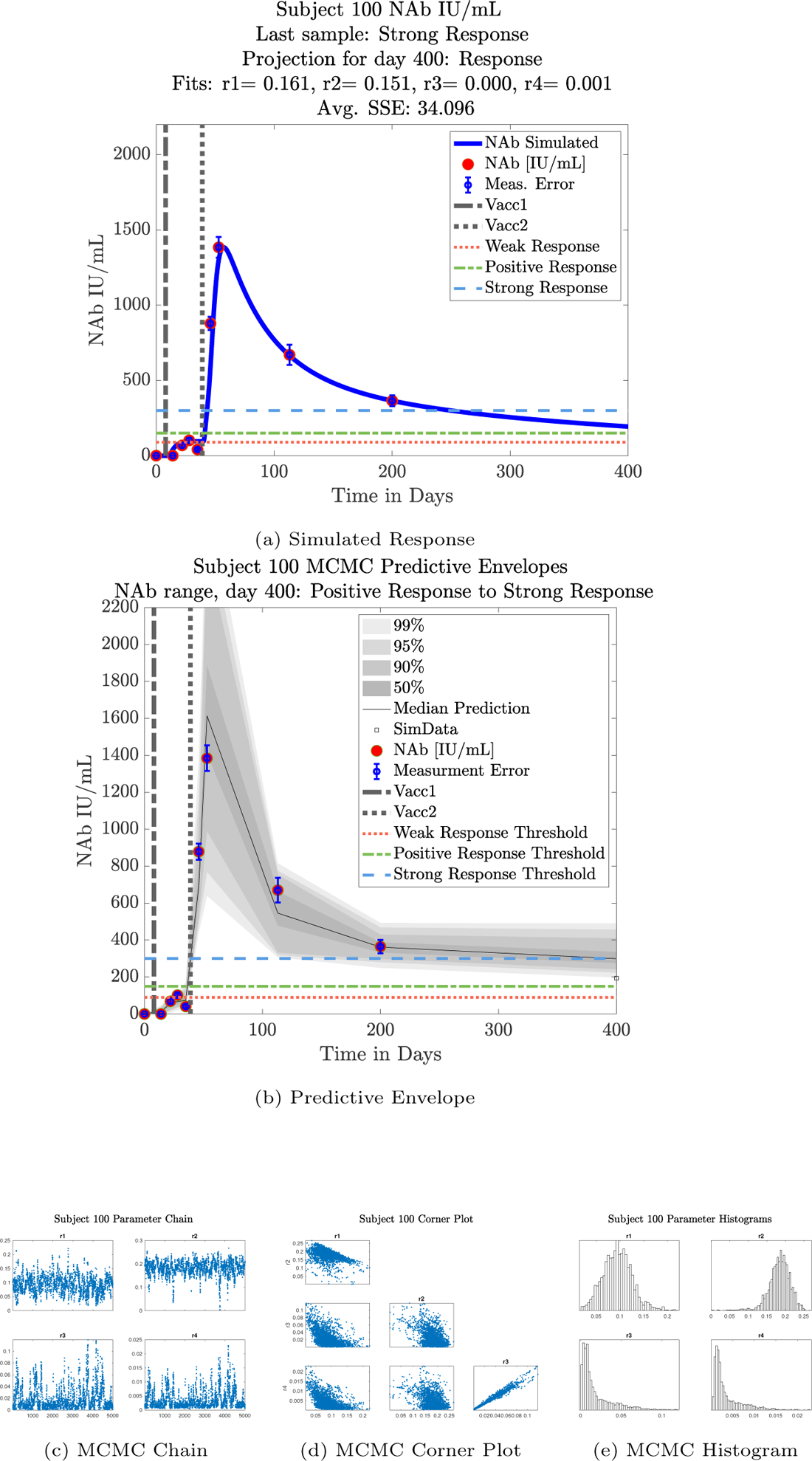
Subject 100

**Figure 39:**
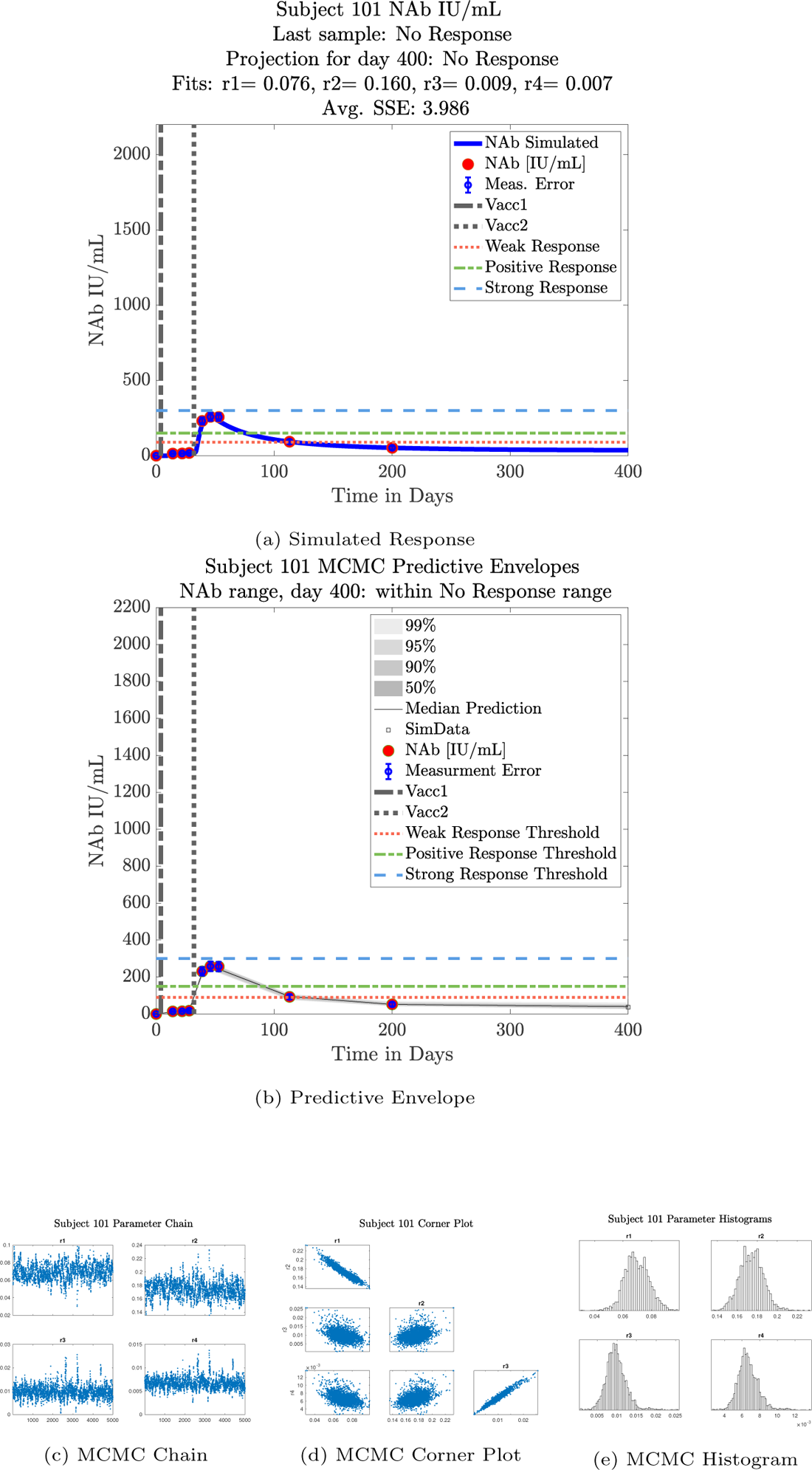
Subject 101

**Figure 40:**
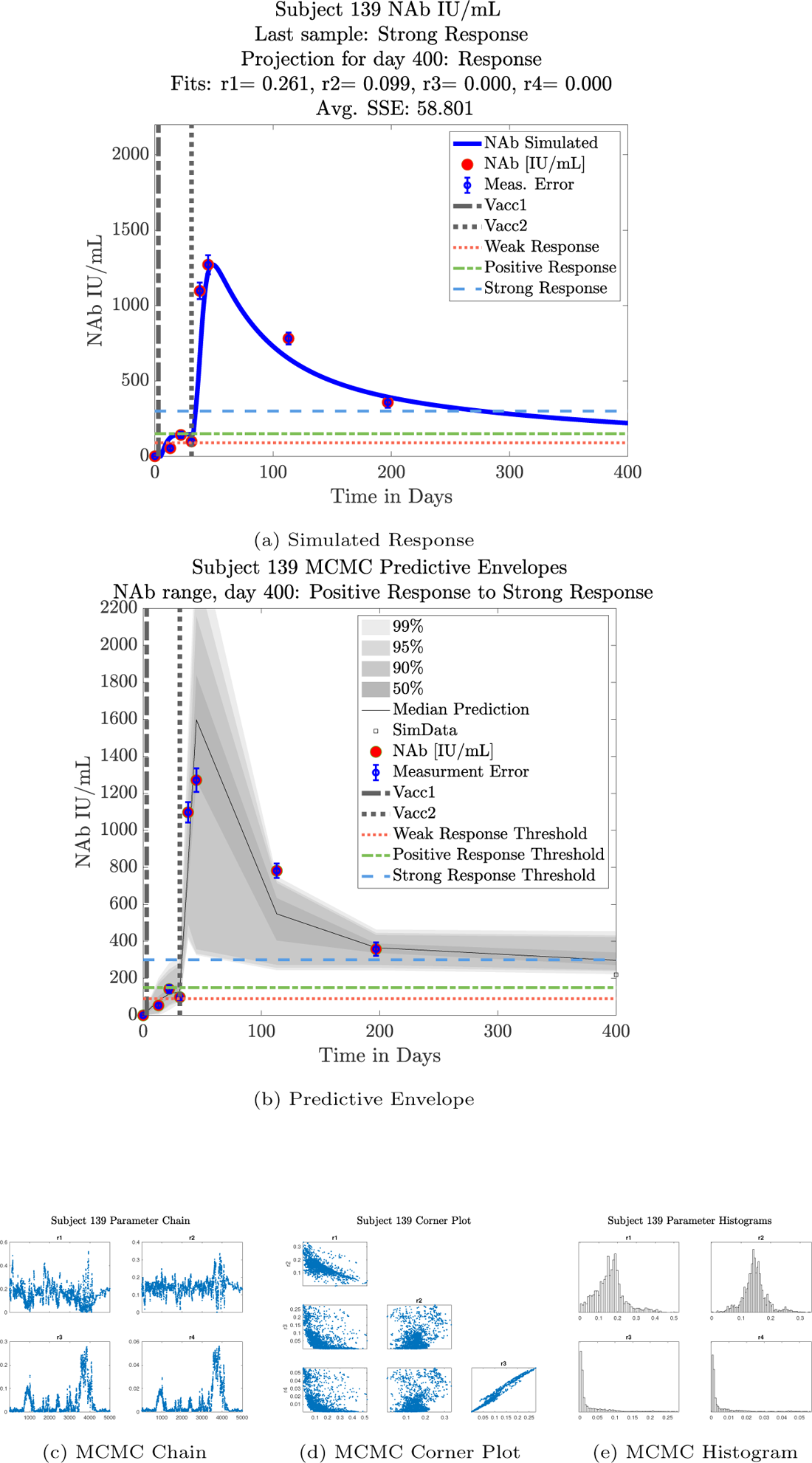
Subject 139

**Figure 41:**
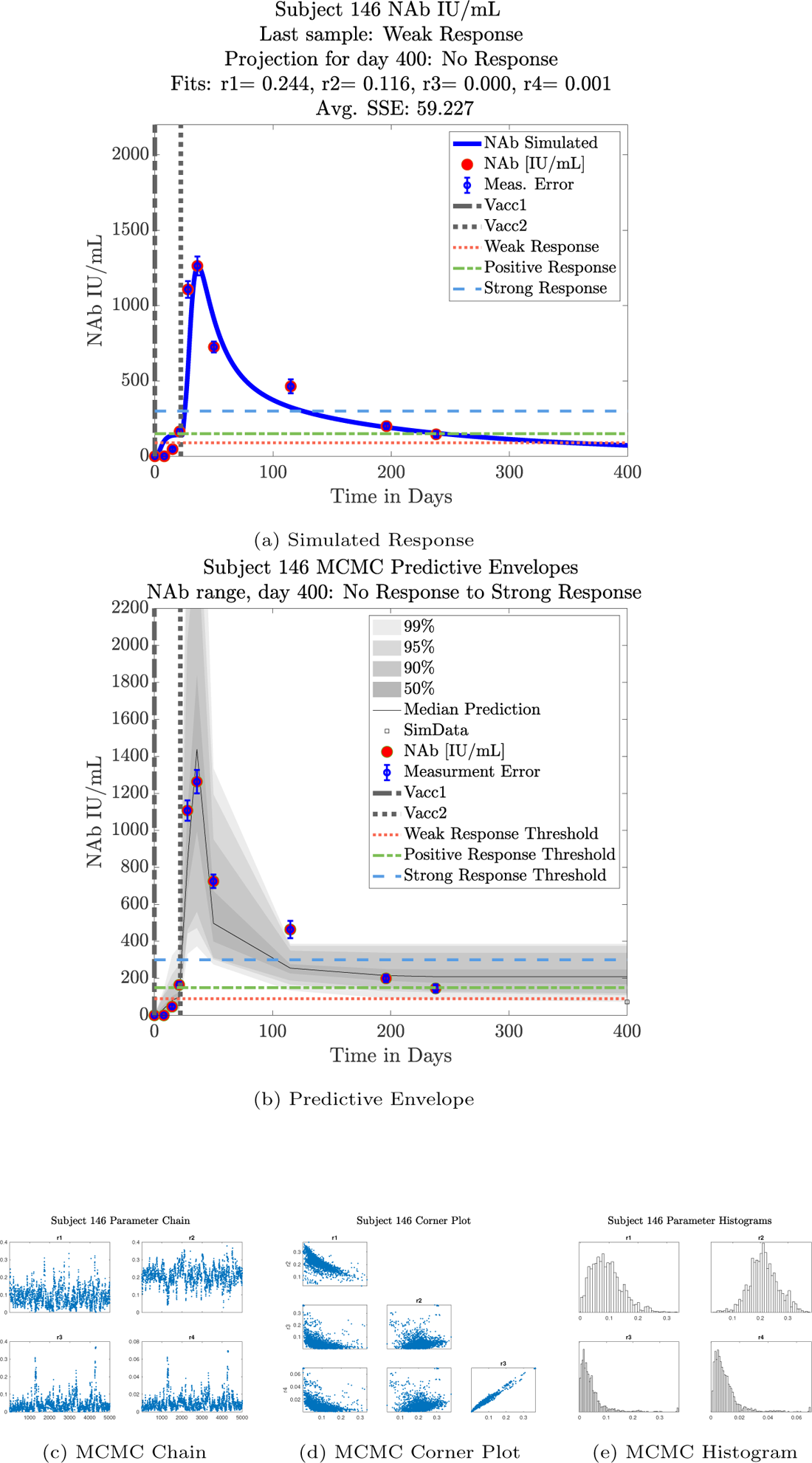
Subject 146

**Figure 42:**
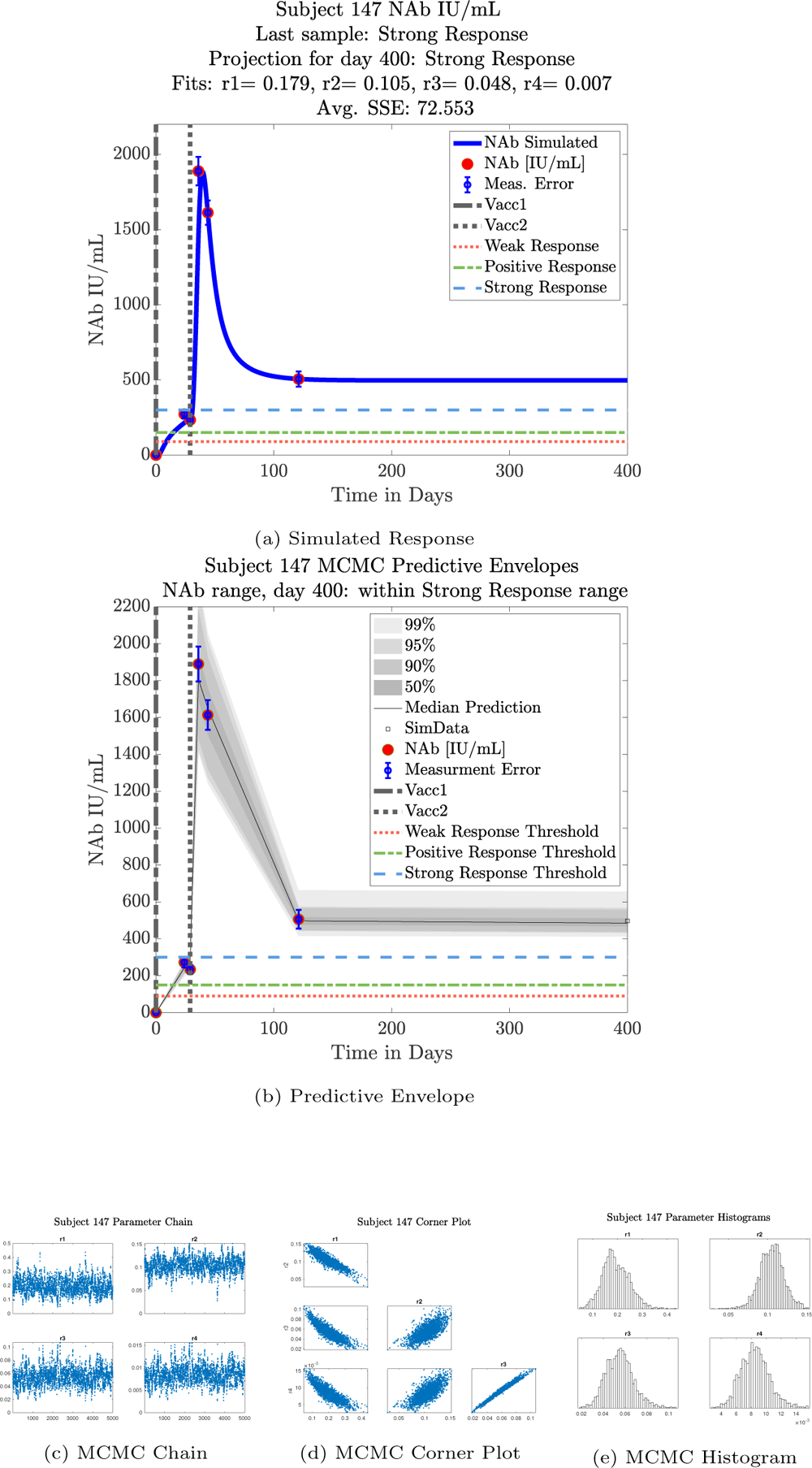
Subject 147

**Figure 43:**
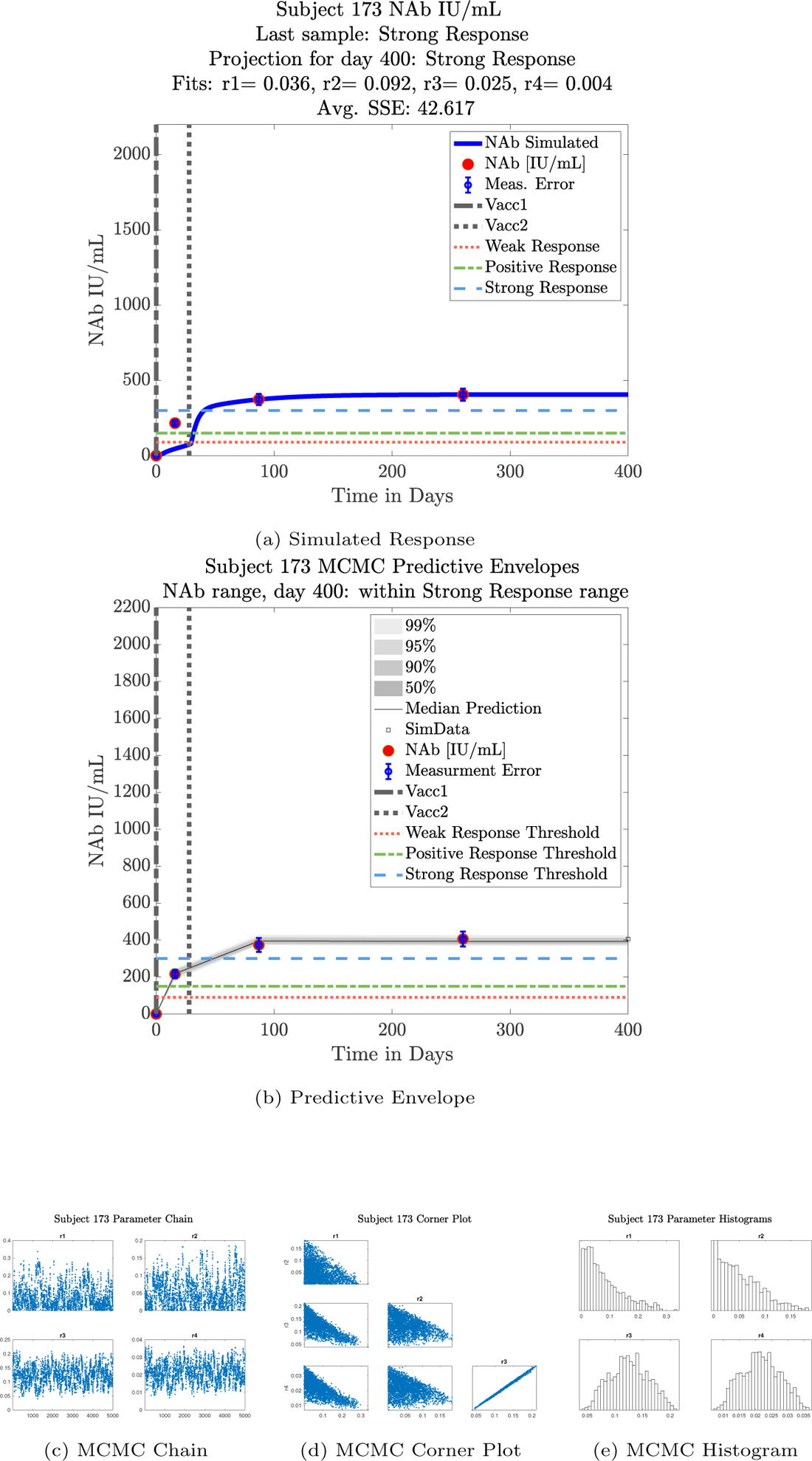
Subject 173

**Figure 44:**
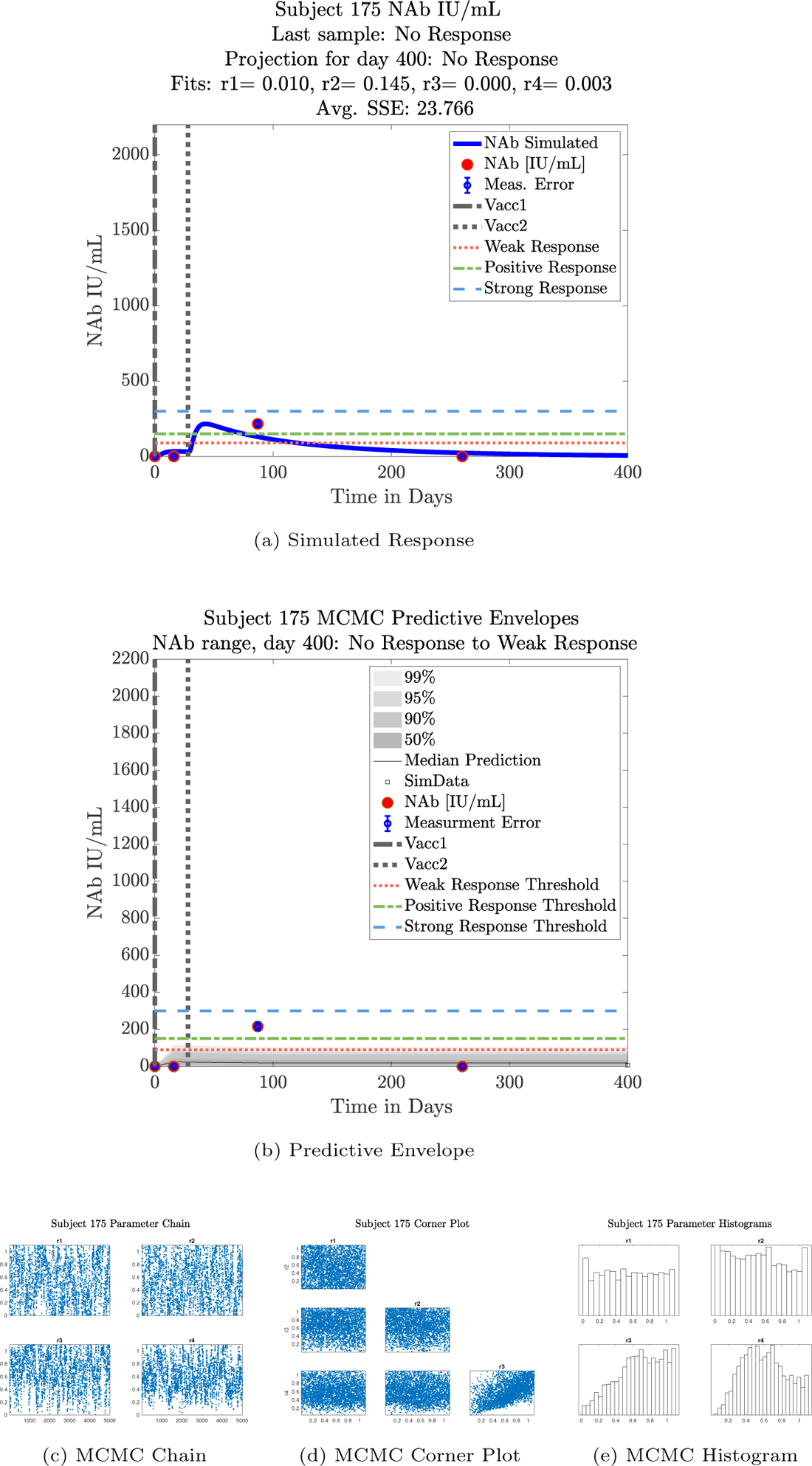
Subject 175

**Figure 45:**
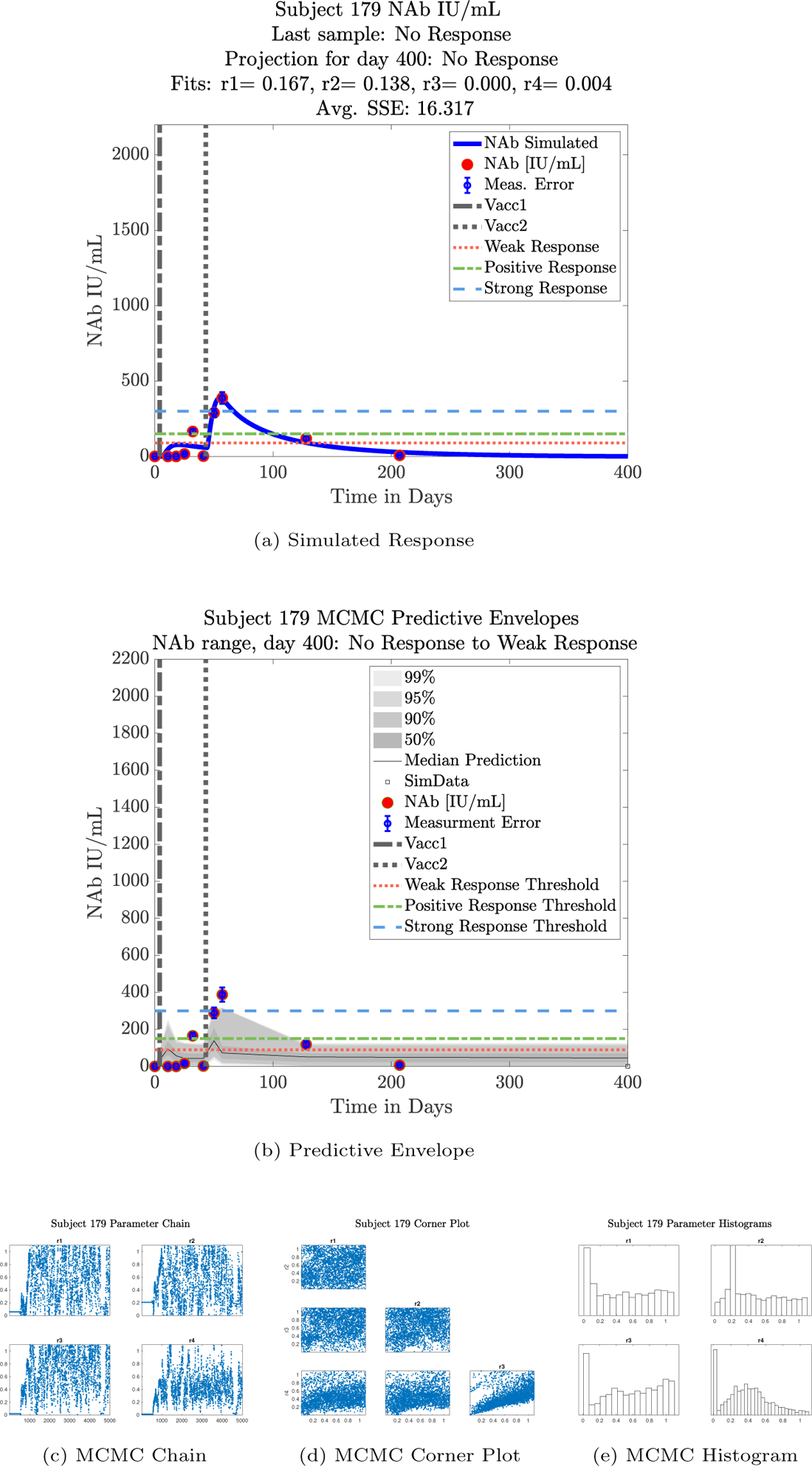
Subject 179

**Figure 46:**
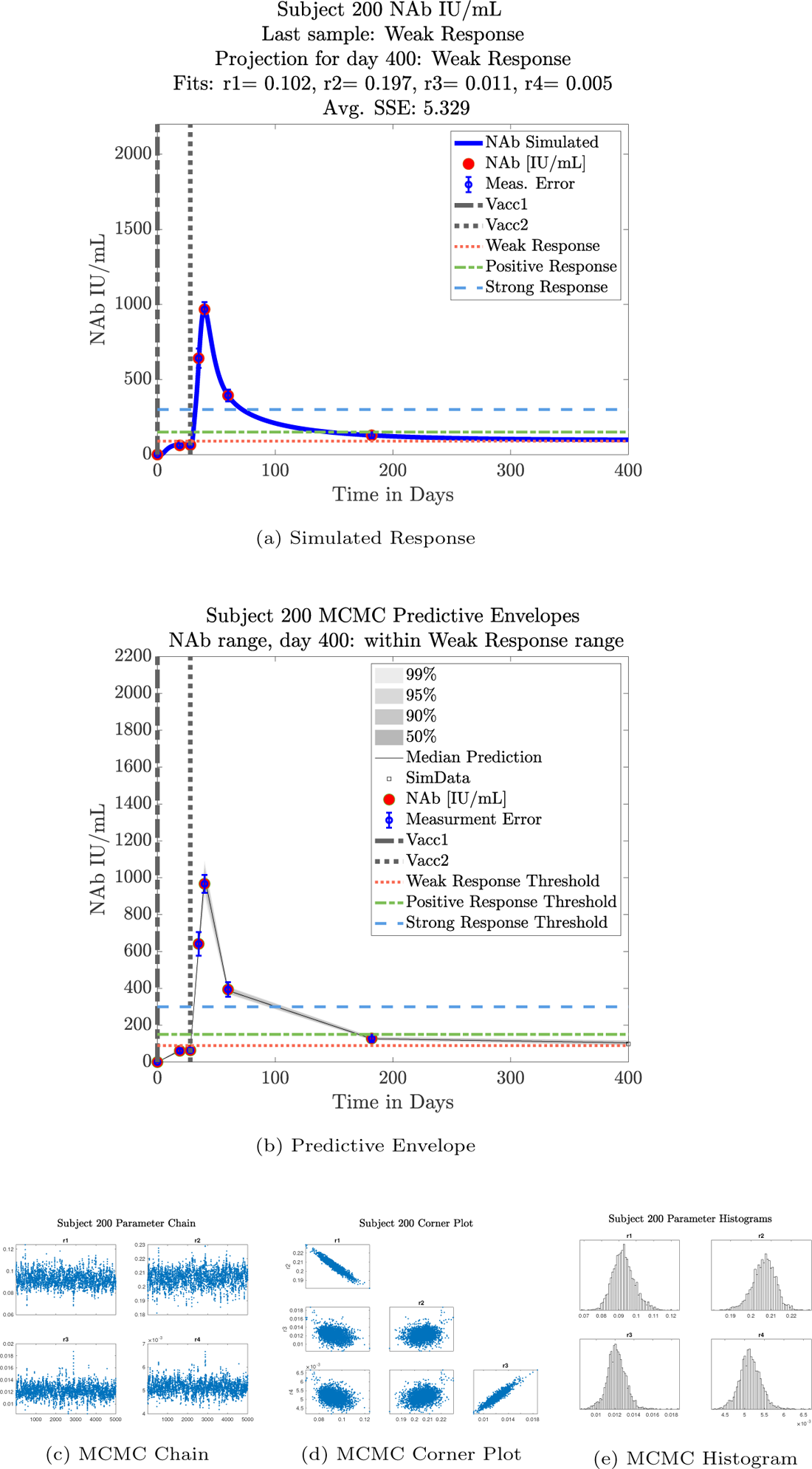
Subject 200

**Figure 47:**
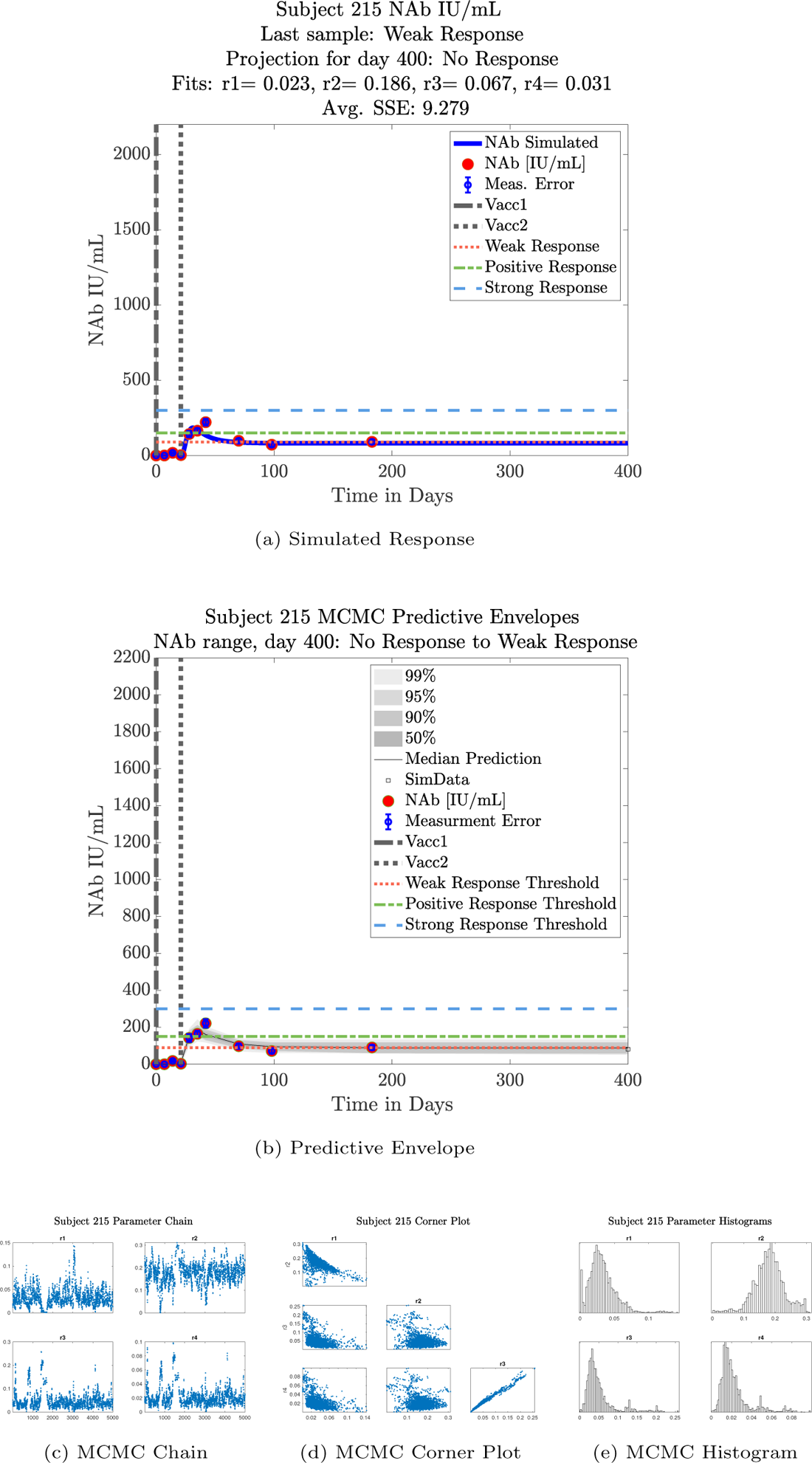
Subject 215

**Figure 48:**
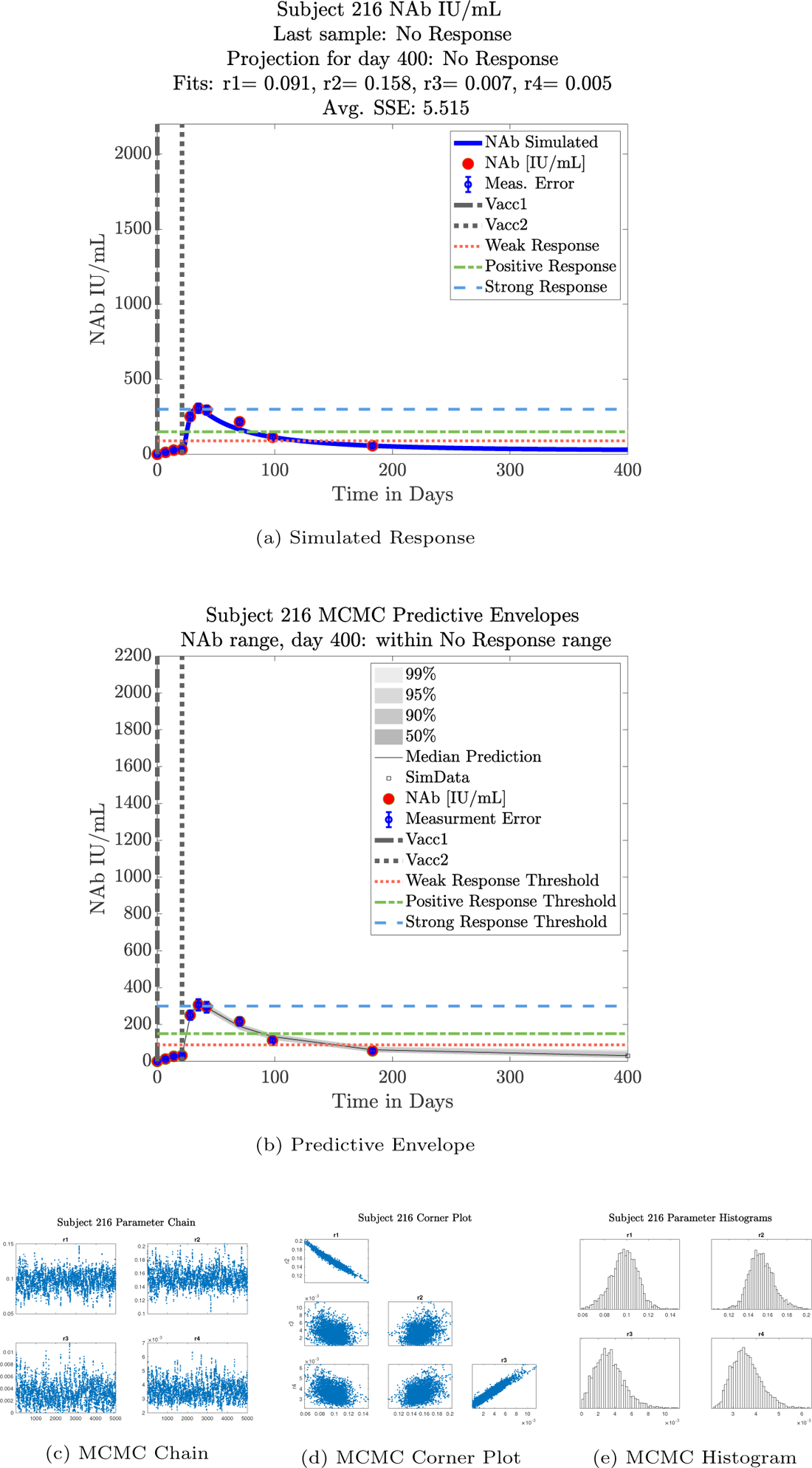
Subject 216

**Figure 49:**
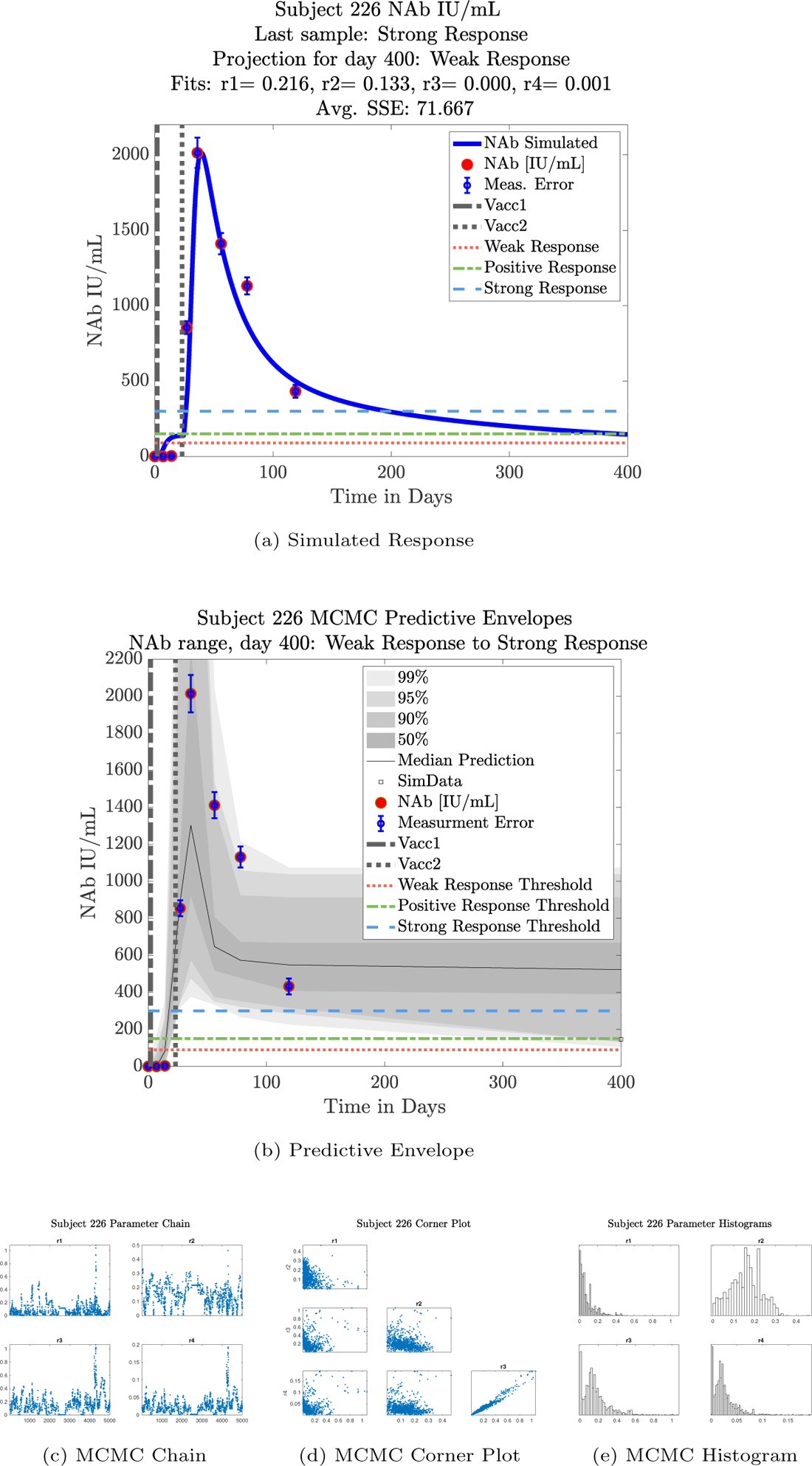
Subject 226

## Supporting information

Article Highlights

---

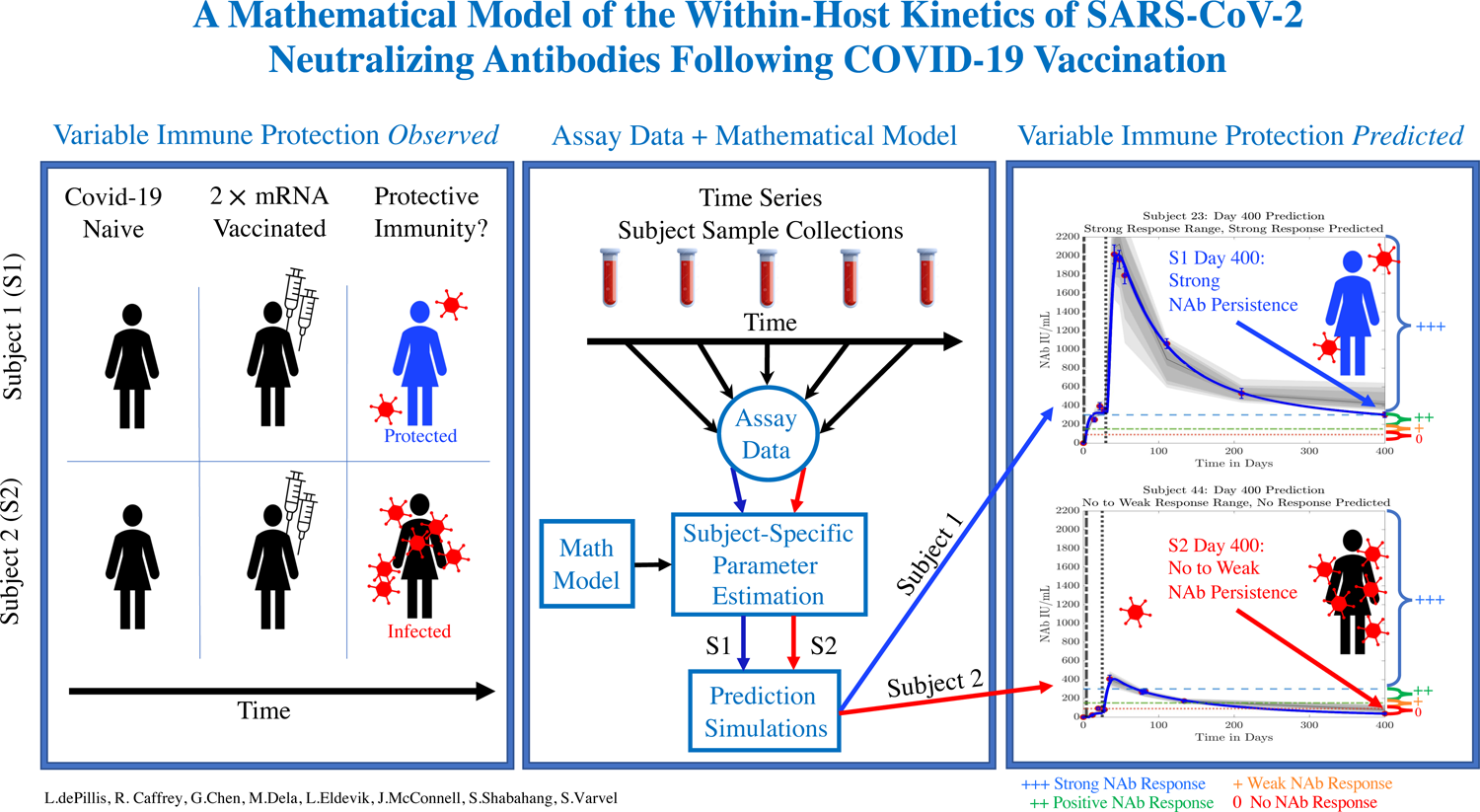

## References

1. D. Khoury, D. Cromer, A. Reynaldi, T. Schlub, A. Wheatley, J. Juno, K. Subbarao, S. Kent, J. Triccas, M. Davenport, Neutralizing antibody levels are highly predictive of immune protection from symptomatic SARS-CoV-2 infection, Nature Medicine 27 (2021) 1205–1211. doi:https://doi.org/10.1038/s41591-021-01377-8.

2. F. Krammer, A correlate of protection for SARS-CoV-2 vaccines is urgently needed., Nature Medicine 27 (2021) 1147–1148. doi:https://doi-org.ccl.idm.oclc.org/10.1038/s41591-021-01432-4.

3. A. Addetia, K. H. D. Crawford, A. Dingens, H. Zhu, P. Roychoudhury, M.-L. H. K. R. Jerome, J. D. Bloom, A. L. Greninger, Neutralizing Antibodies Correlate with Protection from SARS-CoV-2 in Humans during a Fishery Vessel Outbreak with a High Attack Rate, Journal of Clinical Microbiology 58 (11) (2020) e02107–20. doi:https://doi.org/10.1128/JCM.02107-20.

4. S. Dispinseri, M. Secchi2, M. F. Pirillo, M. Tolazzi, M. Borghi, C. Brigatti, M. L. D. Angelis, M. Baratella, E. Bazzigaluppi, G. Venturi, F. Sironi, A. Canitano, I. Marzinotto, C. Tresoldi, F. Ciceri, L. Piemonti, A. C. Donatella Negri and, V. Lampasona, G. Scarlatti, Neutralizing antibody responses to SARS-CoV-2 in symptomatic COVID-19 is persistent and critical for survival, Nature Communications 12 (2670) (2021) e1–12. doi:https://doi.org/10.1038/s41467-021-22958-8.

5. C. D. Murin, I. A. Wilson, A. B. Ward, Antibody responses to viral infections: a structural perspective across three different enveloped viruses, Nature Microbiology 4 (5) (2019) 734–747. doi:https://doi.org/10.1038/s41564-019-0392-y.

6. R. Vandergaast, T. Carey, S. Reiter, C. Lathrum, P. Lech, C. Gnanadurai, M. Haselton, J. Buehler, R. Narjari, L. Schnebeck, A. Roesler, K. Sevola, L. Suksanpaisan, A. Bexon, S. Naik, B. Brunton, S. C. Weaver, G. Rafael, S. Tran, A. Baum, C. A. Kyratsous, K. W. Peng, S. J. Russell, IMMUNO-COV v2.0: Development and Validation of a High-Throughput Clinical Assay for Measuring SARS-CoV-2-Neutralizing Antibody Titers, mSphere 6 (3). doi:https://doi.org/10.1128/mSphere.00170-21.

7. M. Diagne, H. Rwezaura, S. Tchoumi, J. Tchuenche, A Mathematical Model of COVID-19 with Vaccination and Treatment, Computational and Mathematical Methods in Medicine 2021 (2021) 1–16. doi:https://doi.org/10.1155/2021/1250129.

8. D. A. Swan, A. Goyal, C. Bracis, M. Moore, E. Krantz, E. Brown, F. Cardozo-Ojeda, D. B. Reeves, F. Gao, P. B. Gilbert, L. Corey, M. S. Cohen, H. Janes, D. Dimitrov, J. T. Schiffer, Mathematical modeling of vaccines that prevent sars-cov-2 transmission, Viruses 13 (10). doi: 10.3390/v13101921. URL https://www.mdpi.com/1999-4915/13/10/1921

9. C. Li, J. Xu, J. Liu, Y. Zhou, The within-host viral kinetics of SARS-CoV-2, Mathematical Biosciences and Engineering 17 (4) (2020) 2853–2861. doi:10.3934/mbe.2020159.

10. B. J. Nath, K. Dehingia, V. N. Mishra, Y.-M. Chu, H. K. Sarmah, Mathematical analysis of a within-host model of SARS-CoV-2, Advances in Difference Equations 2021 (113) (2021) e1–11. doi:https://doi.org/10.1186/s13662-021-03276-1.

11. F. Pan, T. Ye, P. Sun, S. Gui, B. Liang, L. Li, D. Zheng, J. Wang, R. L. Hesketh, L. Yang, C. Zheng, Time Course of Lung Changes at Chest CT during Recovery from Coronavirus Disease 2019 (COVID-19), Radiology 295 (3) (2020) 715–721. doi: https://doi.org/10.1148/radiol.2020200370.

12. M.-D. Oh, W. B. Park, P. G. Choe, S.-J. Choi, J.-I. Kim, J. Chae, S. S. Park, E.-C. Kim, H. S. Oh, E. J. Kim, E. Y. Nam, S. H. Na, D. K. Kim, S.-M. Lee, K.-H. Song, J. H. Bang, E. S. Kim, H. B. Kim, S. W. Park, N. J. Kim, Viral Load Kinetics of MERS Coronavirus Infection, New England Journal of Medicine 375 (13) (2016) 1303–1305. doi:https://doi.org/10.1056/nejmc1511695.

13. S. Farhang-Sardroodi, C. S. Korosec, S. Gholami, M. Craig, I. R. Moyles, M. S. Ghaemi, H. K. Ooi, J. M. Heffernan, Analysis of host immunological response of adenovirus-based covid-19 vaccines, Vaccines 9 (8). doi:10.3390/vaccines9080861. URL https://www.mdpi.com/2076-393X/9/8/861

14. M. N. Ramasamy, A. M. Minassian, K. J. Ewer, A. L. Flaxman, P. M. Folegatti, D. R. Owens, et al., Safety and immunogenicity of ChAdOx1 nCoV-19 vaccine administered in a prime-boost regimen in young and old adults (COV002): a single-blind, randomised, controlled, phase 2/3 trial, The Lancet 396 (2020) 1979–1993.

15. F. Wu, A. Wang, M. Liu, Q. Wang, J. Chen, S. Xia, Y. Ling, Y. Zhang, J. Xun, L. Lu, S. Jiang, H. Lu, Y. Wen, J. Huang, Neutralizing antibody responses to SARS-CoV-2 in a COVID-19 recovered patient cohort and their implications, medRxivarXiv:https://www.medrxiv.org/ content/early/2020/04/20/2020.03.30.20047365.full.pdf, doi:10.1101/2020.03.30.20047365. URL https://www.medrxiv.org/content/early/2020/04/20/2020.03.30.20047365

16. K. S. Kim, K. Ejima, S. Iwanami, asuhisa Fujita, H. Ohash, Y. Koizum, Y. Asa, S. Nakaoka, K. Watashi, K. Aihara, R. N. Thompson, R. Ke, A. S. Perelson, S. Iwami, A quantitative model used to compare within-host SARS-CoV-2, MERS-CoV, and SARS-CoV dynamics provides insights into the pathogenesis and treatment of SARS-CoV-2, PLoS Biology 19 (3) (2021) e3001128. doi:https://doi.org/10.1371/journal.pbio.3001128.

17. M. Sadria, A. T. Layton, Modeling within-Host SARS-CoV-2 Infection Dynamics and Potential Treatments, Viruses 13 (1141) (2021) e1–15. doi: https://doi.org/10.3390/v13061141.

18. K. Ejima, K. S. Kim, Y. Ito, S. Iwanami, H. Ohashi, Y. Koizumi, K. Watashi, A. I. Bento, K. Aihara, S. Iwami, Inferring Timing of Infection Using Within-host SARS-CoV-2 Infection Dynamics Model: Are “Imported Cases” Truly Imported?, medRxivarXiv:https://www.medrxiv.org/content/early/2020/03/31/2020.03.30.20040519.full.pdf, doi:10.1101/2020.03.30.20040519. URL https://www.medrxiv.org/content/early/2020/03/31/2020.03.30.20040519

19. E. A. Hernandez-Vargas, J. X. Velasco-Hernandez, In-host Modelling of COVID-19 in Humans, medRxivarXiv:https://www.medrxiv.org/content/early/2020/04/15/2020.03.26.20044487.full.pdf, doi:10.1101/2020.03.26.20044487. URL https://www.medrxiv.org/content/early/2020/04/15/2020.03. 26.20044487

20. K. S. Kim, K. Ejima, Y. Ito, S. Iwanami, H. Ohashi, Y. Koizumi, Y. Asai, S. Nakaoka, K. Watashi, R. N. Thompson, S. Iwami, Modelling SARS-CoV-2 Dynamics: Implications for Therapy, medRxiv doi:10.1101/2020.03.23.20040493.

21. S. Sahoo, K. Hari, S. Jhunjhunwala, M. K. Jolly, Mechanistic modeling of the SARS-CoV-2 and immune system interplay unravels design principles for diverse clinicopathological outcomes, bioRxivarXiv:https://www.biorxiv.org/content/early/2020/05/16/2020.05.16.097238.full.pdf, doi:10.1101/2020.05.16.097238. URL https://www.biorxiv.org/content/early/2020/05/16/2020.05.16.097238

22. B. Chatterjee, H. S. Sandhu, N. M. Dixit, The relative strength and timing of innate immune and CD8 T-cell responses underlie the heterogeneous outcomes of SARS-CoV-2 infection, medRxivarXiv:https://www.medrxiv.org/content/early/2021/06/21/2021.06.15.21258935.full.pdf, doi:10.1101/2021.06.15.21258935. URL https://www.medrxiv.org/content/early/2021/06/21/2021.06.15.21258935

23. A. P. Tran, M. Ali Al-Radhawi, I. Kareva, J. Wu, D. J. Waxman, E. D. Sontag, Delicate Balances in Cancer Chemotherapy: Modeling Immune Recruitment and Emergence of Systemic Drug Resistance, Frontiers in Immunology 11. doi:10.3389/fimmu.2020.01376.

24. Matlab optimization toolbox, the MathWorks Inc., Natick, MA, USA (R2021a).

25. M. Laine, MCMCstat for MATLAB. URL https://mjlaine.github.io/mcmcstat/

26. H. Haario, E. Saksman, J. Tamminen, An adaptive Metropolis algorithm, Bernoulli 7 (2001) 223–242. doi:http://dx.doi.org/10.2307/3318737.

27. H. Haario, M. Laine, A. Mira, E. Saksman, DRAM: Efficient adaptive MCMC, Statistics and Computing 16 (2016) 339–354. doi:http://dx.doi.org/10.2307/3318737.

28. MATLAB, version 9.10.0.1739362 (R2021a), The MathWorks Inc., Natick, Massachusetts, 2021.

29. S. Fischinger, C. M. Boudreau, A. L. Butler, H. Streeck, G. Alter, Sex differences in vaccine-induced humoral immunity, Seminars in Immunopathology 41 (2019) 239–249. doi:https://doi.org/10.1007/s00281-018-0726-5.

