## Supplementary material for "A Mathematical Model of the Within-Host Kinetics of SARS-CoV-2 Neutralizing Antibodies Following COVID-19 Vaccination": Article Highlights

1. New ODE model predicts within-host strength of antibody response to mRNA vaccine
2. ODE model tracks subject-specific persistence of SARS-CoV-2 NAbs over time
3. Subject-specific predictive envelopes generated via MCMC application to ODE system
4. Time series data from twice-mRNA-vaccinated Covid19-naive human subjects captured
5. Model reflects range of subject-specific variation in protective immunity
